# A Spin-Glass Metabolic Hamiltonian optimized by Quantum Annealing Reveals Configurational Regimes of Cancer Metabolism

**DOI:** 10.64898/2026.04.05.715441

**Authors:** Ji-Yong Sung, Kyunghyun Baek, Inbeom Park, Jeongho Bang, Jae-Ho Cheong

## Abstract

Understanding why specific metabolic states become stable in cancer has remained a fundamental challenge, as current pathway-centric frameworks lack a unifying physical principle governing global metabolic organization. We introduce the Metabolic Spin-Glass (MSG) model, which represents cellular metabolism using a transcriptome-informed effective Hamiltonian that integrates reference reaction free energies, cofactor-mediated network couplings, and patient-specific transcriptomic fields within a frustrated many-body optimization framework. The Hamiltonian is formulated as a binary optimization problem and solved using hybrid quantum annealing. Embedding gastric cancer transcriptomes (n = 497) reveals that gene expression actively reshapes the effective metabolic optimization landscape in a subtype-associated manner, rather than acting through isolated pathway perturbations. A model-derived order parameter is significantly associated with overall and disease-free survival after adjustment for clinical covariates and the dominant transcriptomic principal components, indicating that it captures prognostic information that is not fully redundant with transcriptomic structure, although it remains substantially correlated with it. This work establishes a transcriptome-informed spin-glass optimization framework for patient-specific characterization and stratification of cancer metabolic organization.

## Introduction

Cancer cells undergo extensive metabolic reprogramming that sustains uncontrolled proliferation, stress resilience, and phenotypic plasticity [1] [2] [3]. For nearly a century, the dominant conceptual framework for understanding this reprogramming has been the Warburg effect [4], the preferential engagement of aerobic glycolysis even in the presence of functionally intact mitochondrial respiration [5] and its descendants in pathway-centric cancer metabolism research. Advances in genomics, transcriptomics, and metabolomics have catalogued an expanding repertoire of altered metabolic reactions in cancers [6], yet these descriptions remain fundamentally explanatory rather than predictive: they identify which reactions change, but cannot explain why particular global metabolic organizations emerge and persist [7] [8] across diverse genetic and environmental contexts.

A central conceptual gap in current models is the absence of a unifying physical principle that connects molecular-level reaction thermodynamics to the system-level organization of metabolic states [9]. Cellular metabolism does not operate as a collection of independently regulated pathways. Instead, reactions are embedded in a densely coupled network in which shared cofactors such as ATP, NADH, NADPH, inorganic phosphate, and protons [10] impose global energetic constraints that link the feasibility of one reaction to the activation of many others [11] [12]. Reactions that draw on the same cofactor can compete through shared cofactor demands, whereas reactions linked through complementary cofactor production and consumption can exhibit cooperative coupling within the network. This network of cooperative and antagonistic couplings generates a system in which the metabolic state of cell reflects a collective, emergent organization rather than a simple superposition of independent pathway activities [13][14][15]. This phenomenology is formally analogous to frustrated many-body systems in condensed-matter physics, most notably spin glasses, where competing pairwise interactions generate rugged optimization landscapes populated by multiple locally stable configurations, slow relaxation dynamics, and sensitivity to small perturbations. In spin glasses, the ground state cannot be inferred from individual pairwise interactions alone; it emerges from their collective competition under global thermodynamic constraints. We propose that cancer metabolism occupies an analogous regime: the favored reaction configuration within the model is not determined by individual enzyme activities but by the collective interplay of reaction free energies, cofactor-mediated coupling, and transcriptome-induced model-derived regulatory biases distributed across the entire metabolic network.

Recent advances in biochemical thermodynamics have made this analogy computationally tractable. Standardized transformed Gibbs free energies (ΔG′) are now available for thousands of metabolic reactions, enabling reaction-level thermodynamic characterization at genome scale [27]. Simultaneously, growing evidence demonstrates that cofactor competition and cooperation introduces emergent constraints that cannot be captured by single-pathway analyses, while high throughput transcriptomics provides patient-resolved quantification of enzyme expression with which to parameterize network-level model-derived energetic biases.

Collectively, these developments create the conditions for a transcriptome-informed, patient-specific model of cancer metabolism grounded in reference reaction energetics.

Importantly, the framework developed here is not intended as a first-principles reconstruction of cellular thermodynamics. Rather, it defines a transcriptome-informed effective statistical-mechanical model in which biochemical energetic priors and transcriptomic constraints are integrated into a unified energy-like objective function. Accordingly, the resulting Hamiltonian values, optimal configurations, and collective variables are interpreted as model-dependent descriptors of metabolic organization rather than direct measurements of cellular Gibbs free energy or microscopic equilibrium states.

Building on these foundations, we introduce a transcriptome-informed effective metabolic spin-glass Hamiltonian that represents cellular metabolism as a frustrated many-body optimization problem defined over classical binary reaction variables.

Each metabolic reaction is represented as a binary activation variable, and the total Hamiltonian value of the system is governed by three additive terms: an intrinsic thermodynamic term encoding the standard transformed Gibbs free energy (ΔG′) of each reaction, a cofactor interaction term, the B matrix, encoding model-defined cooperative and competitive couplings between reactions sharing key redox and energy cofactors, and a transcriptomic field term that modulates the model-defined participation preference each reaction according to patient-specific enzyme abundance derived from gene expression data.

This construction defines a high-dimensional effective optimization landscape over the metabolic configurations represented within the model. Local and global minima correspond to favored reaction configurations within the model under the combined assumptions of reference reaction energetics, cofactor-mediated coupling, and transcriptome-derived regulatory bias.

The resulting Hamiltonian has the mathematical structure of a quadratic unconstrained binary optimization (QUBO) problem which is the canonical formulation of classical spin-glass optimization. Identification of the optimal configuration was performed using D-Wave hybrid quantum annealing (HQA), which integrates quantum annealing with classical optimization routines to efficiently navigate the combinatorial solution space. For the present problem of 235 metabolic reactions, HQA solutions were confirmed to be identical to those obtained by classical integer programming solver, validating the optimality of the identified configurations. This QUBO formulation naturally positions the MSG framework for scaling to genome-wide reaction spaces as quantum hardware continues to advance.

Applying this framework to a gastric cancer cohort of 497 patients across five established molecular subtypes, we demonstrate that transcriptomic reprogramming systematically reshapes the metabolic optimization landscape, generating transcriptome-dependent configurational regimes with distinct solution cluster geometries; the continuous order parameter, but not discrete subtype membership, is further shown to be independently associated with clinical outcome. These results establish the MSG model as a physically motivated, transcriptome-informed framework for cancer metabolic stratification, and demonstrate that spin-glass statistical mechanics, combined with quantum annealing-based optimization, provides a powerful and scalable approach to understanding metabolic heterogeneity in cancer.

## Results

### Transcriptome-associated deformation of the effective metabolic optimization landscape defines subtype-specific metabolic regimes

To investigate how transcriptomic reprogramming reshapes cancer metabolism at a system level [16] [17] [18], we formulated a transcriptome-informed Hamiltonian that integrates reaction free energies, cofactor-mediated interactions, and patient-specific gene expression. Embedding gastric cancer transcriptomes [19] into this framework revealed that gene expression patterns actively deform the underlying metabolic optimization landscape, generating subtype-specific environments (**Figure 1A**).

**Figure 1.**
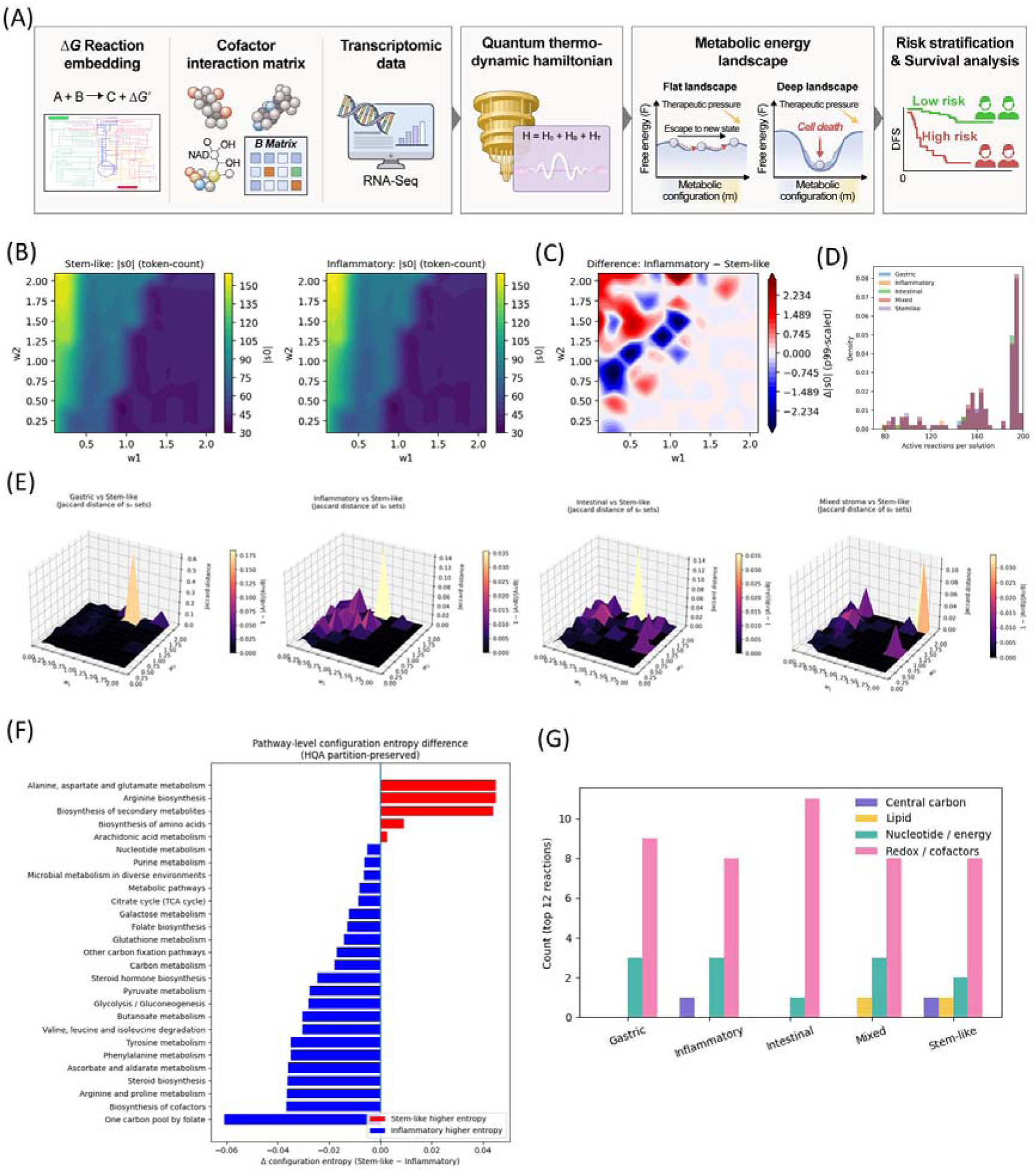
Transcriptome-driven reprogramming of the metabolic optimization landscape in gastric cancer. (A) Schematic of the computational framework. Transcriptome data are mapped onto a thermodynamic metabolic network and integrated into an effective Hamiltonian, minimized by hybrid quantum annealing, to reconstruct patient-specific optimization landscapes, enabling identification of low-objective configurations and subtype-specific configurational regime structure. (B) Reaction-level optimization landscapes across molecular subtypes. Heatmaps show the density of low-energy configurations as a function of Hamiltonian parameters. Inflammatory tumors display a sharply localized density region, whereas stem-like tumors exhibit broader configurational diversity. (C) Difference map of reaction-set size (|s_0_|) between Inflammatory and Stem-like tumors across the (w_1_, w_2_) parameter grid (red, Inflammatory > Stem-like; blue, Stem-like > Inflammatory), corresponding to panel (B). (D) Distribution of quantum-optimized solution ensembles. Stem-like tumors are enriched in extreme solution sizes, while inflammatory and mixed subtypes show broader intermediate distributions. (E) Jaccard distances between the active reaction sets of gastric, inflammatory, intestinal, and mixed subtypes and the stem-like reference configuration, showing subtype-dependent differences in optimized reaction-participation patterns. This stem-like-referenced comparison is shown for completeness; because it uses the stem-like solution as the point of comparison, the apparent separation reflects the choice of reference rather than an intrinsic thermodynamic distinction. A reference-independent analysis in the neutral (m, F_norm) space found no significant separation between subtypes (see Discussion and Supplementary Figure 4). (F) Pathway-level configuration diversity differences (Stem-like vs Inflammatory), highlighting subtype-specific metabolic asymmetry. (G) Functional categorization of entropy-contributing reactions. Redox- and cofactor-associated reactions dominate the top-12 contributing reactions across all five subtypes. Nucleotide/energy-related reactions contribute across all subtypes, whereas central-carbon reactions appear only among the top-12 reactions for stem-like tumors, and lipid-related reactions appear only for stem-like and mixed-stroma tumors.

Contour density maps of active reaction token counts (|S□|) revealed distinct configuration geometries between stem-like and inflammatory tumors (**Figure 1B**). Stem-like tumors [20] [21] [22] [23] [24] [25] occupied a concentrated region of the w□–w□ parameter space, whereas inflammatory tumors exhibited a broader distribution extending toward higher-w□ regimes. Direct subtraction of configuration densities (Inflammatory − Stem-like) further demonstrated localized zones of strong configurational divergence (**Figure 1C**), indicating that these subtypes occupy structurally distinct sectors of model-defined reaction-configuration space rather than representing scaled variations of a shared metabolic backbone. Consistently, the distribution of active reactions per solution showed that stem-like tumors occupy a broader configurational regime compared to other subtypes (**Figure 1D**), suggesting reduced metabolic degeneracy.

To incorporate redox and cofactor-mediated coupling, we constructed a cofactor interaction matrix (B-matrix) and embedded it into a Hamiltonian of the form H = H□ + H_B_ + H_T_ (**Figure 1A**). Here, H□ encodes intrinsic reaction free energy contributions, H_B_ represents cofactor interaction terms, and H_T_ reflects transcriptome-weighted perturbations.

Jaccard distance analysis of active reaction sets revealed subtype-dependent differences in reaction configurations relative to the stem-like reference, with gastric, inflammatory, intestinal, and mixed subtypes exhibiting varying degrees of divergence from the stem-like active reaction set (**Figure 1E**). In contrast, inflammatory tumors displayed broader dispersion, consistent with increased Shannon configuration-diversity index.

To test whether this apparent separation reflects a genuine, reference-independent property of stem-like tumors rather than an artifact of using the stem-like solution as the comparison reference, we recomputed the order parameter m and frustration index F_norm directly from the same solution ensembles across the w□ –w□ sweep, without designating any subtype as a reference. In this neutral (m, F_norm) representation, all five subtypes occupied highly overlapping regions of the space (mean pairwise |Δm| = 0.002–0.006 for all subtype pairs), and stem-like tumors were not more distant from the other four subtypes than those subtypes were from one another. We therefore conclude that the apparent separation of stem-like tumors under a stem-like-referenced Jaccard-distance projection reflected the choice of reference rather than an intrinsic thermodynamic distinction between subtypes. (**Supplementary Figure 4**) Projection of Hamiltonian-derived configurations onto a metabolic optimization landscape revealed two characteristic topologies (schematically illustrated in **Figure 1A**): A deep and broad density region was observed for the stem-like state, whereas inflammatory tumors were characterized by a shallower and narrower density region. The stem-like density region exhibited lower statistical-potential minima and smoother curvature, indicative of enhanced high empirical patient-state density and reduced distributional separation toward alternative metabolic configurations. In contrast, the inflammatory density region showed higher effective free energy and sharper local curvature, consistent with increased susceptibility to metabolic state transitions.

Pathway-level configuration diversity differences between stem-like and inflammatory tumors further clarified functional specialization (**Figure 1F**). Importantly, this metric does not merely reflect pathway activity or flux intensity but rather captures the configurational diversity and state-space heterogeneity of metabolic networks within each tumor subtype. In this context, higher Shannon configuration-diversity index indicates a broader ensemble of accessible microstates, suggesting increased plasticity, redox adaptability, and optimization landscape ruggedness at the pathway level. Stem-like tumors showed higher entropy contributions in primary bile acid biosynthesis, purine metabolism, glutathione metabolism, drug metabolism via cytochrome P450, and related redox-associated pathways. Physically, this implies that stem-like tumors occupy a more distributed and degenerate metabolic state-space in redox and detoxification-associated networks, potentially reflecting enhanced robustness against oxidative and xenobiotic stress through multiple energetically accessible configurations. In contrast, inflammatory tumors exhibited higher entropy in the TCA cycle, steroid hormone biosynthesis, amino acid metabolism, and glycolysis/gluconeogenesis. This pattern suggests that inflammatory tumors allocate configurational flexibility toward core bioenergetic and anabolic circuits, indicating a dynamically reconfigurable central carbon metabolism landscape rather than a simple increase in pathway flux. Thus, Shannon configuration-diversity index should be interpreted as a model-derived descriptor of metabolic state multiplicity and network-level degeneracy, rather than a proxy for expression level or activity alone. Functional classification of top reactions confirmed that stem-like tumors are disproportionately enriched in redox/cofactor-associated reactions relative to central carbon, lipid, and nucleotide/energy metabolism classes (**Figure 1G**).

Collectively, these data support a model in which tumor metabolic phenotypes can be conceptualized as structured configurations within an effective optimization landscape shaped by reaction thermodynamics, cofactor-mediated coupling, and transcriptome-weighted perturbations, with density-region geometry and depth varying descriptively across subtypes within each subtype’s own reduced coordinate space rather than reflecting an established stability hierarchy (see Discussion).

Within this landscape, molecular subtypes occupy visually distinct regions of the patient-derived statistical potential surfaces (**Figure 2C**); however, as discussed below, a reference-independent analysis found no significant difference in the order parameter between stem-like and inflammatory tumors specifically (Mann–Whitney p = 0.69), so this description should not be read as establishing that stem-like tumors occupy a uniquely deep or stable density region relative to other subtypes.

**Figure 2.**
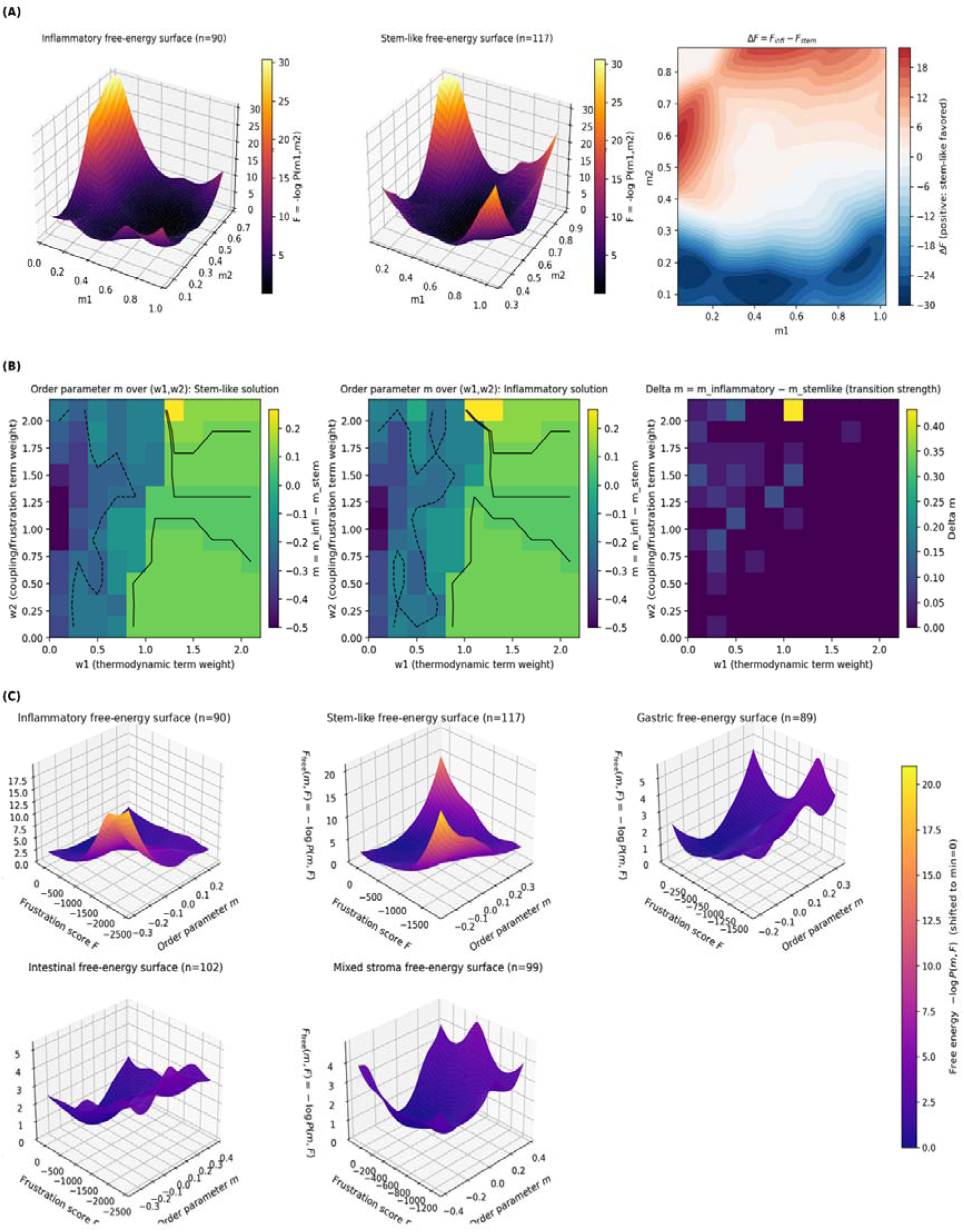
Configurational regime structure of cancer metabolism revealed by collective optimization landscapes. (A) Theoretical reconstruction of subtype-specific statistical-potential landscapes. (left) Inflammatory statistical potential surface ℱ infl (*m*_1_, *m*_2_). (right) Stem-like statistical-potential surface ℱ stem (*m*_1_, *m*_2_). The information-theoretic statistical potential was defined as ℱ = −log*p*(*m*1, *m*2), where *m*1 denotes the first principal component of the standardized patient-level pathway-entropy matrix (see Methods); 2denotes the second; together, they represent data-derived collective coordinates of pathway-level metabolic organization, not independently measured physical order parameters or subtype-overlap metrics. Distinct density-region geometries indicate different stability structures between the two states, consistent with a Kruskal-Wallis test confirming significant subtype separation in this collective-coordinate space (*p* < 0.0001 for both m1 and m2). Statistical-potential difference map Δ ℱ = ℱ_infl_ − ℱ_stem_. Positive values indicate relative stabilization of the stem-like state, whereas negative values indicate dominance of the inflammatory state. (B) Order-parameter behavior in network-coupling and transcriptomic-bias weight space. (Left) Stem-like solution m(w1,w2). (Right) Inflammatory solution m(w1,w2). Here, w1 denotes the relative weight of pairwise network coupling and frustration encoded in the B-matrix, whereas w2 represents the relative weight of the transcriptome-derived local-field term. Contour lines indicate transition boundaries. The difference in the order parameter, Δm=minfl−mstem, highlights regions of maximal transition strength. (C) Patient-derived empirical probability-derived statistical potential landscape reconstructed from molecular subtype distributions. Inflammatory subtype (n = 90), Stem-like subtype (n = 117), Gastric subtype (n = 89), Intestinal subtype (n = 102), Mixed stroma subtype (n = 99). The information-theoretic statistical potential was calculated as ℱ (*m*, *f*_score_) = −log*p* (*m*,_score_), where *m* denotes the order parameter and *F*_score_ represents the frustration score. Each subtype exhibits a characteristic regional topology, reflecting subtype-specific stability structures within a frustrated optimization landscape.

### Empirical Probability-Derived Statistical Potential Landscapes Reveal Distinct Patient-State Distributions in Stem-like and Inflammatory Tumors

To characterize the global organization of tumor-associated patient states, we constructed a two-dimensional empirical probability-derived statistical potential landscape defined over the collective coordinates m and F, based on the empirical probability density of patient-level configurations. The resulting surfaces revealed distinct empirical patient-state density patterns for the stem-like and inflammatory subtypes within the reduced coordinate space (**Figure 2A**).

The stem-like statistical potential surface exhibited a relatively concentrated high-density region centered at positive m, indicating greater empirical occupancy within this region of the reduced patient-state space (**Figure 2A**). In contrast, the inflammatory subtype exhibited a broader and more asymmetric patient-state distribution with greater heterogeneity across the reduced coordinate space (**Figure 2A**). Direct comparison of the two landscapes through the difference map Δ*F* = *F_infl_* - *F_stem_* demonstrates a structured energetic gradient across the (*m*1,*m*2) plane, with a curved transition boundary rather than a linear separatrix (**Figure 2A**), indicating nonlinear coupling between thermodynamic weight and frustration strength.

To further probe the stability structure of these solution clusters, we systematically varied the interaction weight *w*1 (multiplying the B-matrix term *b_ij_*) and the field weight *w*2 (multiplying the local bias term *S_i_*). The Hamiltonian was defined as:

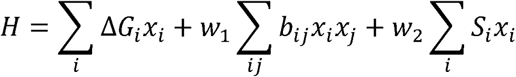

where the coefficient of Δ*G* was fixed to 1. Here, w_1_ controls the relative contribution of pairwise coupling and frustration encoded in the B-matrix, whereas w_2_ regulates the relative contribution of the transcriptome-derived local field S_i_. These parameters are dimensionless model weighting coefficients rather than directly measurable physiological or thermodynamic variables. Accordingly, the w_1_–w_2_ analysis represents a model sensitivity analysis that evaluates how optimized reaction configurations depend on the relative contributions of network coupling and transcriptomic bias. Notably, when w_2_=0, the model is determined solely by the reference reaction-energy and cofactor-interaction terms, without contribution from the transcriptome-derived field, thereby providing a reference configuration for evaluating the influence of transcriptomic weighting. To further probe the stability structure of these solution clusters, we systematically varied the coupling/frustration weight 1 and the local-field weight w_2_ and evaluated the resulting order-parameter values for both solutions (**Figure 2B**). A sharp discontinuity in *m* emerges around *w*_1_=1.0, consistent with a sharp reorganization of the model solution structure in the underlying effective Hamiltonian. Increasing *w*2 expands the solution region of the inflammatory state within this effective model, consistent with increasing weight on the transcriptome-derived field term shifting the relative balance of solution stability toward the inflammatory configuration; we do not interpret this as evidence of a physical ordering field acting on the tumor. The transition strength, quantified as Δ*m* = *m_infl_*− *m_stem_*, exhibits a confined peak within a narrow parameter window (**Figure 2B**), indicating a narrow region of parameter space in which the model’s solution structure is most sensitive to the relative weighting of the network-coupling and transcriptomic-bias terms. Whether this sensitivity reflects a genuine biological transition or is a property of this particular model parameterization has not been independently tested and should be interpreted with appropriate caution. We next reconstructed subtype-specific statistical-potential landscapes to determine whether these theoretical solution clusters manifest in clinical cohorts (**Figure 2C**). Distinct topological features emerge across subtypes. The inflammatory subtype (n=90) displayed a relatively concentrated high-density region within the reduced patient-state space. The stem-enriched subtype exhibited a prominent high-density region, indicating a relatively concentrated empirical distribution of patient states. In contrast, gastric (n = 89) and intestinal (n = 102) subtypes display multi-region structures with increased surface undulation, suggesting coexistence of competing metabolic configurations and elevated frustration. The mixed-stroma subtype (n=99) exhibited a comparatively diffuse patient-state distribution with lower statistical-potential variation. Across subtypes, differences in surface geometry reflect variation in the empirical density and distribution of patient states within the reduced coordinate space. Collectively, these results demonstrate that tumor phenotypes occupy structured energetic solution clusters within ashared reduced configuration space, and that frustration-modulated coupling governs both stability and transition susceptibility. Rather than representing isolated discrete states, molecular subtypes appear as distinct regions embedded within acontinuous configuration manifold, suggesting that tumor evolution may proceed via deformation of the underlying landscape rather than abrupt state switching (**Figure 2C**). Importantly, the topology of this landscape, including its curvature, region depth, and degree of frustration, naturally gives rise to emergent configurational regime behavior across tumor subtypes. Together, these results establish that gastric cancer metabolism is organized into subtype-specific configurational regimes, with stem-like tumors corresponding to a highly ordered, low-entropy regime and inflammatory tumors occupying a more disordered, high-entropy regime. Importantly, these regimes emerge from the global structure of the metabolic optimization landscape rather than from predefined pathway-level constraints, reinforcing the interpretation of cancer metabolism as a configurational regime problem. [26]

### A Continuous Model-Derived metabolic Order Parameter Stratifies Cancer Metabolism into Regime-Structured Solution-Density Regions

To determine whether tumor metabolism can be described by acollective model-derived variable, we quantified a patient-level order parameter *m* derived from kernelized, reaction-resolved free-energy contributions (**Figure 3A**). Importantly, the order parameter does not quantify absolute reaction activation levels. Rather, it measures the alignment between the reference-energy-defined reaction sign σ_r_ and the patient-specific activation state z_ir_. Reactions activated in the same direction as their intrinsic free-energy bias contribute positively to m, whereas activation opposing thermodynamic preference contributes negatively. Thus, *m* captures directional alignment with the reference energetic prior rather than mere expression magnitude. Across the cohort, *m* was continuously distributed and spanned both negative and positive regimes without evidence of discrete clustering (**Figure 3A, 3B**), indicating that metabolic organization is not binary but lies along a continuous axis. When stratified by molecular subtype, subtype-associated distributional trends were observed: stem-like tumors tended to be enriched toward higher m, whereas inflammatory and intestinal subtypes tended toward lower values, with gastric and mixed stroma occupying intermediate positions (**Figure 3C**). However, the overall difference in the order parameter across the five molecular subtypes did not reach statistical significance (Kruskal–Wallis test, *p* = 0.170). Similarly, the frustration score did not differ significantly across the five molecular subtypes (Kruskal–Wallis test, *p* = 0.429). These findings indicate that neither the continuous order parameter nor the frustration index cleanly separates the five molecular subtypes, consistent with the partial, rather than complete, correspondence between SC-MSG-derived configurational regimes and conventional transcriptome-based classification described elsewhere in this study.

**Figure 3.**
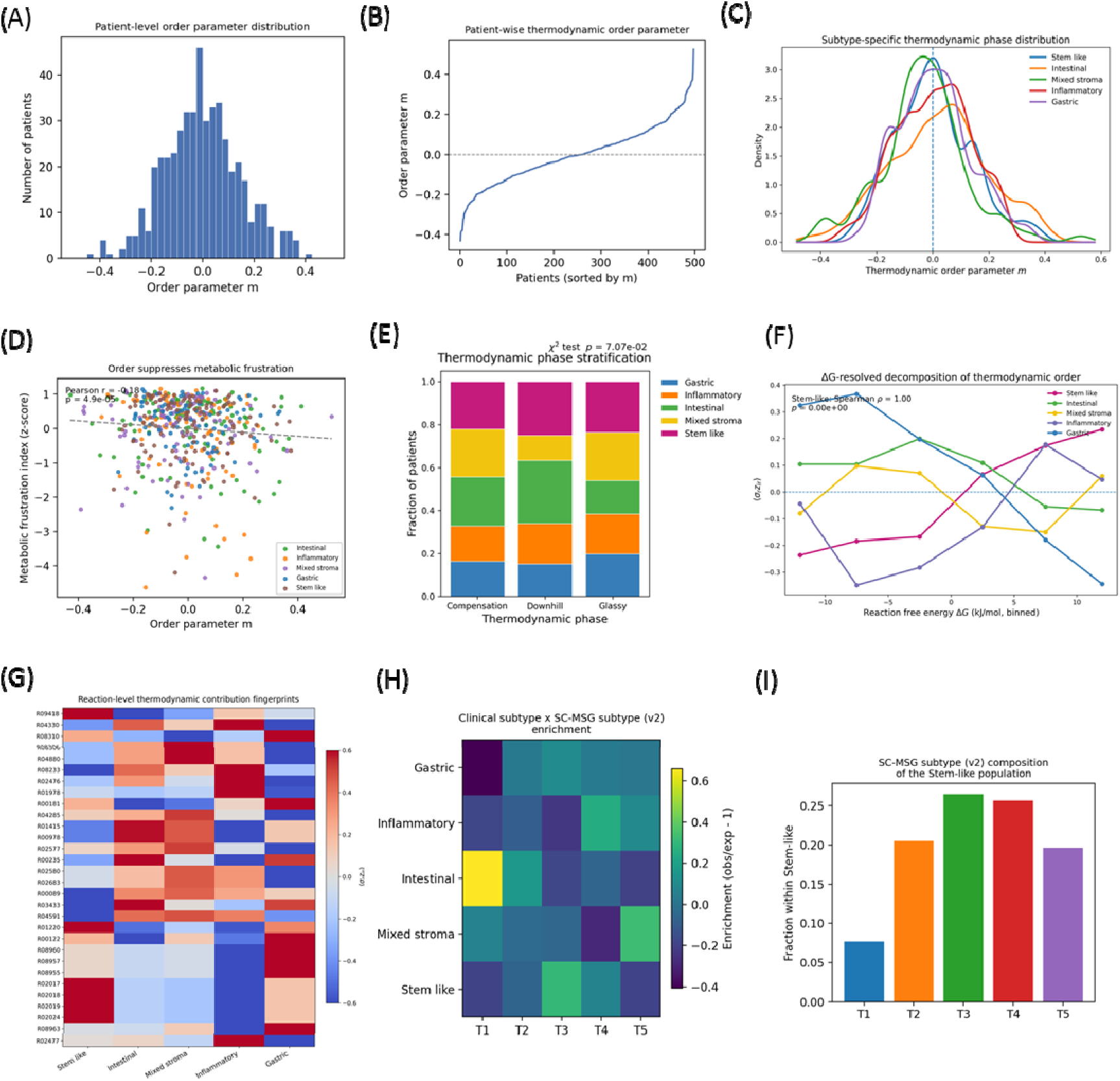
Model-derived metabolic order parameter stratifies configurational regimes and molecular subtypes. (A) Patient-level distribution of the model-derived metabolic order parameter (z-scored per reaction, all 235 reactions; see Methods), showing a near-continuous spectrum across the cohort. (B) Patients ranked by, illustrating a smooth transition from negative to positive order-parameter values without discrete clustering. (C) Subtype-specific density distributions of . Stem-like and mixed stroma subtypes are shifted toward higher order-parameter values, whereas inflammatory and intestinal subtypes exhibit lower values. The dashed line indicates the cohort mean. (D) Inverse correlation between the order parameter and metabolic frustration index (Pearson r = −0.18, *p* = 4.90×10□□), indicating that higher is associated with lower metabolic frustration. (E) Stratification of patients into three configurational regimes (Compensation, Downhill, Glassy). Stacked bar plot shows subtype composition within each regime (χ² test *p* = 7.07×10 □²). (F) ΔG-resolved decomposition of the order parameter. Contributions to *m* are shown as a function of binned reaction free energy, revealing subtype-specific energetic fingerprints. (G) Reaction-level thermodynamic contribution fingerprints across subtypes. Heatmap displays normalized contributions (σ□ z) for representative reactions, highlighting coordinated energetic rewiring in the stem-like state. The corresponding KEGG reaction identifiers and detailed biochemical annotations are provided in Supplementary Data 5. (H) Enrichment heatmap linking clinical molecular subtypes to SC-MSG-derived subtypes (T1–T5), revealing non-random correspondence. (I) Fractional composition of SC-MSG-derived subtypes within the stem-like group, indicating preferential occupancy of high-order states.

We next examined whether an increasing order parameter m reflects reduced metabolic conflict. Indeed, *m* was significantly negatively correlated with a reaction-level frustration index (Pearson r< 0, p≪0.001), demonstrating that higher order corresponds to diminished antagonistic coupling across reactions (**Figure 3D**). This relationship supports a spin glass like interpretation in which frustration is progressively relieved as the network condenses into a coherent configurational regime. Stratification of patients into compensation, downhill and glassy regimes based on global configurational-regime structure revealed marked redistribution of molecular subtypes across regimes (**Figure 3E**). Stem-like tumors were preferentially enriched in the glassy-like regime, consistent with a model-favored yet frustrated metabolic configuration, whereas inflammatory and intestinal tumors were overrepresented in compensation or downhill states.

Decomposition of *m* across reaction free-energy bins further revealed subtype-specific energetic fingerprints (**Figure 3F**). Stem-like tumors exhibited dominant positive contributions from strongly exergonic (negative ΔG) reactions, consistent with coordinated alignment along energetically downhill pathways. In contrast, inflammatory and gastric subtypes displayed greater weight in near-equilibrium or mildly uphill bins. Reaction-level contribution maps confirmed structured, subtype-specific energetic contribution patterns rather than diffuse flux variation (**Figure 3G**).

Detailed biochemical annotations of the reactions displayed in Figure 3G, including KEGG reaction identifiers, common reaction names, reaction equations, reference ΔG values, and pathway annotations, are provided in **Supplementary Data 5**.

To determine whether stem-like and inflammatory tumors occupy distinguishable free-energy density regions independently of any subtype-discriminant projection, we evaluated the free-energy profile F(m) = −log P(m) for each subtype directly from the order parameter m as defined in Methods, without constructing a projection axis from subtype-specific optimized solutions. Stem-like and inflammatory patients showed no significant difference in m (Mann– Whitney p = 0.69), and their free-energy profiles overlapped almost completely, with no discernible double-well structure or energy barrier separating the two subtypes.

Mapping clinical molecular subtypes onto SC-MSG-derived subtypes (T1–T5) revealed structured yet incomplete correspondence (**Figure 3H, 3I**). Several molecular subtypes showed preferential enrichment in specific SC-MSG-derived subtypes-for example, Gastric tumors in T3, Intestinal tumors in T4, Inflammatory tumors in T5, and Mixed stroma tumors in T1. However, the correspondence was not one-to-one, and residual heterogeneity persisted within each clinical category. Notably, the stem-like subtype did not exhibit strong enrichment in any single SC-MSG-derived subtype; instead, it was distributed across multiple SC-MSG-derived subtypes, with T1 accounting for the largest fraction, followed by T5, while T3 and T2 were present at lower levels and T4 was minimally represented (**Figure 3I**). This distribution indicates that the stem-like population spans severalconfigurational states, supporting the notion that configurational stratification captures metabolic structure that only partially overlaps with transcriptomic classification.

Together, these results demonstrate that patient-level model-derived metabolic order parameters provide a principled and quantitative framework for stratifying cancer metabolism. By capturing both continuous variation and discrete configurational structure, this approach bridges molecular subtype classification with a physically grounded landscape model. Specifically, the continuous model-derived metabolic order parameter suppresses metabolic frustration and stratifies tumors into distinct configurational regimes (**Figure 3**). Molecular subtypes also show structured, though incomplete, correspondence between clinical molecular subtypes and the derived configurational regimes and SC-MSG-derived subtypes (**Figure 3E, 3H, 3I**), indicating that this framework captures metabolic structure that only partially overlaps with conventional transcriptome-based classification; this structural distinctness did not, however, translate into improved survival prediction relative to molecular subtypes (H0-D).

### The Continuous Order Parameter, but Not Discrete Subtypes, Is Associated with Survival

To determine whether the model-derived metabolic order parameter captures clinically meaningful tumor heterogeneity, we visualized the multivariable Cox models introduced above as forest plots (**Figure 4A, 4B**).

**Figure 4.**
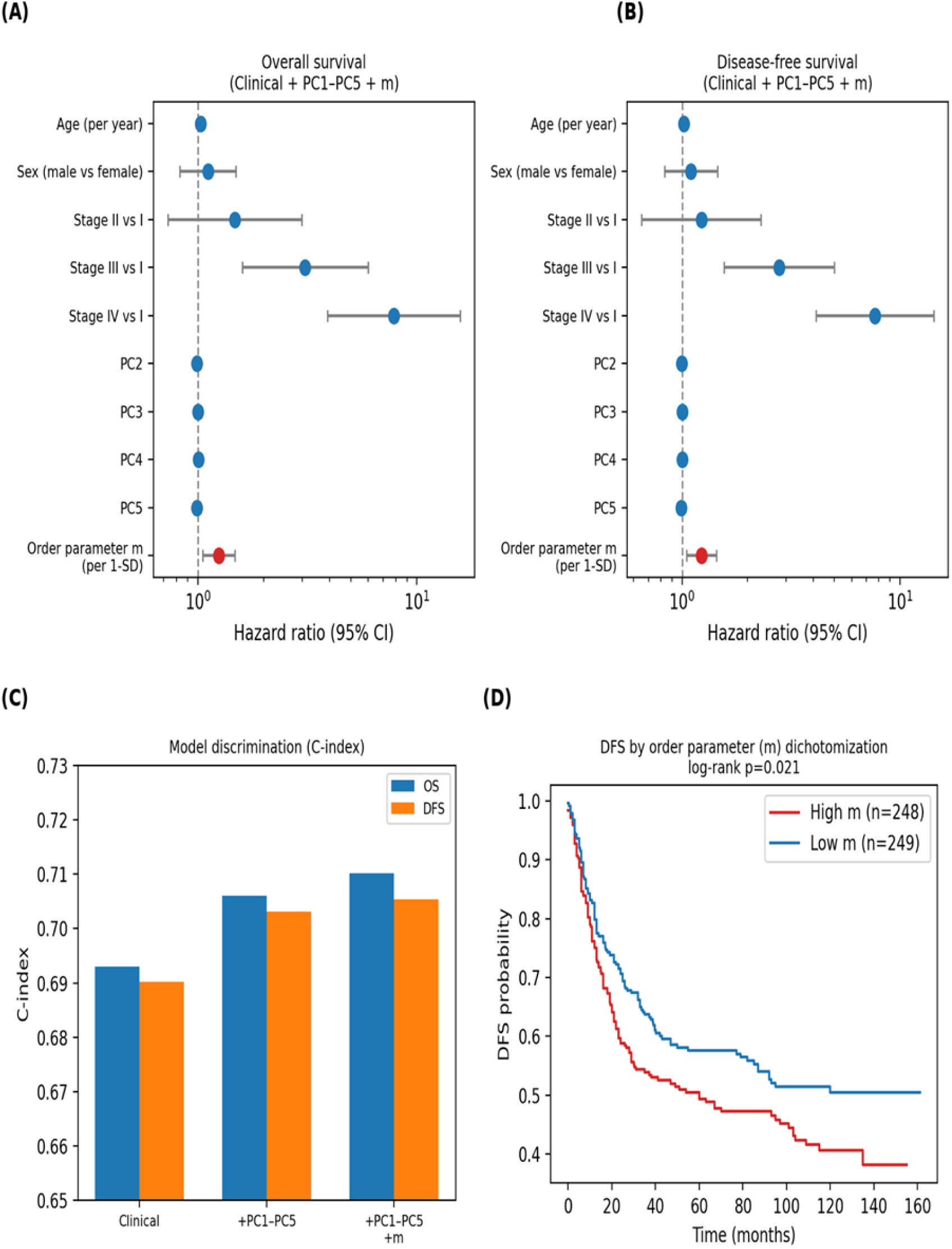
The model-derived metabolic order parameter (m) is associated with survival independently of clinical covariates and the leading transcriptomic principal components. Analyses include all 497 patients (OS: 228 events; DFS: 251 events). (A) Forest plot of the multivariable Cox proportional-hazards model for overall survival (clinical covariates + PC1–PC5 + m). Dots show hazard ratios; bars show 95% confidence intervals. m is expressed per 1-SD increase (HR = 1.25, 95% CI 1.05–1.48). All tests are two-sided. (B) Corresponding forest plot for disease-free survival (m: HR = 1.23, 95% CI 1.05–1.44). (C) Model discrimination (Harrell’s C-index) for Clinical-only, Clinical+PC1–PC5, and Clinical+PC1–PC5+m models, for OS and DFS. Adding m improves discrimination only modestly (ΔC-index = +0.004 for OS and +0.002 for DFS). (D) Kaplan–Meier disease-free survival curves for patients dichotomized at the cohort median of m, shown for visualization only; the continuous, adjusted models in (A) and (B) are the primary analyses. High-m tumors show worse DFS than low-m tumors (unadjusted log-rank *p* = 0.021).

Expressed per 1-SD increase in m, the hazard ratio was 1.25 (95% CI 1.05–1.48) for OS and 1.23 (95% CI 1.05–1.44) for DFS (**Figure 4A, 4B**), and model discrimination improved only modestly with the addition of m (ΔC-index = +0.004 for OS and +0.002 for DFS; **Figure 4C**). Consistent with this continuous association, dichotomizing patients at the cohort median m showed significantly worse DFS for high-m tumors (log-rank *p* = 0.021; **Figure 4D**). As a secondary, exploratory analysis, we also examined discrete SC-MSG-derived subtypes (T1– T5), obtained by clustering patients on (|m|, F_norm) (forcing k = 5 for continuity with the five-subtype scheme used throughout this study, although BIC favored k = 3). Using these assignments, overall survival did not differ significantly across all five subtypes, in either the whole cohort or stem-like tumors alone (log-rank *p* = 0.295 and 0.242, respectively). Among stem-like tumors, the most divergent pair by pairwise testing was T1 vs T3, which showed nominally worse OS and DFS for T1 (log-rank *p* = 0.044 and 0.037), with T1 combining higher order and lower (more cooperative) frustration than T3 by construction, as order and frustration are the variables used to define the T1/T3 clusters; no test statistic is reported for this comparison. However, T1 comprised only 9 stem-like patients, this comparison was one of 10 pairwise tests without correction for multiple comparisons, and the T1-vs-T3 DFS difference was not significant when the same subtypes were compared across the whole cohort (*p* = 0.677). We therefore consider the discrete subtype findings hypothesis-generating rather than confirmatory, and present them, together with the corresponding molecular-subtype composition and (m, F_norm) landscape analyses, in **Supplementary Figure 2** rather than as a main figure.

To determine whether the model-derived metabolic order parameter provided prognostic information beyond the underlying transcriptomic structure, we performed additional benchmark analyses using principal components derived directly from the transcriptomic expression matrix. PC1–PC5 and PC1–PC10 explained 37.9% and 40.0% of the variance in m, respectively, indicating that the order parameter is related to, but not completely determined by, the dominant transcriptomic structure. However, after adjustment for age, sex, pathological stage, and PC1–PC5, m was independently associated with overall survival (per-1-SD HR = 1.25, 95% CI 1.05–1.48, two-sided *p* = 0.011) and disease-free survival (per-1-SD HR = 1.23, 95% CI 1.05–1.44, two-sided *p* = 0.012). Addition of m significantly improved model fit for both overall survival (likelihood-ratio *p* = 0.010) and disease-free survival (likelihood-ratio *p* = 0.011). Sensitivity analyses using PC1–PC10 yielded consistent, and slightly stronger, results (incremental likelihood-ratio *p* = 0.007 for OS and *p* = 0.010 for DFS) (**Supplementary Table 1–6**). We further probed the robustness of this association with four additional sensitivity analyses. Extending adjustment to PC1–PC20 (75.9% of total variance) attenuated the association for m to below conventional significance (OS: per-1-SD HR = 1.54, 95% CI 0.95–2.51, *p* = 0.083; DFS: HR = 1.38, 95% CI 0.87–2.21, *p* = 0.175), indicating that the independence of m’s prognostic association is sensitive to the number of transcriptomic principal components adjusted for. In contrast, adding clinical molecular subtype (stem-like, inflammatory, gastric, intestinal, mixed stroma) as a categorical covariate alongside PC1–PC5 left m significant (OS: HR = 1.40, 95% CI 1.10–1.78, *p* = 0.006; DFS: HR = 1.38, 95% CI 1.10–1.74, *p* = 0.005). A bootstrap 95% CI for the change in C-index attributable to m (Clinical+PC1–PC5 model, OS endpoint, 1,000 resamples) excluded zero (bootstrap mean ΔC-index = 0.010, 95% CI 0.001–0.025); this bootstrap-based mean differs from the +0.004 direct point estimate reported above because it is the average increment across resampled datasets rather than a single estimate on the full cohort, and five-fold cross-validation (OS endpoint) showed a consistent, out-of-sample improvement in both partial log-likelihood (−1.854 vs. −1.839) and C-index (0.684 vs. 0.691) when m was added. A Schoenfeld residual test found no evidence against the proportional-hazards assumption for m specifically (*p* = 0.53), although the assumption was violated for the categorical stage covariates, a common and separate limitation of this cohort unrelated to m. Taken together, m’s incremental prognostic value is statistically detectable, modest in magnitude, and robust to molecular-subtype adjustment and cross-validation, but **n**ot robust to adjustment for a larger number of transcriptomic principal components, underscoring that it should be interpreted as a modest, model-dependent contribution rather than a fully transcriptome-independent signal. Full numerical results for all five sensitivity analyses are provided in **Supplementary Table 10**

Full methodology and code are provided in the accompanying Supplementary Code repository. Thus, m-based stratification was associated with both OS and DFS in univariable analysis, and this association remained statistically significant after adjustment for clinical covariates and the dominant transcriptomic principal components — indicating that, in the present cohort, the order parameter captures prognostic information that is not fully redundant with transcriptomic structure, even though it remains substantially correlated with it (R² = 37.9–40.0%).

Collectively, these findings indicate that the model-derived metabolic order parameter (m) is associated with stem-like tumor heterogeneity and, modestly, with clinical outcome, whereas frustration was associated with tumor heterogeneity but was not independently linked to survival in this analysis. However, the additional transcriptomic PCA-adjusted analyses indicate that the prognostic information captured by m is at least partly shared with the underlying transcriptomic structure.

## Methods

### Construction of the thermodynamically annotated human metabolic reaction network

A thermodynamically annotated and transcriptomically mappable human metabolic reaction network was constructed by integrating curated human biochemical reactions with GPR annotations, patient transcriptomic data, reaction stoichiometries, and available reference thermodynamic parameters.

The reaction network used in the SC-MSG framework was not defined a priori as a manually selected or intentionally reduced metabolic network. Rather, the final reaction set resulted from the intersection of the data requirements necessary for consistent model parameterization. Reactions were retained only when they (i) were annotated as human metabolic reactions, (ii) had available gene–protein–reaction (GPR) information, (iii) contained GPR-associated genes that could be mapped to the patient transcriptomic dataset, and (iv) had an available standard transformed Gibbs free-energy value, 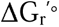, for use as a reference biochemical energetic prior. Reactions lacking one or more of these required data components could not be consistently integrated into the patient-specific SC-MSG framework and were therefore excluded. Thus, the reduction in reaction number arose from cross-database annotation and data-mapping constraints rather than from an a priori selection of specific metabolic pathways or a deliberately simplified metabolic core. Accordingly, the resulting network should be interpreted as a thermodynamically annotated and transcriptomically mappable subset of the human metabolic network that could be consistently parameterized using the available data, rather than as a predefined metabolic network or a complete genome-scale reconstruction.

The complete model inputs are provided as supplementary data. Supplementary Data 1 contains the included human reactions, reaction equations, and reference 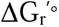 values; Supplementary Data 2 contains the patient-specific transcriptomics-derived reaction matrix; and Supplementary Data 3 contains the corresponding reaction–reaction interaction matrix (B-matrix). Standard transformed Gibbs free-energy changes 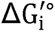 for individual reactions were obtained from the eQuilibrator database and used as reference biochemical energetic priors under standardized conditions. These values characterize the baseline energetics of individual reactions but do not represent patient-specific intracellular thermodynamic driving forces. Under intracellular conditions, the Gibbs free-energy change of reaction i depends on metabolite activities through the reaction quotient Qi, according to

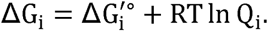

Because patient-matched quantitative metabolomic measurements were unavailable for the present cohort, the concentration-dependent RTln Q_i_ contribution was not explicitly incorporated. Therefore, the 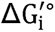 term provides a common reference energetic prior across patients rather than an estimate of the physiological free-energy change of each reaction.

Let x_i_∈{0,1} denote the model-defined participation state of reaction i, where x_i_=1 indicates that reaction i is included in a given optimized reaction-participation configuration and x_i_=0 indicates that it is not included. This binary variable does not represent the magnitude, physical quantization, or presence or absence of metabolic flux. Rather, it constitutes a coarse-grained representation of reaction participation that enables formulation of the collective reaction-selection problem as a quadratic unconstrained binary optimization problem. Because the present SC-MSG objective function does not explicitly impose steady-state mass-balance constraints (Sv=0), reaction-capacity constraints, or metabolic-task requirements, the resulting reaction-participation configurations are not guaranteed to constitute stoichiometrically feasible steady-state flux distributions. Accordingly, the optimized configurations should be interpreted as relative reaction-participation patterns within the effective model rather than as quantitative or biochemically validated steady-state flux distributions. Similarly, x_i_=0 does not imply that reaction i is biochemically inactive or dispensable in the corresponding tumor.

The binary representation retains the reference thermodynamic and interaction terms as energetic priors for ranking reaction configurations but is not assumed to preserve the complete thermodynamic structure of the continuous metabolic flux space. In particular, the present formulation does not explicitly represent flux magnitudes, reaction kinetics, metabolite concentration-dependent driving forces, or mass-balanced steady-state flux distributions. The resulting landscape should therefore be interpreted as an effective configuration landscape over coarse-grained reaction-participation patterns rather than as a complete reconstruction of the continuous thermodynamic state space of cellular metabolism.

To capture network-level interactions associated with shared energetic and redox cofactors, we constructed a cofactor-aware interaction network between reactions. Reactions sharing key energy or redox cofactors, including ATP/ADP, NADH/NAD, NADPH/NADP, inorganic phosphate Pi, and protons, were considered coupled within the effective model. For each pair of reactions i and j, a coupling coefficient B_ij_ was computed based on the stoichiometric consumption or production of these cofactors. Reactions that consumed or produced the same cofactor in the same direction were assigned negative couplings, representing cooperative interactions within the effective objective function, whereas reactions operating in opposite directions with respect to a shared cofactor were assigned positive couplings, representing competitive interactions. The magnitude of each coupling was weighted according to the model-defined biochemical importance of the corresponding cofactor.

### Molecular subtype–specific transcriptomic embedding

To incorporate tumor-specific metabolic constraints, gastric cancer microarray-based transcriptomic data were embedded into a curated metabolic subnetwork consisting of 235 selected reactions, chosen on the basis of data and parameter availability as described in Methods above. Samples were stratified by molecular subtype. For each reaction *i*, the reaction-level expression *E*^-^*i*(*k*) for subtype *k* was defined as the mean gene expression value across all patients within that subtype. The subtype-specific transcriptomic bias was defined as S_i_^(k)^ = −log(1+E^-^_i_^(k)^), such that reactions associated with higher average enzyme expression receive a greater model-defined preference for participation, whereas reactions associated with lower expression receive a lower participation preference. This term represents a transcriptome-derived regulatory bias within the effective objective function and does not imply a direct alteration of the biochemical Gibbs free energy of the reaction. This subtype-dependent bias was incorporated into the third component of the metabolic energy functional,

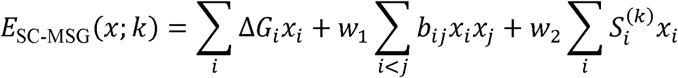

thereby incorporating molecular subtype-specific transcriptional information into the effective configuration landscape.

### Sung-Cheong Metabolic Spin-Glass Model (SC-MSG)

The SC-MSG Hamiltonian is defined as an effective, transcriptome-informed objective function rather than a microscopic physical Hamiltonian derived from the molecular dynamics of the cell. Its purpose is to integrate heterogeneous biochemical and molecular constraints into a common optimization framework. Consequently, the numerical value of the Hamiltonian should not be interpreted as the absolute Gibbs free energy of a cellular metabolic state. Instead, relative Hamiltonian values quantify the preference of reaction configurations under the assumptions and parameterization of the model. Within this effective formulation, metabolic organization was represented using binary reaction-participation variables, x_i_∈{0,1}, where x_i_=1 indicates that reaction i is included in a model-derived configuration. These variables represent coarse-grained reaction participation rather than continuous flux magnitude or intrinsic quantization of metabolic activity. The Hamiltonian introduced above can be written as

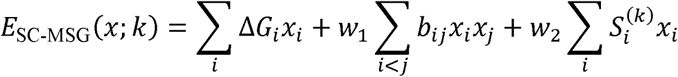

where 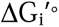 denotes the standard transformed Gibbs free-energy change used as a reference biochemical energetic prior, b_ij_ represents the model-defined cofactor-mediated coupling between reactions i and j, and S_i_ denotes a transcriptome-derived regulatory field representing enzyme expression-associated reaction-participation bias. The first term captures the intrinsic thermodynamic preference of individual reactions, favoring configurations composed of energetically favorable reactions. The second term encodes network-level interactions arising from shared cofactors, such that reactions consuming or producing the same cofactor in the same direction contribute negative couplings (model-defined cooperative), whereas reactions operating in opposing directions contribute positive couplings (model-defined competitive coupling). This interaction term introduces frustration into the system, rendering the metabolic network a many-body thermodynamic system analogous to frustrated spin models. The third term incorporates transcriptomic information as a model-defined regulatory field, assigning a greater participation preference to reactions associated with highly expressed enzymes and a lower participation preference to reactions associated with weakly expressed enzymes. This transcriptome-derived field does not represent a direct modification of reaction Gibbs free energy and does not substitute for the concentration-dependent RTlnQi contribution. The reference reaction-energy term was assigned a unit coefficient, whereas the cofactor-interaction and transcriptomic-field terms were weighted by w_1_ and w_2_, respectively. The parameters w_1_ and w_2_ are dimensionless model weighting parameters rather than directly measurable thermodynamic state variables. Specifically, w_1_ scales the relative contribution of model-defined cofactor-mediated reaction interactions, whereas w_2_ scales the contribution of the transcriptome-derived regulatory field. Variation of these parameters constitutes a model sensitivity analysis that evaluates how the optimized reaction configurations depend on the relative weighting of the objective-function components. Accordingly, changes across the w_1_– w_2_ parameter space are interpreted as reorganizations of the solution structure within the effective model rather than as responses to experimentally controlled thermodynamic variables. Because the three terms represent quantities with different physical and statistical origins, the Hamiltonian is interpreted as a dimensionless effective scoring functional after model-specific scaling and weighting. The optimization therefore ranks metabolic reaction configurations according to their relative compatibility with the incorporated thermodynamic, network, and transcriptomic constraints, rather than estimating an absolute cellular energy. We refer to this Hamiltonian-based formulation of cellular metabolism as the Sung-Cheong Metabolic Spin-Glass Model (SC-MSG). The resulting Hamiltonian has the mathematical structure of a frustrated spin-glass system, whose optimal-configuration search is equivalent to solving a quadratic unconstrained binary optimization (QUBO) problem. Together, this Hamiltonian defines a high-dimensional effective optimization landscape over all possible metabolic configurations. Distinct metabolic phenotypes correspond to local or global minima of this landscape, and transitions between phenotypes arise from changes in the relative dominance of reaction free energies, network couplings, and transcriptomic fields. Within this framework, metabolic organization is represented as an effective frustrated many-body optimization problem, in which favored reaction configurations emerge from the collective interaction of biochemical energetic priors, network couplings, and transcriptomic fields.

### Optimization of lowest-objective metabolic configurations

To identify the lowest-objective metabolic configurations under the assumptions of the SC-MSG model, we minimized H^(p)^(x) for each molecular subtype. The Hamiltonian *H*(*p*)(*x*) is formulated as a quadratic unconstrained binary optimization (QUBO) problem, in which binary variables represent the activation states of metabolic reactions. In general, QUBO problems are considered NP hard; consequently, increasing problem size, corresponding here to the number of reaction activation variables, leads to an exponential growth of the search space. In this work, to enhance computational tractability while retaining biological relevance, we restricted the optimization to 235 reactions selected on the basis of data and parameter availability (see Methods).

In practice, several approaches can be employed to solve QUBO problems, including classical optimization solvers, standard quantum annealing (QA), and the Quantum Approximate Optimization Algorithm (QAOA). To address the computational challenges associated with dense QUBO instances, we adopted a hybrid quantum annealing (HQA) provided by D-Wave, which integrates quantum annealing with classical optimization routines to efficiently solve large-scale problems. Recent large-scale benchmarking studies [30] have demonstrated that HQA systematically outperforms standard quantum annealing, particularly for dense and high-dimensional QUBO problems. Consistent with these findings, in our metabolic optimization setting, HQA consistently yielded lower objective-function values than those obtained using standard QA, indicating more effective identification of the lowest-objective configuration within the SC-MSG model. As a consistency check, we also solved the same QUBO problems using a classical integer programming (IP) solver, which has been shown to outperform other classical solvers [30]. As a result, the minimum objective-function values and reaction activation patterns obtained from IP and HQA were identical, indicating consistent identification of the same optimal solutions by both methods. Although IP and HQA yield identical solutions for the present problem size, previous benchmarking studies [30] suggest that HQA may become increasingly beneficial as the number of reactions grows. The resulting optimized configuration vectors x(p) represent the reaction-participation patterns favored under the reference energetic, network-interaction, and transcriptomic constraints encoded in the SC-MSG objective function. (**Supplementary material 1, 2, 3, 4, 5, 6**)

### Identification of metabolic configurational regimes

In the present study, the term “configurational regime” is used operationally to describe regions of the effective model characterized by qualitatively distinct collective reaction configurations. These regimes should not be interpreted as equilibrium configurational regimes established through a microscopic partition function, thermodynamic-limit analysis, or experimentally measured critical behavior. For each patient, the optimized reaction-participation configuration *x*(*p*) was summarized using pathway-level order parameters. Each order parameter quantified the relative activation of a predefined metabolic module, including glycolysis, the tricarboxylic acid (TCA) cycle, oxidative phosphorylation, and the pentose phosphate pathway, by aggregating the binary activation states of the reactions belonging to each pathway. This coarse-grained representation mapped the microscopic reaction configuration onto macroscopic metabolic observables. Within the Hamiltonian framework, pathway activation serves as a natural order parameter because it captures collective behavior emerging from many interacting reactions. Although individual reaction states may vary, the aggregated pathway-level activation remains stable and reflects the dominant metabolic organization of the optimized configuration, analogous to magnetization in statistical physics. Tumors were classified into distinct configurational regimes based on these order parameters. Regime boundaries were defined as regions in parameter space where the dominant pathway-level order parameter changed sharply as a function of Hamiltonian control parameters (e.g., the relative weights assigned to intrinsic thermodynamic terms and network coupling terms). Mathematically, a regime transition was identified when the global minimum of the Hamiltonian shifted discontinuously between configurations characterized by different dominant pathway activations. Each patient was then assigned to a configurational regime on this basis, and the distribution of molecular subtypes across regimes was examined as described in the Results

### Construction of collective coordinates (m □, m□)

To define a low-dimensional state space capturing dominant pathway-level metabolic variation, we constructed two continuous collective coordinates, denoted as m1 and m2, from the patient-level pathway-entropy matrix. Pathway-level entropy values were standardized across patients, and principal-component analysis (PCA) was applied to the resulting standardized matrix. The scores of the first and second principal components were used as the two collective coordinates and rescaled to the interval [0,1], yielding m1and m2, respectively.

These coordinates represent data-derived low-dimensional axes of pathway-level metabolic variation rather than independently measured physical order parameters. Subtype-specific probability distributions, P(m_1_,m_2_), were estimated in this two-dimensional coordinate space, and the corresponding statistical potential landscapes were calculated as F(m_1_,m_2_) = −ln[P(m1,m2)+ε], where ε is a small positive constant introduced to avoid numerical divergence in sparsely populated regions. The resulting subtype-specific statistical potential landscapes are presented in Figure 2.

### Input data and preprocessing

The final reaction-level expression matrix comprised 235 human metabolic reactions associated with 220 unique genes across 497 patients. Gene-expression values were mapped to their corresponding metabolic reactions using the available gene–reaction associations. Because the relationship between genes and reactions was not necessarily one-to-one, a single reaction could be associated with multiple genes, and a single gene could be mapped to multiple reactions. For reactions associated with multiple genes, the expression values of the mapped genes were averaged to generate a single patient-specific reaction-level expression score. PCA was performed using the patient-by-reaction expression matrix, with the 497 patients as observations and the 235 reaction-level expression scores as feature variables. Thus, the PCA features represented metabolic reactions rather than individual genes. Expression values were log-transformed where appropriate and standardized across patients (zero mean and unit variance) to ensure comparability across genes. Each patient state was represented as a vector 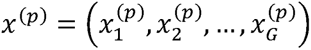 in the G-dimensional gene expression space.

### Identification of dominant collective axes

To extract dominant modes of coordinated variation, we performed principal component analysis (PCA) on the patient-by-gene expression matrix. PCA was applied to the covariance matrix of standardized expression values, yielding orthogonal eigenvectors ranked by explained variance. The first two principal components were retained as candidate collective axes, capturing the largest fraction of inter-patient variability. These axes were subsequently rotated to align with biologically interpretable transcriptional programs.

### Definition of collective coordinates

For each patient p, the collective coordinates were defined as linear projections: 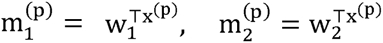 where *w*_1_ and *w*_2_ are the rotated weight vectors corresponding to the stem-like and inflammatory axes, respectively. Importantly, m□ and m□ do not correspond to individual genes, but represent emergent collective degrees of freedom arising from correlated gene expression patterns across the cohort.

### Interpretation as order parameters

Within this framework, m and m function as continuous order parameters describing patient states in a reduced state space. Low-dimensional coordinates enable the construction of effective optimization landscapes and probabilistic state distributions, while retaining information about high-dimensional transcriptional interactions. Thus, m and m represent biologically aligned collective coordinates that map high-dimensional transcriptional states onto a low-dimensional state space suitable for energy-based and landscape-level analysis.

### Construction of Subtype-Specific Probability-Derived Statistical Potential Landscapes

To characterize the global organization of patient transcriptional states across molecular subtypes, we constructed subtype-specific empirical probability-derived statistical potential landscapes in a reduced two-dimensional collective coordinate space defined by the order parameter m and the frustration score *F*, conceptually related to probability-derived entropy potentials used to reconstruct cellular-state landscapes [29]. For each patient *p*, standardized gene expression profiles were projected onto biologically aligned collective axes to obtain coordinates (*m*(*p*),*F*(*p*)), where *m* captures coherent alignment along a predefined biological gradient and *F* quantifies network-level antagonistic tension derived from signed interaction couplings. For each molecular subtype (Inflammatory, Gastric, Intestinal, and Mixed stroma), the empirical joint probability density *p* (*m*,*F*) was estimated using two-dimensional Gaussian kernel density estimation (KDE) applied to the distribution of patient coordinates within that subtype. Bandwidth selection followed Scott’s rule to ensure consistent smoothing across subtypes. The KDE estimator yields a normalized density satisfying

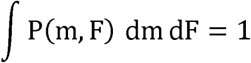

An empirical probability-derived statistical potential was constructed from the estimated patient-state density using the following negative log-density transformation:

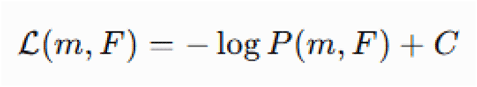

where L(m,F) denotes the empirical probability-derived statistical potential, P(m,F) denotes the normalized empirical density of patient states, ε is a small positive constant introduced to avoid numerical divergence in regions of near-zero density, and C is an arbitrary additive constant. For comparability across subtypes, *C* was chosen such that the global minimum within each subtype was shifted to zero:

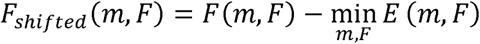

This negative log-density transformation provides an information-theoretic representation of the relative occupancy of observed patient states. It does not assume a Boltzmann distribution, an effective physical temperature, or an equilibrium ensemble generated by the SC-MSG Hamiltonian. Importantly, the resulting surface represents an empirical probability-derived statistical potential inferred from the observed distribution of patient states. It was not derived from the SC-MSG Hamiltonian, a partition function, or an equilibrium Boltzmann ensemble and should therefore not be interpreted as thermodynamic free energy. Accordingly, region depth, width, and barrier-like structures are interpreted as relative statistical features of the observed patient-state distribution and not as directly measurable biochemical free-energy differences or kinetic transition barriers. Local minima of the statistical potential correspond to high-density regions of the observed patient-state distribution. Surface curvature and intervening low-density regions were used as descriptive geometric features of the empirical distribution and should not be interpreted as physical stability measures, kinetic transition barriers, or evidence of metastable configurational states. Statistical-potential surfaces were visualized as continuous three-dimensional surfaces over the (*m, F*_score_) plane. Identical computational procedures, bandwidth selection rules, and normalization steps were applied across subtypes to ensure strict comparability of landscape topology. All density estimation and surface reconstruction were implemented in Python using NumPy, SciPy, and Matplotlib.

### Model-derived metabolic order parameter and frustration index

The term “order parameter” is used here in an effective coarse-grained sense to denote a collective coordinate summarizing high-dimensional reaction-level organization. It is not derived as an equilibrium order parameter from a microscopic partition function and should therefore be interpreted as a model-dependent measure of collective alignment among score-based reaction states. To quantify patient-specific collective metabolic configurations, we mapped reaction-level reference energetic information and transcriptome-derived reaction weights onto an effective spin-like representation. For each metabolic reaction r, we defined a binary model-derived reaction-state variable σ_r_∈{−1,+1}, based on the sign of the transcriptomics-weighted reference energetic score E_r_^TW^

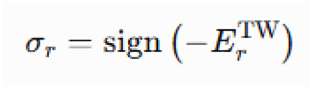

where E_r_^TW^ denotes the transcriptomics-weighted reference energetic score for reaction r, calculated by combining the standard transformed Gibbs free-energy change, 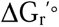, used as a reference biochemical energetic prior, with the patient-specific gene expression-derived reaction weight. The transcriptomic weight represents enzyme expression-associated reaction-participation potential and does not modify the biochemical Gibbs free energy of the reaction. Therefore, E_r_^TW^is a model-derived reaction score rather than an estimate of the physiological Gibbs free-energy change, thermodynamic driving force, or reaction directionality. Because patient-specific intracellular metabolite activities were unavailable, the concentration-dependent RTlnQ_r_ contribution was not included. Accordingly, the physiological reaction Gibbs free-energy change,:

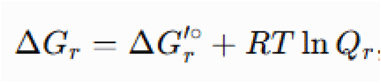

was not estimated in the present study. Negative values of E_r_^TW^ correspond to σ_r_=+1, whereas positive values correspond to σ_r_=−1. This sign-based classification reflects the model-defined reference energetic score and should not be interpreted as determining physiological reaction favorability or directionality.

### Model-derived metabolic order parameter

We defined the model-derived metabolic order parameter *m* as the mean alignment of reaction states:

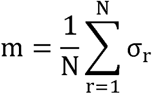

where *N* is the total number of reactions considered. In the patient-level formulation, reaction states were weighted by normalized activation scores (z_ir), such that the order parameter reflects directional alignment between intrinsic reaction directionality (σ_r) and patient-specific reaction activity rather than absolute activation magnitude.

The magnitude ∣*m*∣ quantifies global configurational coherence:

- ∣*m*∣=1: highly ordered metabolic state (uniform directional bias)
- ∣*m*∣=0: disordered state with balanced directional heterogeneity

This measure captures the degree of collective configurational alignment independent of transcriptomic subtype labels. In the present study, the total number of reactions (N = 235) corresponds to a reaction subnetwork retained after filtering for data and parameter availability, as described in Methods, rather than a set curated for biological centrality. The 235 reactions used in the SC-MSG model were, as a set, preferentially associated with genes showing medium-to-high relative expression in the Y497 cohort compared with the transcriptomic background (194/235 reactions, 82.6%; median percentile 57.3 vs. 49.9, Mann–Whitney p = 0.0055; **Supplementary Table 7**, **Supplementary Figure 1**). Of the 235 reactions, 221 (94%) were annotated to at least one KEGG metabolic pathway, with the remaining 14 lacking a curated pathway assignment (**Supplementary Table 8**). Restricting the analysis to this thermodynamically coherent reaction subset ensures computational tractability while preserving the dominant energetic degrees of freedom that shape the metabolic landscape.

### Patient-level extension of the order parameter

To extend this thermodynamic definition to patient-specific metabolic states, we introduced a weighted formulation of the order parameter that incorporates reaction-level activation derived from transcriptomic data. For each patient *i*, we defined:

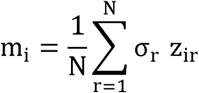

where *z_ir_* denotes the normalized activation score of reaction *r* in patient *i*. Normalization was performed across patients (z-score per reaction) to ensure comparability of activation magnitudes. In this formulation, σ_r_ encodes the intrinsic thermodynamic direction determined by the sign of the effective free energy, whereas *z_ir_* captures the patient-specific tendency toward reaction activation. Consequently, *m_i_* quantifies the directional alignment between intrinsic thermodynamic bias and observed reaction activity, rather than absolute expression or activation magnitude.

### Metabolic frustration index

To quantify higher-order thermodynamic incompatibilities within the metabolic network, we constructed a reaction–reaction interaction matrix *Brs*, encoding cooperative (*Brs* < 0) and antagonistic (*Brs* > 0) couplings derived from stoichiometric constraints, metabolite-sharing topology, and reaction coupling structure. Because the interaction term enters the Hamiltonian with a positive sign, cooperative interactions are represented by negative coupling coefficients that lower the system energy when compatible reactions are simultaneously active. The metabolic frustration index *F* was defined as:

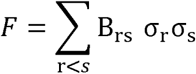

where *B_rs_* denotes the signed interaction between reactions *r* and *s*, and σ*_s_*∈{−1, +1} represents the thermodynamic state of reaction *r*. Negative couplings (*Brs*< 0) correspond to cooperative interactions, whereas positive couplings (*Brs* > 0) represent antagonistic interactions. For a symmetric interaction matrix *B* with zero diagonal, this expression can equivalently be written in quadratic form as

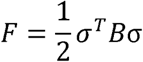

This quantity measures the extent to which the thermodynamic configuration {σr} is compatible or incompatible with the signed interaction structure of the metabolic network. To enable cross-patient comparison, we computed a normalized frustration score defined as

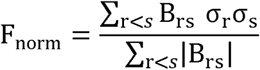

Under this definition, higher values of *F* _norm_ indicate greater incompatibility between the reaction configuration and the signed interaction network, reflecting increasingly frustrated and glass-like metabolic configurations.

### Definition of SC-MSG-derived subtypes

Patients were embedded in a two-dimensional configurational regime space defined by the magnitude of the order parameter (|m|) and normalized frustration (F_norm). Unsupervised clustering was performed in this thermodynamic coordinate system using Gaussian mixture modeling (GMM), with the optimal number of clusters determined based on the Bayesian information criterion (BIC). Applying this procedure to the order parameter and frustration index (see Methods above and the accompanying Supplementary Code), BIC favored a three-cluster solution over a five-cluster solution (ΔBIC = 25.0).

To preserve continuity with the five-subtype scheme used throughout this study, we report here the forced five-cluster (k = 5) solution, designated SC-MSG-derived subtypes T1–T5; the classification comprised T1 (n = 47), T2 (n = 109), T3 (n = 101), T4 (n = 119), and T5 (n = 121). Because BIC favors k = 3, the five-subtype structure should be regarded as a secondary, exploratory stratification rather than a statistically preferred classification; the corresponding results are presented as exploratory findings in Supplementary Figure 2 rather than as a main figure. These SC-MSG-derived subtypes were defined independently of the molecular expression-based classifications (i.e., stem-like, inflammatory, gastric, and intestinal) and were subsequently compared with molecular subtypes and clinical outcomes. Patient-level SC-MSG-derived subtype assignments, together with the corresponding molecular subtypes, order-parameter values, and normalized frustration scores, are provided in **Supplementary Data 4**.

### Statistical analysis

Differences in the order parameter and frustration score across molecular and SC-MSG-derived subtypes were assessed using the Kruskal–Wallis test. The association between molecular and SC-MSG-derived subtype classifications was evaluated using Pearson’s chi-square test of independence. Survival differences between SC-MSG-derived subtypes were assessed using Kaplan–Meier analysis and log-rank testing. Associations between thermodynamic coordinates and recurrence risk were evaluated using Cox proportional hazards modeling. Clinical covariates (age, sex, pathological stage) used in the Cox models are provided **in Supplementary Data 6**. All statistical tests were two-sided, and a *p*-value < 0.05 was considered statistically significant. All computations were performed in Python using NumPy and SciPy, and matrix operations were vectorized for computational stability.

### Pathway-level configuration diversity analysis

To quantify subtype-specific differences in metabolic organization, we computed pathway-level configuration diversity using the partition structure derived directly from HQA solutions. Each HQA solution decomposes the active reaction space into two disjoint subsets, *s*0 and *s*1, representing the OFF and ON components within the solution manifold. Reaction tokens provided by HQA were first mapped to KEGG reaction identifiers according to their index positions in the 235-reaction metabolic model, ensuring consistent alignment with curated KEGG pathway annotations. For each pathway *p*, the partition was projected as (*s*0∩*ρ*, *s*1∩*ρ*),preserving the internal OFF/ON decomposition rather than collapsing reactions into a single active set. For each molecular subtype (Stem-like and Inflammatory), we estimated the empirical distribution of distinct partition-preserved patterns across HQA solutions and computed Shannon configuration-diversity index as *S*α, *p* = −∑ρ(pattern)logρ(pattern), where *p*(pattern) denotes the relative frequency of each observed pathway-specific partition configuration. Subtype differences were quantified as Δ*S*p = *S*_Stem_, *p* − *S*_Infl_, such that positive values indicate greater configurational diversity in Stem-like tumors and negative values indicate greater diversity in Inflammatory tumors. Pathways with ∣Δ*Sp*∣ < 10^−3^ were excluded from visualization to remove negligible differences. Horizontal bar plots were generated in Python (NumPy, pandas, Matplotlib), with positive values displayed in dark red and negative values in dark blue, and exported at 600 dpi resolution and in vector format for publication. This partition-preserved entropy framework captures higher-order combinatorial organization of pathway states while retaining structural information embedded in the HQA solution decomposition.

## Discussion

This study presents a transcriptome-informed effective optimization-based representation of cancer metabolism and shows that malignant metabolic phenotypes can be described as collective low-energy configurations within a frustrated many-body optimization framework rather than solely as the cumulative outcome of isolated pathway-level alterations. [31] By unifying reaction-level Gibbs free energies, cofactor-mediated thermodynamic couplings, and patient-specific transcriptomic constraints within a single energy-based framework, the MSG model redefines the metabolic state of a tumor cell as a configurational regime which is a structured configuration of the optimization landscape shaped by gene expression and constrained by biochemical thermodynamics. [32] [33]

### Physical interpretation and scope of the SC-MSG framework

The SC-MSG model should be interpreted as a transcriptome-informed effective statistical-mechanical framework rather than a first-principles reconstruction of cellular thermodynamics. Although reference reaction Gibbs energies provide physically grounded energetic priors, the cofactor-interaction and transcriptomic-field terms are coarse-grained representations whose contributions depend on model construction and parameterization. Consequently, the resulting Hamiltonian values do not represent absolute cellular Gibbs free energies, and the optimized configurations should not be regarded as experimentally established equilibrium ground states. Similarly, the probability-derived landscapes represent effective statistical potentials, while the order and frustration parameters are collective, model-dependent descriptors of metabolic organization. The value of the framework therefore lies in integrating biochemical thermodynamic information and transcriptomic regulation into a unified network-constrained representation, rather than in claiming a complete microscopic thermodynamic description of cellular metabolism.

The subtype-associated metabolic organization observed in the present study is broadly concordant with independent multi-omics analyses of gastric cancer. Previous transcriptomic, spatial, organoid, and single-cell studies have demonstrated characteristic stem-like metabolic and phenotypic programs in gastric cancer.[21] [20] In parallel, integrated transcriptomic, metabolomic, and spatial-metabolomic studies have identified gastric cancer subtypes with differential enrichment of TCA-cycle and lipid metabolism, nucleotide and amino-acid metabolism, and glycan and energy metabolism.[34] Together, these independent observations support the broader interpretation that molecularly distinct gastric cancer subtypes exhibit differences in metabolic organization, consistent with the subtype-associated metabolic heterogeneity observed in our SC-MSG analysis.

### Reconceptualizing cancer metabolism as a configurational regime problem

The central conceptual contribution of this work is the shift from a pathway-centric to an optimization-centric description of cancer metabolism. Classical models identify which reactions are upregulated or downregulated in tumors, but they do not provide a model-based account of why certain global metabolic organizations are stable across heterogeneous genetic and environmental contexts.

Our model, in contrast, provides a framework in which metabolic reprogramming in cancer is represented as a collective reorganization of an effective optimization landscape, shaped by the interplay of intrinsic reaction thermodynamics, cofactor-mediated network coupling, and transcriptome-induced field perturbations; as discussed above, whether this reorganization is specifically attributable to the thermodynamic (ΔG-based) construction, as opposed to other similarly weighted combinations of correlated reaction-level measurements, remains an open question. The MSG model advances existing approaches in three respects. First, it provides a patient-specific metabolic optimization landscape whose topology is actively deformed by tumor transcriptomes, allowing distinct metabolic phenotypes to emerge as optimal configurations of the Hamiltonian rather than as predefined pathway templates. Second, it introduces a model-derived metabolic order parameter *m* that quantifies collective alignment between intrinsic reaction free-energy bias and patient-specific reaction activity, enabling continuous and subtype-agnostic patient stratification. Third, by formulating the biological problem as a QUBO, it establishes a computationally actionable interface between cancer metabolic modeling and quantum annealing hardware.

### The model-based role of frustration in metabolic heterogeneity

The many-body structure of the MSG Hamiltonian introduces frustration through competing cofactor demands, rendering the metabolic optimization landscape intrinsically rugged. The frustration represented in the SC-MSG model is motivated by competing cofactor demands within the biochemical architecture of metabolism, although its magnitude and landscape-level manifestation remain dependent on the model-defined interaction structure and parameterization. Cofactors such as NADH and ATP are simultaneously consumed and produced by multiple pathways, making it impossible for all reactions to be simultaneously favored within the model. The optimized reaction configuration therefore represents a compromise, namely a locally stable configuration that minimizes global energy under inherent biochemical constraints.

Different tumor subtypes occupy visually distinct regions of the patient-derived statistical-potential surfaces shown in **Figure 2C**, with differences in the empirical density and topology of patient states across subtypes. We caution, however, against interpreting these subtype-specific surfaces as establishing a fixed thermodynamic hierarchy of region depth or stability across subtypes: our reference-independent analysis of the analogous stem-like-versus-inflammatory comparison (see Discussion, “The model-derived order parameter as a continuous, independent prognostic axis,” and **Supplementary Figure 4**) found no significant difference in the order parameter m between these two subtypes when evaluated without a subtype-discriminant projection axis (Mann–Whitney p = 0.69), and no discernible double-well structure separating them. We therefore regard the subtype-specific density-region geometries in **Figure 2C** as descriptive of the empirical distribution of patient states in each subtype’s own reduced coordinate space, rather than as established evidence of a specific, ranked thermodynamic stability hierarchy (e.g., that stem-like tumors are uniquely or maximally stabilized) across subtypes.

The subtype-dependent metabolic and optimization landscapes identified in the present study are broadly consistent with independent molecular and multi-omics evidence. Previous transcriptomic, spatial-transcriptomic, patient-derived xenograft, organoid, and single-cell studies have demonstrated that the gastric, inflammatory, intestinal, stem-like, and mixed molecular subtypes represent biologically distinct tumor states characterized by differences in tumor-cell programs, stromal interactions, immune contexture, epithelial–mesenchymal plasticity, and clinical behavior. In particular, mesenchymal/stem-like gastric cancers exhibit tumor-cell-intrinsic TGF-β activation, partial epithelial–mesenchymal transition, stem-like properties, and treatment resistance. [21] Independent integrated transcriptomic, metabolomic, and spatial-metabolomic studies have further identified gastric cancer subgroups with differential enrichment of TCA-cycle and lipid metabolism, nucleotide and amino-acid metabolism, and glycan and energy metabolism.[35] High-resolution spatial metabolomic profiling has also revealed metabolically distinct gastric cancer subtypes associated with differences in nucleotide abundance, DNA metabolism, immune-cell composition, clinicopathological characteristics, and clinical outcomes. [34] Collectively, these observations support the biological plausibility of the subtype-associated metabolic heterogeneity captured by the SC-MSG framework. Because these independent studies did not directly measure Hamiltonian energy, order parameters, or metabolic frustration, they provide biologically concordant evidence rather than direct external validation of the SC-MSG-derived optimization landscapes.

We addressed this directly with four null-model control analyses (full methodology and code provided in the accompanying Supplementary Code repository). (H0-A) To test whether the discrete SC-MSG-derived subtypes (T1–T5) simply reproduce a naive transcriptomic clustering, we compared them with a five-cluster solution obtained by Gaussian mixture clustering on the leading transcriptomic principal components (PC1–PC5) of the same 235-reaction expression matrix. Agreement was minimal (adjusted Rand index = 0.021; best one-to-one matched overlap = 29.2%), indicating that the SC-MSG-derived subtypes are not a re-description of transcriptomic clusters. (H0-B) To test whether the survival association of the continuous order parameter m depends specifically on the thermodynamic (ΔG-based) sign convention σ, rather than on any similarly constructed balanced combination of the 235 correlated reaction z-scores, we recomputed m using permuted sign vectors and compared the resulting univariate Cox coefficient for m to the coefficient obtained with the real σ. Under full, reaction-independent permutation (1,000 permutations), the real coefficient was not distinguishable from the null distribution (empirical *p* = 0.524). To ask whether this reflected the correlation structure among reactions within the same metabolic pathway rather than a property of σ itself, we repeated the test under two constrained permutation schemes that respect KEGG pathway membership (51 pathway blocks spanning the 235 reactions): permuting the overall sign of each pathway block as a unit while preserving each block’s internal (within-pathway) sign pattern exactly as observed (empirical *p* = 0.258), and permuting signs only within each pathway block while preserving the real between-pathway pattern exactly as observed (empirical *p* = 0.907, i.e., the real within-pathway assignment performed **worse** than the median permutation). None of the three permutation schemes supported a specific role for the thermodynamic sign convention. (**Supplementary Table 9** and **Supplementary Figure 3**) (H0-C) To test whether the survival association of the frustration index depends on the specific topology of the reaction-pair coupling matrix B, we compared the real coefficient to a null distribution obtained from 200 degree- and sign-preserving rewirings of B (double-edge-swap, separately for positive and negative edges). The real coefficient exceeded the rewired-network distribution more often than chance but did not reach conventional significance (empirical *p* = 0.105). (H0-D) Head-to-head comparison of the SC-MSG-derived subtypes against the size-matched transcriptomic PC-based five-cluster solution (H0-A) showed neither classification to be significantly associated with survival (log-rank *p* = 0.295 vs. 0.197 for OS), with near-identical discrimination (C-index 0.528 vs. 0.527), indicating that the discrete subtype scheme confers neither an advantage nor a disadvantage relative to a naive transcriptomic baseline. Taken together, these null-model analyses show that the discrete SC-MSG-derived subtypes perform comparably to, rather than better than, naive transcriptomic clustering (H0-A), and they also show that the prognostic association of the continuous order parameter m is statistically robust yet not demonstrably specific to the thermodynamic (ΔG-based) sign convention used to construct it: a similarly constructed balanced linear combination of the same 235 correlated reaction z-scores, with signs assigned independently of ΔG, achieves comparable predictive power. We regard this as an important open question rather than a settled negative result, and we did not identify a specific alternative explanation (e.g., a dominant co-expression axis correlated with both m and outcome) within the scope of the present analysis. We report it transparently here because it directly bears on the causal interpretation of our central claim, and we encourage future work—ideally incorporating orthogonal, non-transcriptomic thermodynamic measurements—to resolve whether the reaction-specific thermodynamic direction is mechanistically responsible for the observed prognostic association, or whether it is one of many similarly weighted combinations of correlated metabolic reaction activity that would perform comparably. The B-matrix network term did not show statistically significant specificity (H0-C, p = 0.105), and the discrete subtype scheme performed comparably to, rather than better than, a naive transcriptomic baseline (H0-D).

(H0-A′) To determine whether the ΔG and B-matrix terms are independently necessary for the subtype-level Hamiltonian optimization itself — as distinct from H0-B/C, which test the downstream patient-level order parameter — we re-solved H^(p)^(x) for each of the five molecular subtypes across the full published w□ –w□ sweep (605 subtype × weight combinations) with the ΔG coefficient and w jointly set to zero. In every combination, the ablated model collapsed to a trivial, degenerate solution (all 235 reactions inactive), because the transcriptomic field term S is strictly positive for every reaction in every subtype; this is a structural property of the linear field term rather than direct evidence of thermodynamic specificity. (H0-A″) An initial decomposition comparing ΔG-only and B-matrix-only reconstructions of the full-model solution across w□ was confounded in two ways: at low w□ the B-matrix-only model inherits the same all-positive-field degeneracy tested in H0-A′, and the ΔG-only model (fixed at w□=0) is a static reference whose apparent “decline” against the full model at increasing w partly reflects the full model moving away from that fixed point rather than any change in ΔG’s importance — confirmed directly by the near-identical decay when the full-model solution at w□ =0.1 is compared against itself at higher w (Jaccard 0.42–0.43 at w□ ≥1.1, matching the ΔG-only decay). To remove both confounds, we instead permuted the identities of the 235 ΔG values across reactions (preserving their distribution), re-solved the complete Hamiltonian (true B-matrix and transcriptomic field, permuted ΔG) at the same (w□, w□) as the full model, and compared the resulting configuration to the full-model solution by Jaccard — repeated 50–100 times per subtype at w□ {0.1, 1.1, 2.1} (w□ =1.1) across all five molecular subtypes. Because the configuration obtained with the true, unpermuted ΔG assignment is by construction the full-model solution itself (Jaccard = 1.00), no permutation replicate can exceed it and no empirical p-value is defined for this comparison. The informative quantity is instead the extent to which permuting ΔG identities displaces the optimized configuration. Across 750 replicates, the permuted-ΔG Jaccard averaged 0.26 at w□ = 0.1 (maximum ≤ 0.35), 0.81 at w□ = 1.1 (≤ 0.87) and 0.84 at w□ = 2.1 (≤ 0.88); equivalently, the fraction of the optimized configuration attributable to the specific reaction-level assignment of ΔG values is approximately 0.74 at w□ = 0.1 but only 0.16–0.19 at w□ ≥ 1.1. The specific assignment of ΔG values — not merely the presence of some sign-varying linear term — therefore shapes the optimized configuration, but this contribution is strongly weight-dependent: it dominates in the low-coupling regime and attenuates sharply as w□ increases, where permuted ΔG assignments already reproduce 81–84% of the configuration. These Jaccard comparisons are not subject to solver noise: the SC-MSG optimization at each (w□, w□) is solved to global optimality by a classical integer-programming solver, which returns identical optimal configurations to the HQA solver for this problem size (see Methods); repeated solves at the same (w□, w□) are therefore deterministic, and the reported Jaccard values reflect the permutation of ΔG identities alone. We therefore conclude that the thermodynamic (ΔG) term contributes real, reaction-specific information to the optimized Hamiltonian configuration, while this configuration-level specificity does not translate into ΔG-specific prognostic information at the patient level (H0-B, H0-C, H0-D). (**Supplementary Table 11**, **Supplementary Figure 5**)

### Interpreting m as a network-coherent composite biomarker

Because the null-model analyses above indicate that the prognostic value of m is not specifically attributable to the ΔG-based sign convention, its clinical utility is better understood as arising from the coordinated, network-constrained structure of the underlying reaction set than from a validated, reaction-level thermodynamic directionality. The 235 reactions entering the model were not sampled at random: they were selected jointly for GPR annotation, transcriptomic mapping, and reference free-energy data, and are further coupled through the B-matrix by shared redox and energy cofactors, for which H0-C did not reach statistical significance (p = 0.105), in contrast to the null result for the ΔG sign vector alone. We therefore interpret m primarily as a structured composite score of coordinated expression across a curated, cofactor-linked metabolic reaction network, of which the specific ΔG-based linear combination tested here is one of several similarly weighted combinations of correlated reaction activity that could achieve comparable prognostic performance. This interpretation is consistent with the reaction-level contribution patterns in **Figure 1G** and **Figure 3F-G**, which show that redox- and cofactor-associated reactions contribute disproportionately to subtype-specific signal, and with independent multi-omics evidence that these same pathways are recurrently reprogrammed across gastric cancer subtypes. Under this view, the prognostic value of m reflects the biological coherence of coordinated metabolic gene co-expression within a transcriptome-informed, cofactor-coupled network, rather than a demonstrated causal role for the specific direction of individual reaction free energies. Distinguishing co-expression-driven coherence from genuine thermodynamic directionality will require future work incorporating patient-specific metabolomics or isotope-resolved flux measurements that can probe reaction directionality independently of transcriptomic measurement.

### The model-derived order parameter as a continuous, independent prognostic axis

The continuous model-derived metabolic order parameter m, rather than the discrete SC-MSG-derived subtypes, provides the most robust evidence that this framework captures clinically relevant information beyond conventional classification: in multivariable Cox models adjusted for clinical covariates and the dominant transcriptomic principal components, m remained independently associated with both overall and disease-free survival (per-1-SD hazard ratio 1.25 and 1.23, respectively), albeit with only a modest improvement in model discrimination (ΔC-index = +0.004 for OS and +0.002 for DFS). In contrast, discrete SC-MSG-derived subtypes T1–T5, obtained by forcing a five-cluster solution on (|m|, F_norm) for continuity with our original analysis plan, did not correspond to significantly different survival outcomes overall (log-rank *p* = 0.295, whole cohort), and Bayesian information criterion in fact favored a three-cluster solution, indicating that the five-subtype structure is not strongly supported by the data. An exploratory, secondary comparison identified T1 and T3 as the most divergent pair within stem-like tumors (log-rank *p* = 0.044 for OS, 0.037 for DFS), with T1 (n = 9) showing higher order but **lower** (more cooperative) frustration than T3.

Because this comparison was one of ten pairwise tests without correction for multiple comparisons and was based on a small subgroup, we regard it as hypothesis-generating rather than confirmatory, and present it in **Supplementary Figure 2** rather than as a main finding. A metabolic-entropy comparison for this analysis could not be recomputed, because it requires patient-specific optimized binary reaction-participation states from the HQA solver that were not available. Taken together, these results indicate that the principal clinically actionable signal in this framework is the continuous order parameter m itself, rather than a specific discrete SC-MSG-derived subtype or a particular combination of high order and high frustration.

### Hybrid quantum annealing as a solver for metabolic configuration optimization

The equivalence of the MSG Hamiltonian to a QUBO problem provides a direct interface with quantum optimization hardware. Importantly, quantum annealing serves here strictly as a computational solver for the classical energy minimization problem; no quantum mechanical effects are invoked in the biological model itself. For the present problem scale of 235 reactions, D-Wave HQA and classical integer programming solver yielded identical solutions, confirming global optimality of the identified configurations and establishing that the biological conclusions of the MSG model are not contingent on the choice of solver. As the metabolic network is extended toward genome scale, however, the combinatorial complexity of the configuration space will increasingly challenge classical approaches. This challenge arises less from the raw size of the binary search space, which is already astronomically large at 235 reactions yet remains tractable for classical integer programming, than from the density of pairwise couplings introduced by the B-matrix: unlike the sparse stoichiometric matrices used in constraint-based flux methods, cofactor sharing links reactions across otherwise distant regions of metabolism, and the number of such pairwise terms grows combinatorially as the network is extended toward genome-scale reconstructions comprising on the order of several thousand to over ten thousand reactions. Prior benchmarking studies indicate that HQA offers systematic advantages precisely for this class of dense, high-dimensional QUBO instances, where classical branch-and-bound solvers lose the sparsity structure they rely on for efficient pruning; identifying the specific network size at which this crossover occurs for genome-scale metabolic Hamiltonians is an important direction for future benchmarking. This positions the MSG framework as a biologically motivated application domain for near-term quantum annealing, one in which the QUBO formulation arises naturally from the physics of the biological problem.

### Limitations and future directions

The SC-MSG framework is an effective statistical-mechanical model and does not constitute a first-principles reconstruction of cellular thermodynamics. The Hamiltonian combines reference reaction energetics with coarse-grained cofactor-interaction and transcriptomic-field terms; therefore, its numerical values should be interpreted as relative model-dependent scores rather than absolute cellular Gibbs free energies. The probability-derived effective free-energy landscapes similarly characterize the statistical organization of patient states and do not directly quantify biochemical free-energy differences or kinetic transition barriers. In addition, the effective order parameter is a coarse-grained measure of thermodynamically weighted reaction alignment rather than an equilibrium order parameter derived from a microscopic partition function. The binary representation of reaction activation does not capture continuous metabolic flux, and the current framework does not explicitly incorporate intracellular metabolite concentrations, reaction quotients, chemical potentials, spatial compartmentalization, temperature-dependent dynamics, or non-equilibrium entropy production.

The reaction network analyzed in this study does not represent a complete genome-scale reconstruction and is not assumed to preserve the full global stoichiometric, topological, or thermodynamic structure of human cellular metabolism. Because reaction inclusion was constrained by the joint availability of human annotation, GPR information, transcriptomic mapping, and reference 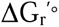 values, the resulting network may underrepresent reactions or pathways with incomplete annotation or unavailable thermodynamic data. Moreover, because the SC-MSG interaction matrix and model-derived landscape depend on the composition and topology of the included reaction set, reaction-coverage and network-composition effects may influence the inferred order, frustration, and configurational structure.

The current SC-MSG framework does not explicitly evaluate whether each optimized reaction-participation configuration forms a fully connected metabolic network, supports a non-zero mass-balanced steady-state flux distribution, preserves essential metabolic tasks, or excludes thermodynamically infeasible internal cycles. Therefore, the inferred configurations should not be interpreted as complete biochemical flux states: in the absence of an enforced mass-balance constraint, an optimized reaction-participation pattern reflects the model’s thermodynamic and transcriptomic tendency, or bias, toward activating a given reaction, not a quantitative metabolic flux through it. Integration with genome-scale constraint-based modeling, flux-balance analysis, thermodynamic flux analysis, and metabolic-task feasibility testing will be required to evaluate the stoichiometric and thermodynamic feasibility of individual model-derived configurations. [28] Because transcriptomic data contribute directly to the model, part of the observed subtype and survival structure may reflect organization already present in the underlying transcriptomic manifold. Future studies should therefore incorporate metabolomics, isotope-resolved flux measurements, patient-specific reaction free energies, independent clinical cohorts, and transcriptome-only and network-randomized control models to determine the specific contribution of thermodynamic constraints beyond transcriptomic structure. Extension to genome-scale reaction networks and continuous or probabilistic reaction variables may further improve the biochemical realism and generalizability of the framework.

## Conclusion

This work establishes a transcriptome-informed spin-glass optimization framework for representing cancer metabolic organization and demonstrates its compatibility with quantum annealing–based optimization. By integrating reference biochemical energetics, cofactor-mediated network interactions, transcriptome-derived regulatory fields, and QUBO-based optimization within a unified effective Hamiltonian, the SC-MSG model provides a network-level representation of metabolic heterogeneity. The resultinglow-objective configurations, optimization landscapes, and collective order parameters should be interpreted as model-dependent descriptors rather than direct measurements of cellular Gibbs free energy or equilibrium configurational regimes. Within this scope, the framework provides a computational bridge between biochemical thermodynamic information, transcriptomic heterogeneity, and complex-systems optimization and offers a scalable strategy for patient-specific metabolic characterization.

## Code Availability

All analysis and visualization scripts used in this study, including those for principal-component analysis, Gaussian mixture model clustering with Bayesian information criterion– based model selection, pathway-level analysis and visualization, configuration-landscape analysis, survival analysis, and generation of the corresponding figures and subpanels, are provided as Supplementary Code.

## Declarations

### Ethics approval and consent to participate

N/A

### Consent for publication

N/A

### Availability of data and materials

All data, quantum computing codes, and supporting materials underlying the findings of this study are available in the Supplementary Materials accompanying this article.

## AI Use Declaration

AI tools were used only for English grammar correction and language polishing.

## Authors’ contributions

Conceptualization: JYS, JHC; Methodology: JYS, KHB; Data analysis: JYS; Investigation: JYS, KHB, IBP; Writing-original draft: JYS; Writing-review & editing: JYS, KHB, JHC; Supervision: JYS, JHC; Project administration: JYS, JHC; Funding acquisition: JHC; Formal analysis: JYS; Interpretation of the results: JYS, JHC, KHB, JB. All authors have read and agreed to the published version of the manuscript.

## Acknowledgements

MID (Medical Illustration & Design), as a member of the Medical Research Support Services of Yonsei University College of Medicine, providing excellent support with medical illustration.

---

**Supplementary Table 1. Summary of benchmark analyses evaluating the incremental contribution of the SC-MSG model-derived metabolic order parameter (m).**

This table summarizes the benchmark analyses performed to evaluate whether the SC-MSG-derived model-derived metabolic order parameter (m) provides additional prognostic information beyond conventional clinical variables and transcriptomic principal components (PCs). Cohort size, PCA summary, variance explained by transcriptomic PCs, and multivariable survival model performance are summarized.

**Supplementary Table 2. Association between transcriptomic principal components and the SC-MSG model-derived metabolic order parameter.**

This table reports the proportion of transcriptomic variance explained by each principal component (PC1–PC10) together with its Pearson correlation with the SC-MSG model-derived metabolic order parameter (m). *P* values were calculated using two-sided Pearson correlation tests.

**Supplementary Table 3. Benchmark comparison of Cox proportional hazards models with and without the SC-MSG model-derived metabolic order parameter.**

This table compares Cox proportional hazards models for overall survival (OS) and disease-free survival (DFS) using clinical covariates alone, transcriptomic principal components, and the SC-MSG model-derived metabolic order parameter (m). Model performance is summarized by log-likelihood, Akaike information criterion (AIC), likelihood-ratio statistics, and concordance index (C-index).

**Supplementary Table 4. Regression coefficients of multivariable Cox proportional hazards models.**

This table provides the estimated regression coefficients, hazard ratios (HRs), 95% confidence intervals (CIs), and two-sided Wald-test *P* values for all variables included in the multivariable Cox proportional hazards models presented in Supplementary Table 3.

**Supplementary Table 5. Performance of transcriptomic principal component–based subtype classification with and without the SC-MSG model-derived metabolic order parameter.**

This table summarizes five-fold cross-validation results for molecular subtype classification using transcriptomic principal components (PC5 or PC10) with or without inclusion of the SC-MSG model-derived metabolic order parameter (m). Classification performance is reported as overall accuracy, balanced accuracy, and macro F1 score.

**Supplementary Table 6. Patient-level transcriptomic principal component scores and SC-MSG model-derived metabolic order parameter.**

This table lists the molecular subtype assignment, SC-MSG model-derived metabolic order parameter (m), and transcriptomic principal component scores (PC1–PC10) for each patient included in the benchmark analyses. These data were used for the correlation, classification, and multivariable survival analyses described in the manuscript.

**Supplementary Table 7. Relative expression level of the 235 metabolic reactions used in the SC-MSG model compared to the whole transcriptome.**

This table reports, for each of the 235 reactions included in the SC-MSG Hamiltonian, the cohort-mean expression level of the mapped gene(s) in the Y497 gastric cancer cohort (n = 497 patients) and its percentile rank within the full 12,922-gene expression panel. Reactions are ranked in descending order of expression percentile and classified as High (≥66.7th percentile), Medium (33.3–66.7th percentile), or Low (<33.3rd percentile). For each reaction, the number of mapped genes, the number of genes covered by the expression panel, and any genes not represented in the panel are also listed. These values were used to assess the relative transcriptomic relevance of the 235 reactions and are summarized in Supplementary Figure 1.

**Supplementary Table 8. Pathway and gene annotations for the 235 metabolic reactions used in the SC-MSG model.**

This table provides, for each of the 235 reactions included in the SC-MSG Hamiltonian, the associated gene(s) used to construct the reaction-level expression values (as described for Supplementary Data 2) together with the corresponding KEGG pathway annotation(s). Pathway assignments are reported as both KEGG pathway identifiers and pathway names, retrieved from the KEGG PATHWAY database. Of the 235 reactions, 221 (94%) are annotated to at least one KEGG metabolic pathway; the remaining 14 reactions, which lack a curated KEGG pathway assignment, are marked as “Not assigned.” This table is provided to clarify the biological context and pathway membership of the reactions used in the model.

**Supplementary Table 9. Null-model control analyses for the model-derived order parameter and frustration index.**

This table reports four null-model analyses (H0-A to H0-D) testing whether the prognostic and classificatory value of the SC-MSG framework depends on its specific thermodynamic construction, rather than on generic properties of aggregating correlated reaction-level measurements. For each analysis, the question addressed, method, test statistic or metric, result, and interpretation are provided. Full methodology and code are provided in nullmodel_analysis/ (Supplementary Code).

**Supplementary Table 10. Sensitivity analyses for the Cox model association of the order parameter m.**

This table reports five sensitivity analyses probing the robustness of the independent association between the model-derived order parameter m and overall survival (OS) and disease-free survival (DFS) in the multivariable Cox model (clinical covariates + PC1–PC5 + m), including extension of the principal-component adjustment from PC1–PC5 to PC1–PC20, adjustment for clinical molecular subtype, a bootstrap confidence interval for the incremental C-index attributable to m, five-fold cross-validated model performance, and a Schoenfeld residual test of the proportional-hazards assumption. Hazard ratios for m are expressed per 1-SD increase (SD = 0.147) for interpretability. Full methodology and code are provided in sensitivity_analysis/ (Supplementary Code).

**Supplementary Table 11. H0-A′: Ablation of the Hamiltonian optimization itself (ΔG and B-matrix terms removed).**

Complementary to H0-A (which tests whether the discrete SC-MSG-derived subtypes reproduce naive transcriptomic clustering), this table reports analyses testing whether the ΔG (reaction free-energy) and B-matrix (cofactor-coupling) terms of the Hamiltonian materially change the optimized binary reaction configuration itself, independent of any downstream prognostic readout. Reported analyses include joint removal of both terms (H0-A′(i)), a decomposition into ΔG-only and B-matrix-only contributions (H0-A′(ii), superseded and retained for transparency), and a permutation-based test of whether the true, reaction-specific ΔG assignment reproduces the full-model configuration better than randomized reassignments at matched coupling weights (H0-A″). For each analysis, the question addressed, method, test statistic or metric, result, and interpretation are provided. Full grid results are provided in the accompanying Data sheet, and code is provided in nullmodel_analysis/ (Supplementary Code).

**Supplementary Data 1. Human metabolic reactions and thermodynamic annotations**

This table contains the 235 human metabolic reactions included in the Metabolic Spin-Glass (MSG) model. For each reaction, the KEGG reaction identifier, reaction equation, standard transformed Gibbs free energy (ΔG′), and associated reaction annotations are provided. These reaction-specific thermodynamic parameters were used to construct the intrinsic energy term of the metabolic Hamiltonian.

**Supplementary Data 2. Patient-by-reaction expression matrix**

This table contains the reaction-level expression matrix for the gastric cancer cohort (n = 497). Gene expression values were mapped to metabolic reactions by averaging the expression levels of genes associated with each reaction. Each row represents a patient, and each column represents one of the 235 metabolic reactions included in the Hamiltonian model. These values were used to construct the patient-specific transcriptomic field.

**Supplementary Data 3. Cofactor interaction (B) matrix**

This table provides the pairwise cofactor interaction matrix (B matrix) describing model-defined cooperative and competition among the 235 metabolic reactions. Positive values indicate cooperative interactions, whereas negative values indicate competitive interactions mediated through shared energy or redox cofactors. The B matrix constitutes the pairwise interaction term of the Metabolic Spin-Glass Hamiltonian.

**Supplementary Data 4. Clinical and SC-MSG-derived subtype annotations**

This table summarizes the patient identifiers together with the corresponding molecular subtype and model-derived SC-MSG-derived subtype assignments. The SC-MSG-derived subtype was obtained from clustering based on the Hamiltonian-derived order parameter and frustration score and was used for downstream survival and comparative analyses.

**Supplementary Data 5. Reaction annotations for Figure 3G**

This table provides detailed annotations for the reactions displayed in Figure 3G, including KEGG reaction identifiers, reaction names, reaction equations, reference ΔG′ values, and pathway classifications. These annotations facilitate biological interpretation of the reaction-level thermodynamic contributions shown in Figure 3G.

**Supplementary Data 6. Clinical information of the Y497 gastric cancer cohort**

This table contains the clinical information for the 497 gastric cancer patients analyzed in this study, including patient identifiers, molecular subtype, demographic characteristics, pathological stage, survival outcomes, recurrence status, and other clinicopathological variables used in the statistical analyses.

**Supplementary Figure 1.**
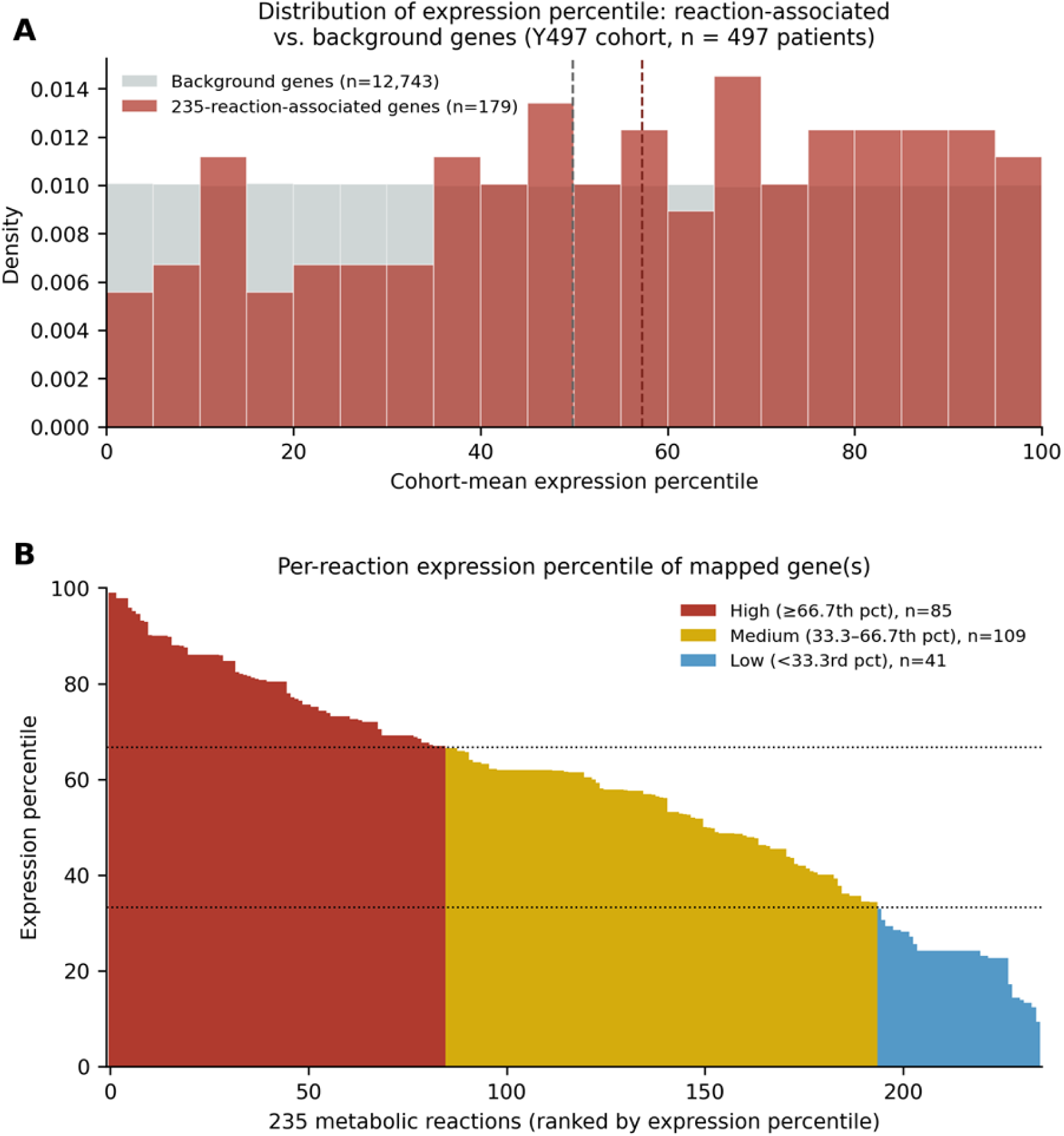
Relative expression level of the 235 metabolic reactions used in the SC-MSG model compared to the whole transcriptome. (A) Distribution of cohort-mean expression percentile for genes mapped to the 235 reactions (n = 179) versus all remaining genes in the expression panel (n = 12,743), in the Y497 gastric cancer cohort (n = 497 patients). Dashed lines indicate group medians. Genes associated with the 235 reactions showed significantly higher expression percentiles than background genes (median 57.3 vs. 49.9; two-sided Mann–Whitney U test, P = 0.0055). (B) Per-reaction expression percentile of the mapped gene(s), ranked in descending order across all 235 reactions and color-coded by relative expression category: High (≥66.7th percentile, red, n = 85), Medium (33.3–66.7th percentile, yellow, n = 109), and Low (<33.3rd percentile, blue, n = 41). Dotted horizontal lines indicate the tertile thresholds (33.3rd and 66.7th percentile). Underlying values are provided in Supplementary Table 7.

**Supplementary Figure 2.**
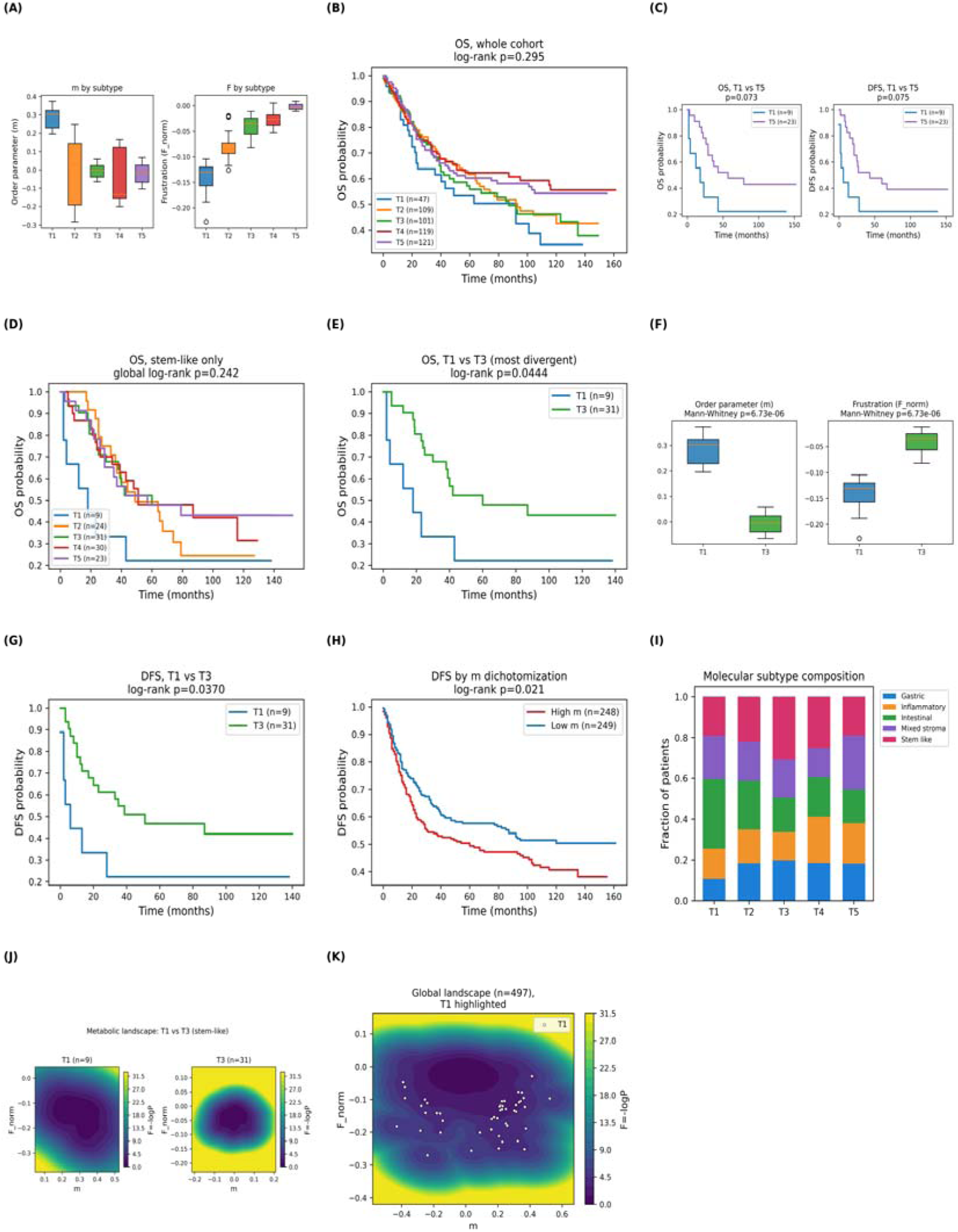
Exploratory analysis of discrete SC-MSG-derived subtypes (T1–T5) based on the order parameter and frustration index. T1–T5 were derived by Gaussian mixture clustering on (|m|, F_norm) (k = 5 forced for continuity with the five-subtype scheme used throughout this study; BIC favored k = 3). (A) Order parameter (m) and frustration (F_norm) by subtype within stem-like tumors. (B) Overall survival (OS) by subtype, whole cohort (log-rank *p* = 0.295). (C) OS and disease-free survival (DFS), T1 vs T5 within stem-like tumors (not significant). (D) OS by subtype, stem-like tumors only (log-rank *p* = 0.242). Sample sizes within stem-like tumors: T1 (n = 9), T2 (n = 24), T3 (n = 31), T4 (n = 30), T5 (n = 23). (E) OS, T1 vs T3 (most divergent pair by pairwise testing; log-rank *p* = 0.044). (F) Order parameter and frustration, T1 vs T3, shown for reference only; T1 and T3 were defined by clustering on these two variables, so the between-group difference is circular by construction and no test statistic is reported. (G) DFS, T1 vs T3, stem-like tumors (log-rank *p* = 0.037; not significant across the whole cohort, *p* = 0.677). (H) DFS by order parameter (m) dichotomization (reproduced from Figure 4D). (I) Molecular subtype composition within each SC-MSG-derived subtype. (J) Metabolic optimization landscape, T1 vs T3, stem-like tumors. (K) Global metabolic optimization landscape (n = 497), T1 highlighted. Given the small size of the T1 subgroup (n = 9 within stem-like tumors) and the absence of correction for multiple comparisons (10 pairwise tests across subtypes), the discrete-subtype findings in this figure should be considered hypothesis-generating rather than confirmatory. A metabolic-entropy comparison for this analysis could not be recomputed, because it requires patient-specific optimized binary reaction-participation states from the HQA solver that were not available.

**Supplementary Figure 3.**
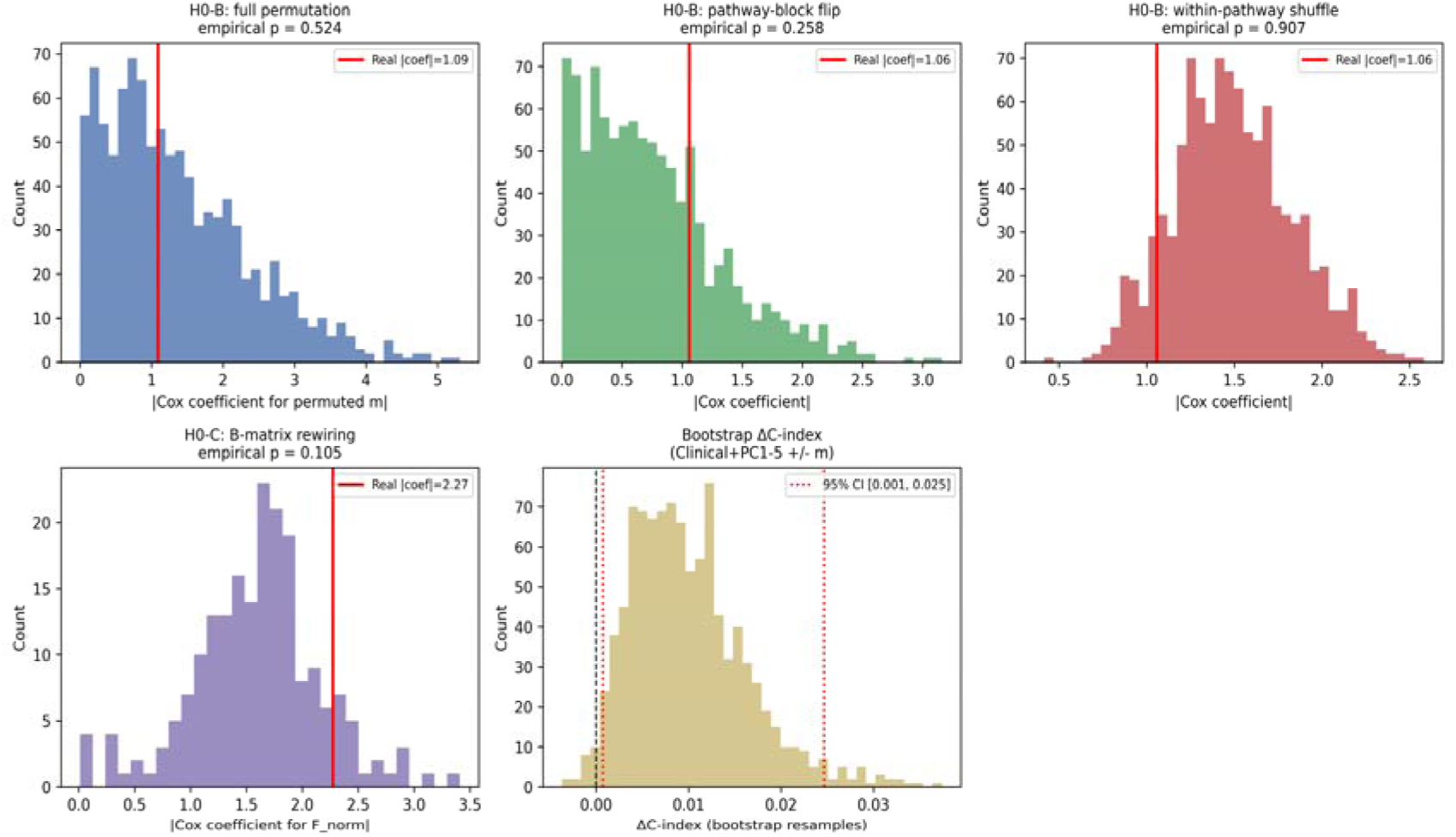
Null-model permutation distributions and bootstrap diagnostics for the order parameter and frustration index. Top row, left to right: null distributions of the absolute univariate Cox coefficient for m under three permutation schemes for the thermodynamic sign convention σ□ — full permutation across all 235 reactions (empirical *p* = 0.524), whole-pathway sign flip preserving each of 51 KEGG pathway blocks’ internal pattern (empirical *p* = 0.258), and within-pathway-only shuffle preserving the real between-pathway pattern (empirical *p* = 0.907). Red lines mark the real (ΔG-based) coefficient. Bottom row: null distribution of the absolute Cox coefficient for the frustration index F_norm under 200 degree- and sign-preserving rewirings of the reaction-pair coupling matrix B (empirical *p* = 0.105); and the bootstrap distribution (1,000 resamples) of the incremental C-index attributable to m in the Clinical+PC1–5 model, with the 95% CI [0.001, 0.025] marked (dotted red lines) relative to zero (dashed black line). See Supplementary Table 9 (null-model analyses) and Supplementary Table 10 (sensitivity analyses) for full numerical results, and the accompanying Supplementary Code repository (nullmodel_analysis/, sensitivity_analysis/) for methodology.

**Supplementary Figure 4.**
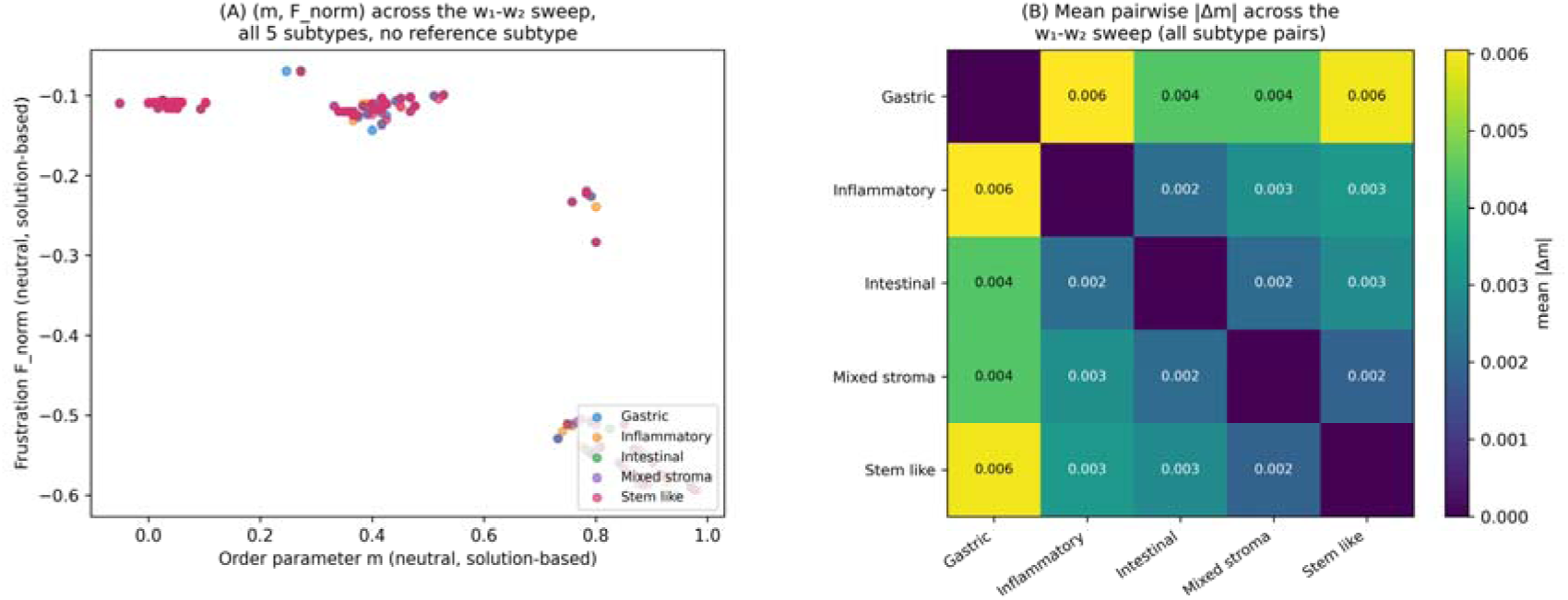
Testing whether apparent stem-like subtype separation depends on the choice of reference subtype. The order parameter m and frustration index F_norm were recomputed directly from the HQA solution ensembles across the w□ –w□ sweep for all five molecular subtypes, without designating any subtype as a comparison reference (see Discussion). (A) Overlay of (m, F_norm) for all five subtypes across the w□ – w □sweep. Subtypes occupy highly overlapping regions of this neutral space, with no visually distinct clustering by subtype. (B) Mean pairwise |Δm| across the w□ –w□ sweep for all ten subtype pairs. Stem-like tumors are not more distant from the other four subtypes (mean |Δm| = 0.003–0.006) than those subtypes are from one another (e.g., Gastric vs. Inflammatory = 0.006), indicating that the apparent stem-like separation observed under a stem-like-referenced projection reflects the choice of reference rather than an intrinsic thermodynamic distinction between subtypes.

**Supplementary Figure 5.**
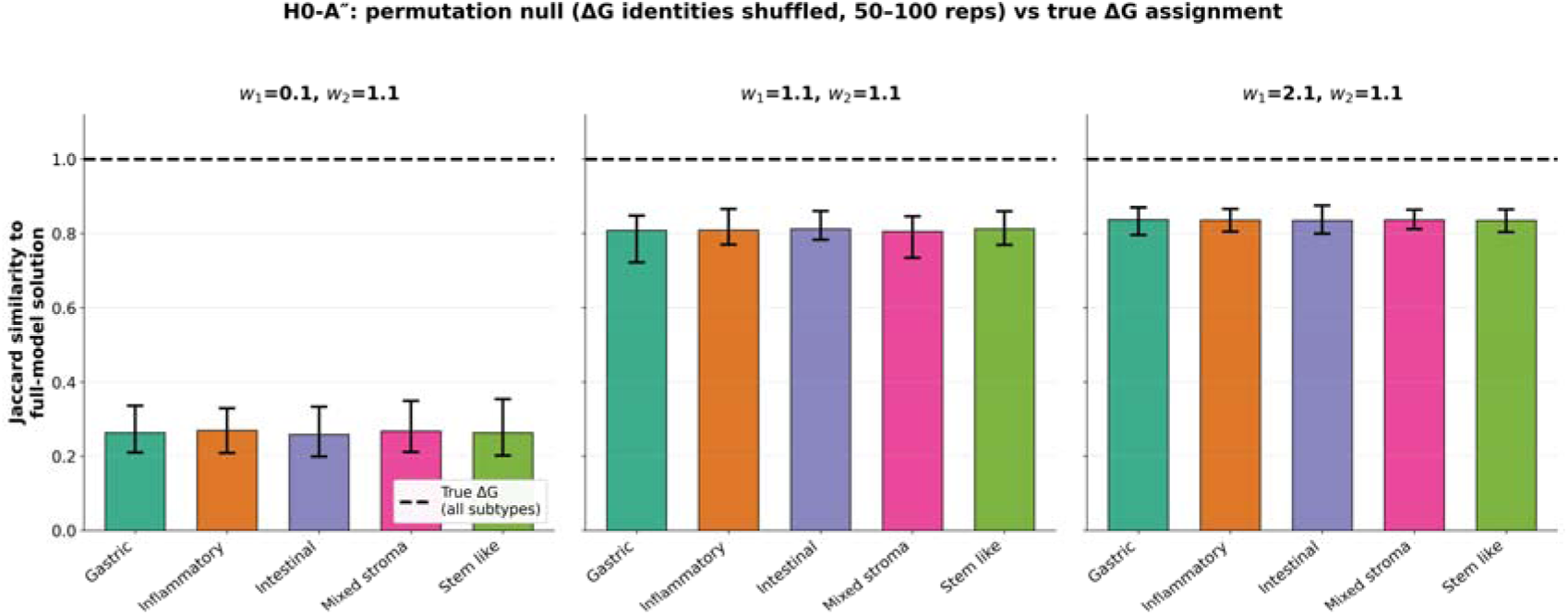
H0-A″: permutation control for ΔG specificity in the Hamiltonian optimization. For each of five molecular subtypes and three representative coupling weights (w□ = 0.1, 1.1, 2.1; w□ = 1.1), the identities of the 235 reference ΔG values were permuted across reactions (distribution preserved), the complete Hamiltonian (true B-matrix and transcriptomic field, permuted ΔG) was re-solved, and the resulting configuration was compared to the full-model solution by Jaccard similarity (50–100 permutation replicates per subtype × weight combination; bars show mean ± range). The dashed line marks the Jaccard obtained with the true, unpermuted ΔG assignment (1.00 in every subtype and weight, since this configuration is the full-model solution itself). Permuting ΔG identities displaced the optimized configuration to a degree that grew smaller as w□ increased: across 750 replicates, the permuted-ΔG Jaccard averaged 0.26 at w□ = 0.1, 0.81 at w□ = 1.1, and 0.84 at w□ = 2.1, indicating that the specific reaction-level assignment of ΔG values — not merely the presence of some sign-varying linear term — shapes the optimized configuration, most strongly at low coupling weight. This result also serves as the negative control for the earlier, confounded ΔG-only vs. B-matrix-only decomposition (see main text H0-A″), which compared a fixed w □=0 reference against a moving full-model target and is superseded by this analysis. Full numerical results are provided in **Supplementary Table 11**.

**Supplementary Code 1**

Supplementary Code1. Python scripts for reproducing the Metabolic Spin-Glass (MSG) analysis and figure generation.

This supplementary code provides the Python scripts used to perform the analyses described in this study, including reaction-level transcriptomic preprocessing, construction of the Metabolic Spin-Glass (MSG) Hamiltonian, model-derived order parameter and frustration calculations, Gaussian mixture model clustering, pathway-level Shannon configuration-diversity analysis, statistical-potential landscape reconstruction, parameter-sweep analyses, Kaplan–Meier survival analysis, Cox proportional hazards regression, and generation of the main and supplementary figures. The scripts are intended to reproduce the analytical workflow described in the manuscript. Exact numerical reproduction requires the accompanying Supplementary Data files and identical software dependencies.

**Supplementary Code 2**

Supplementary Code 2. Computation of the patient-level model-derived order parameter and its associated transcriptome-adjusted survival benchmark analyses.

This supplementary code provides the Python script (‘Supplementary_Code_2_order_parameter_and_survival_benchmarks.py’) used to compute the patient-level model-derived order parameter (m) according to the formula given in Methods (all 235 reactions; σ □= sign of the reaction free energy, −ΔG □; z-scored per reaction across the full 497-patient cohort). The script also regenerates the associated survival benchmark analyses (Supplementary Tables 1–6), including the variance in m explained by transcriptomic principal components (PC1–PC5, PC1–PC10), multivariable Cox proportional-hazards models and likelihood-ratio tests for overall and disease-free survival, and four additional robustness analyses (PC1–PC20 attenuation, molecular-subtype adjustment, a bootstrap confidence interval for the change in concordance index, and five-fold cross-validated survival-model performance). Transcriptomic principal component scores are computed within this script by standard PCA on the same 235-reaction expression matrix used elsewhere in the manuscript. Exact numerical reproduction requires the accompanying Supplementary Data and the software versions specified in the script header; a built-in reproducibility check compares the output against the values reported in this manuscript.

**Supplementary Material 1–5.** Subtype-specific SC-MSG-optimized reaction configurations across the w□ –w□ parameter sweep.

Each file reports the HQA-optimized SC-MSG configuration for one molecular subtype (1, Gastric; 2, Inflammatory; 3, Intestinal; 4, Mixed stroma; 5, Stem-like), obtained by minimizing H^(p)(x) = Σ □ΔG□ x□ + w □Σ□ < b□□ x □x □+ w □Σ S □^(p)x□ across the same w□ –w□ grid used throughout the manuscript (0.1–2.1 in steps of 0.2 for each weight; 121 combinations), where S□ ^(p) is the subtype-averaged expression value of reaction i. For every (w□, w□) pair, the 235 reactions are split into s0_tokens (x□ = 0) and s1_tokens (x□ = 1) — the inactive and active sets returned by the D-Wave hybrid quantum annealer. These solutions are the source data for the parameter-sweep, optimization-landscape, solution-ensemble, and active-reaction-set Jaccard analyses in Figure 1B–E.

**Supplementary Material 6.** Code for constructing and solving the subtype-specific SC-MSG QUBO problems (Supplementary_material6_Code_subtype.ipynb).

This notebook generates Supplementary Material 1–5. For a chosen molecular subtype, it builds the SC-MSG Hamiltonian as a QUBO over the w□ –w□ grid from the ΔG, B-matrix, and subtype-averaged expression data, solves it with D-Wave’s hybrid quantum annealer (LeapHybridSampler), cross-checks the result against an equivalent integer-programming solution obtained with PuLP, and saves the resulting active and inactive reaction sets to solutions_<Subtype>.csv. The final two cells produce the parameter-sweep visualizations shown in Figure 1B–E.

## Reference

1. Warburg, O., On the Origin of Cancer Cells. Science, 1956. 123(3191): p. 309–314.

2. Liberti, M.V. and J.W. Locasale, The Warburg Effect: How Does it Benefit Cancer Cells? Trends in Biochemical Sciences, 2016. 41(3): p. 211–218.

3. Hanahan, D. and Robert A. Weinberg, Hallmarks of Cancer: The Next Generation. Cell, 2011. 144(5): p. 646–674.

4. Vander Heiden, M.G., L.C. Cantley, and C.B. Thompson, Understanding the Warburg Effect: The Metabolic Requirements of Cell Proliferation. Science, 2009. 324(5930): p. 1029–1033.

5. Lunt, S.Y. and M.G. Vander Heiden, Aerobic Glycolysis: Meeting the Metabolic Requirements of Cell Proliferation. Annual Review of Cell and Developmental Biology, 2011. 27(1): p. 441–464.

6. Pavlova, Natalya N. and Craig B. Thompson, The Emerging Hallmarks of Cancer Metabolism. Cell Metabolism, 2016. 23(1): p. 27–47.

7. DeBerardinis, R.J. and N.S. Chandel, Fundamentals of cancer metabolism. Science Advances, 2016. 2(5).

8. Badur, M.G. and C.M. Metallo, Reverse engineering the cancer metabolic network using flux analysis to understand drivers of human disease. Metabolic Engineering, 2018. 45: p. 95–108.

9. Henry, C.S., L.J. Broadbelt, and V. Hatzimanikatis, Thermodynamics-Based Metabolic Flux Analysis. Biophysical Journal, 2007. 92(5): p. 1792–1805.

10. Xiao, W., et al., NAD(H) and NADP(H) Redox Couples and Cellular Energy Metabolism. Antioxidants & Redox Signaling, 2018. 28(3): p. 251–272.

11. Hamilton, Joshua J., V. Dwivedi, and Jennifer L. Reed, Quantitative Assessment of Thermodynamic Constraints on the Solution Space of Genome-Scale Metabolic Models. Biophysical Journal, 2013. 105(2): p. 512–522.

12. Zerfaß, C., M. Asally, and O.S. Soyer, Interrogating metabolism as an electron flow system. Current Opinion in Systems Biology, 2019. 13: p. 59–67.

13. Ataman, M. and V. Hatzimanikatis, Heading in the right direction: thermodynamics-based network analysis and pathway engineering. Current Opinion in Biotechnology, 2015. 36: p. 176–182.

14. Dai, Z. and J.W. Locasale, Thermodynamic constraints on the regulation of metabolic fluxes. Journal of Biological Chemistry, 2018. 293(51): p. 19725–19739.

15. Bekiaris, P.S. and S. Klamt, Network-wide thermodynamic constraints shape NAD(P)H cofactor specificity of biochemical reactions. Nature Communications, 2023. 14(1).

16. Sung, J.Y. and J.H. Cheong, Intercellular communications and metabolic reprogramming as new predictive markers for immunotherapy responses in gastric cancer. Cancer Communications, 2022. 42(6): p. 572–575.

17. Sung, J.-Y. and J.-H. Cheong, Pan-Cancer Analysis Reveals Distinct Metabolic Reprogramming in Different Epithelial–Mesenchymal Transition Activity States. Cancers, 2021. 13(8).

18. Sung, J.Y., et al., Metabolic subtype reveals potential therapeutic vulnerability in acute promyelocytic leukaemia. Clinical and Translational Medicine, 2022. 12(7).

19. Cheong, J.-H., et al., Predictive test for chemotherapy response in resectable gastric cancer: a multi-cohort, retrospective analysis. The Lancet Oncology, 2018. 19(5): p. 629–638.

20. Sung, J.Y. and J.H. Cheong, Gene signature related to cancer stem cells and fibroblasts of stem-like gastric cancer predicts immunotherapy response. Clinical and Translational Medicine, 2023. 13(8).

21. Sung, J.Y. and J.H. Cheong, Prognosis-related gene signature is enriched in cancer-associated fibroblasts in the stem-like subtype of gastric cancer. Clinical and Translational Medicine, 2022. 12(6).

22. Sung, J.-Y. and J.-H. Cheong, The Matrisome Is Associated with Metabolic Reprograming in Stem-like Phenotypes of Gastric Cancer. Cancers, 2022. 14(6).

23. Sung, J.Y. and J.H. Cheong, Alternative lengthening of telomeres is mechanistically linked to potential therapeutic vulnerability in the stem-like subtype of gastric cancer. Clinical and Translational Medicine, 2021. 11(9).

24. Sung, J.Y., et al., A subtype of cancer-associated fibroblast expressing syndecan-2 (SDC2) predicts survival and immune checkpoint inhibitor response in gastric cancer. Clinical and Translational Medicine, 2024. 14(12).

25. Sung, J.-Y., J.-H. Cheong, and E.T. Kim, Spatially resolved endothelial signaling via nampt-itga5 drives immune evasion in stem-like gastric cancer. Cancer Immunology, Immunotherapy, 2025. 74(10).

26. Sung, J.-Y. and J.-H. Cheong, Quantum thermodynamics with information and coherence. Quantum Information Processing, 2026. 25(6).

27. Flamholz, A., et al., eQuilibrator--the biochemical thermodynamics calculator. Nucleic Acids Research, 2011. 40(D1): p. D770–D775.

28. Frades, I., et al., Genome Scale Modeling to Study the Metabolic Competition between Cells in the Tumor Microenvironment. Cancers, 2021. 13(18).

29. Banerji, C.R.S., et al., Cellular network entropy as the energy potential in Waddington’s differentiation landscape. Scientific Reports, 2013. 3(1).

30. Kim, S., et al., Quantum annealing for combinatorial optimization: a benchmarking study. npj Quantum Information, 2025. 11(1).

31. Rietman, E.A., et al., Thermodynamic measures of cancer: Gibbs free energy and entropy of protein–protein interactions. Journal of Biological Physics, 2016. 42(3): p. 339–350.

32. Fan, W., et al., Unraveling principles of thermodynamics for genome-scale metabolic networks using graph neural networks. Cell Systems, 2025. 16(10).

33. Yizhak, K., et al., Modeling cancer metabolism on a genome scale. Molecular Systems Biology, 2015. 11(6).

34. Wang, J., et al., Spatial Metabolomics Identifies Distinct Tumor-Specific Subtypes in Gastric Cancer Patients. Clinical Cancer Research, 2022. 28(13): p. 2865–2877.

35. Chen, H., et al., Molecular characterization and clinical relevance of metabolic signature subtypes in gastric cancer. Cell Reports, 2024. 43(7).

