## Supplementary figures and images for "A Spin-Glass Metabolic Hamiltonian optimized by Quantum Annealing Reveals Configurational Regimes of Cancer Metabolism"

### supplementary figure 1

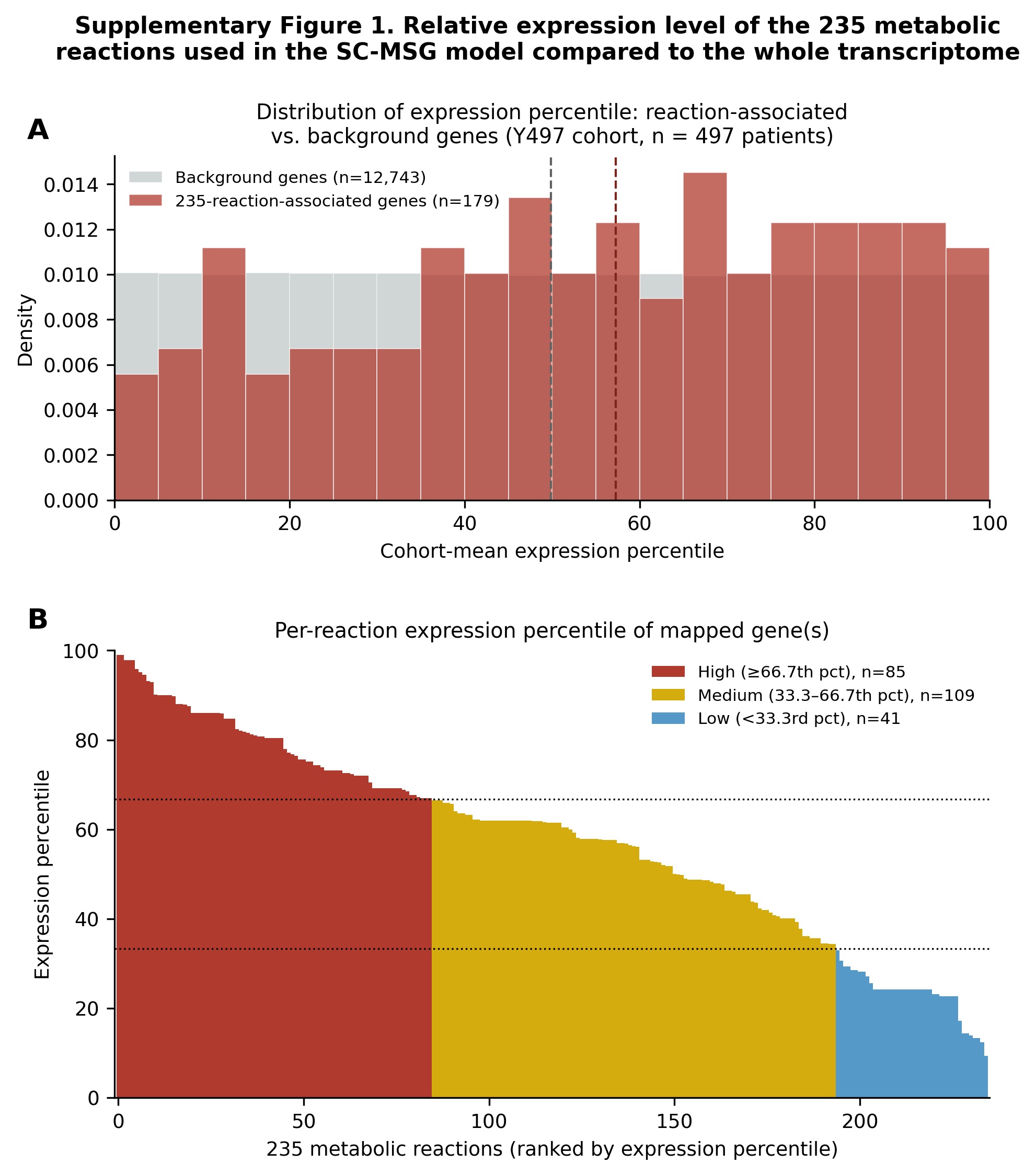

### supplementary figure 2

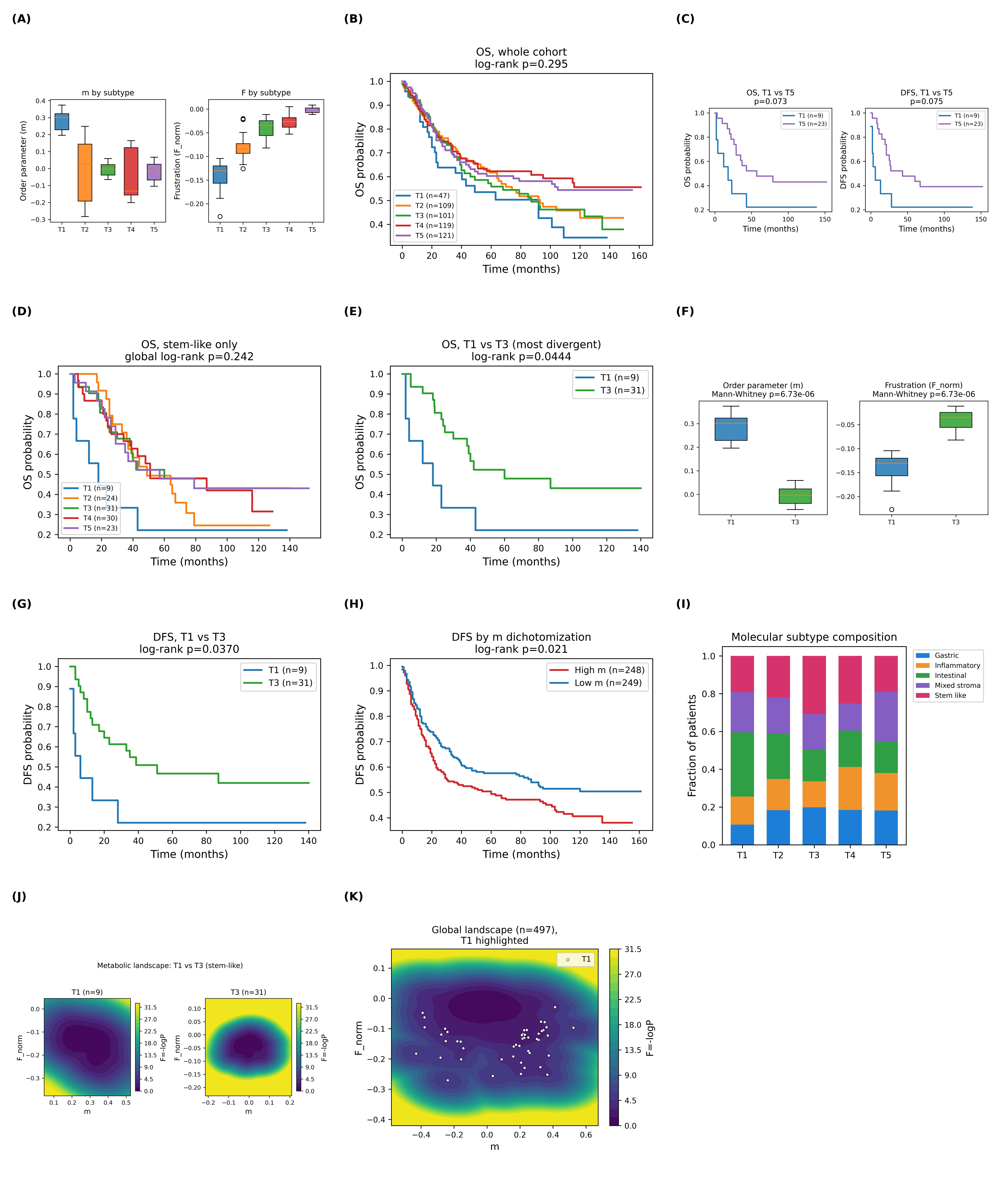

### supplementary figure 3

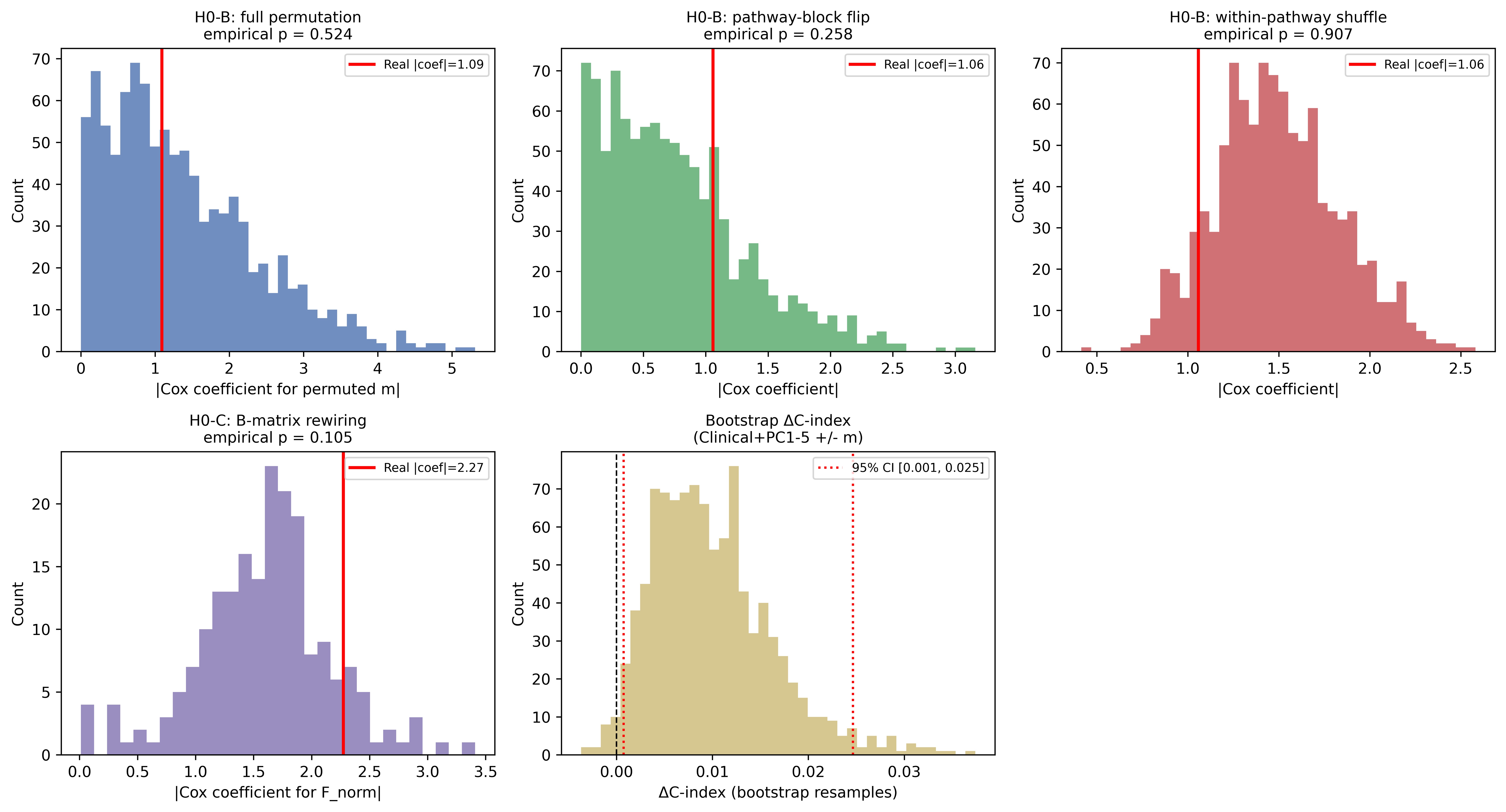

### supplementary figure 4

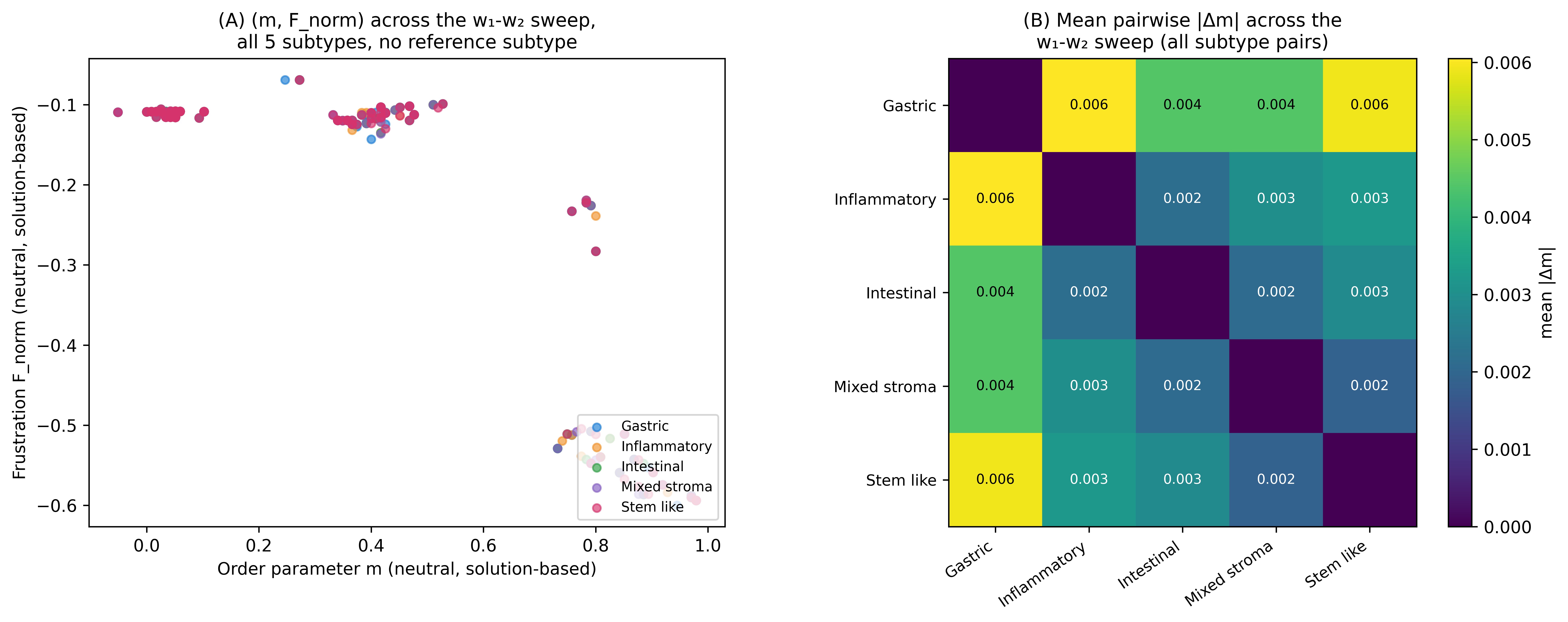

### supplementary figure 5

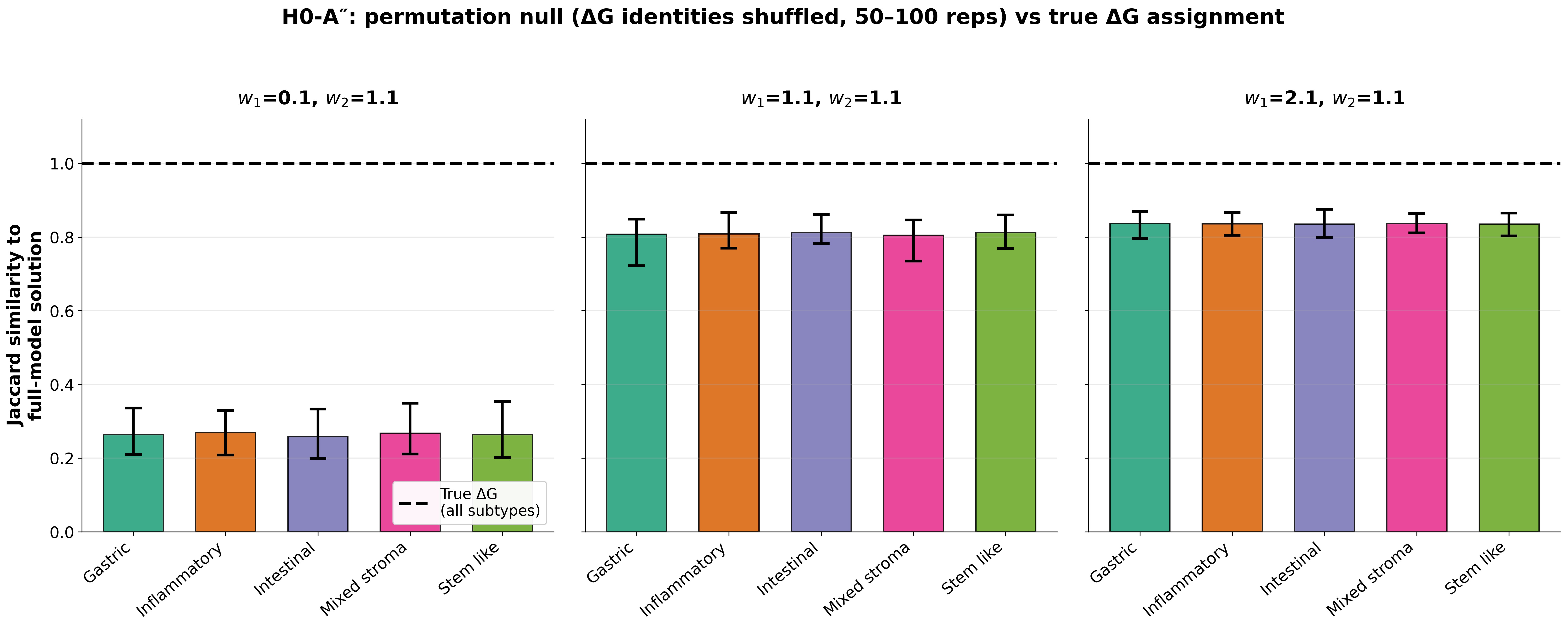
