## supplementary material 6 for "A Spin-Glass Metabolic Hamiltonian optimized by Quantum Annealing Reveals Configurational Regimes of Cancer Metabolism": Supplementary material6_Code_subtype.ipynb

```
# %% [code]
from collections import defaultdict
from dwave.system import LeapHybridSampler
import itertools
import random
import networkx as nx
import matplotlib.pyplot as plt
import pandas as pd
import numpy as np
import csv
import pulp

# %% [code]
target="Gastric" # Subtype name ex) Stem like/Mixed stroma/Intestinal/Inflammatory/Gastric

# DataSet Import Code

bdf = pd.read_csv('b_matrix_human_235.csv') #b_matrix import
bm = bdf.iloc[0:, 1:].astype(float).values
n_b = bm.shape[0]

gdf = pd.read_csv('Human_reactions_235_with_deltaG_equation.csv')  #dG_matrix import
gm = gdf["dG"].astype(float).values
n_g = gm.shape[0]

cdf = pd.read_csv("y497_clinic.txt", sep="\t")  # clinical data import
ids_by_type = { # Dictionary collecting Sample IDs by TYPE
    t: g["Sample"].tolist()
    for t, g in cdf.groupby("TYPE")
}
reac = pd.read_csv("Y497_reaction235_patient_matrix_mean.csv")  # S_matrix import
reac = reac.set_index("Reaction")
group_means = {}
for name, id_list in ids_by_type.items():  # ids_by_type: TYPE -> list of IDs
    cols = [c for c in id_list if c in reac.columns]
    group_means[name] = reac[cols].mean(axis=1)
S_sub=list(group_means[target])
n_s = len(S_sub)


# %% [code]
# HQA solver (Quantum), D-Wave API prior registration required

sol_list=list() # List to store solutions
for w_j in np.linspace(0.1,0.3,2): # w_j: [0.1, 2.1], 11 points specified
    for w_s in np.linspace(0.1,0.3,2): # w_s: [0.1, 2.1], 11 points specified
        w_j=float(w_j)
        w_s=float(w_s)
        Q = defaultdict(int)

        for i in range(n_b):
            for j in range(i,n_b):
                if bm[i,j]!=0:
                    Q[(i+1,j+1)]+= w_j*float(bm[i,j]) # b_ij Quadratic Term


        for i in range(n_g):
            Q[(i+1,i+1)]+= float(gm[i]) # delta G Linear Term


        for i in range(n_s):
            Q[(i+1,i+1)]+= w_s*float(S_sub[i]) # Personal S Linear Term

        sampler = LeapHybridSampler()           
        response_qubo = sampler.sample_qubo(Q)

        for sample, E in response_qubo.data(fields=['sample','energy']): 
            S0 = [k for k,v in response_qubo.first.sample.items() if v == 0] # Set of reactions with X=0 (inactive reactions)
            S1 = [k for k,v in response_qubo.first.sample.items() if v == 1] # Set of reactions with X=1 (active reactions)

        sol_list.append([[w_j,w_s],S0,S1]) # Store parameter combination, S0 list, and S1 list

# %% [code]
# MIP solver (Classical)

def solve_qubo_pulp(Q, n_nodes): # Function that converts a QUBO problem into a MIP problem and solves it
    prob = pulp.LpProblem("QUBO_Linearized", pulp.LpMinimize)
    x = [pulp.LpVariable(f"x{i}", cat="Binary") for i in range(n_nodes)]
    
    # Auxiliary variables z_ij = x_i * x_j: convert quadratic terms into linear terms
    z = {}
    for (i,j), val in Q.items():
        if i < j:
            z[(i,j)] = pulp.LpVariable(f"z{i}_{j}", 0, 1)
            prob += z[(i,j)] >= x[i-1] + x[j-1] - 1 
            prob += z[(i,j)] <= x[i-1]
            prob += z[(i,j)] <= x[j-1]
    
    # Linear objective function
    obj = sum(Q[(i+1,i+1)] * x[i] for i in range(n_nodes) if (i+1,i+1) in Q)
    obj += sum(val * z[(i,j)] for (i,j), val in Q.items() if i < j)
    prob += obj
    
    status = prob.solve(pulp.PULP_CBC_CMD(msg=1, timeLimit=300))
    
    return [int(v.value()) for v in x if v.value() is not None], pulp.value(prob.objective)

sol_list=list() # List to store solutions
for w_j in np.linspace(0.1,2.1,11):   # w_j: [0.1, 2.1], 11 points specified
    for w_s in np.linspace(0.1,2.1,11): # w_s: [0.1, 2.1], 11 points specified
        w_j=float(w_j)
        w_s=float(w_s)
        S0=list()
        S1=list()
        Q = defaultdict(int)

        for i in range(n_b):
            for j in range(i,n_b):
                if bm[i,j]!=0:
                    Q[(i+1,j+1)]+= w_j*float(bm[i,j]) # b_ij Quadratic Term


        for i in range(n_g):
            Q[(i+1,i+1)]+= float(gm[i]) # delta G Linear Term


        for i in range(n_s):
            Q[(i+1,i+1)]+= w_s*float(S_sub[i]) # Personal S Linear Term

        n_nodes = max(max(k) for k in Q.keys())  # Automatic determination
        solution, energy = solve_qubo_pulp(Q, n_nodes)

        for i in range(len(solution)):
            if solution[i]==0:
                S0.append(i+1) # Set of reactions with X=0 (inactive reactions)
            else :     
                S1.append(i+1) # Set of reactions with X=1 (active reactions)
        sol_list.append([[w_j,w_s],S0,S1])  # Store parameter combination, S0 list, and S1 list

# %% [code]
# Save raw data file: stored in the order
# (patient name, w_j = w1 value, w_s = w2 value, S0 list, S1 list)

def format_float_short(x, ndigits=10): # Float → clean string: 0.300000 → '0.3', 1.000000 → '1'
    x_rounded = round(x, ndigits)
    s = f"{x_rounded:.{ndigits}f}"
    s = s.rstrip('0').rstrip('.')  # Remove trailing zeros and decimal point
    return s
    return s

def clean_s0_tokens(s0_list):
    """Convert s0 list → space-separated string: [3, 4, 5] → '3 4 5'"""
    if isinstance(s0_list, (list, tuple)):
        return ' '.join(map(str, s0_list))
    else:
        return str(s0_list)

target_re=target.replace(' ','_')

# sol_list is assumed to have the form: [((w1, w2), s0_list, s1_list), ...]
with open('solutions_%s.csv' %target_re, 'w', newline='', encoding='utf-8') as f:
    writer = csv.writer(f)
    writer.writerow(['w1', 'w2', 's0_tokens', 's1_tokens'])  # Header

    for (w_pair, s0_raw, s1_raw) in sol_list:
        w1, w2 = float(w_pair[0]), float(w_pair[1])

        # Clean formatting for w1, w2
        w1_str = format_float_short(w1)  # e.g.: '0.3'
        w2_str = format_float_short(w2)

        # Convert s0 tokens to a clean space-separated format
        s0_str = clean_s0_tokens(s0_raw)  # e.g.: '3 4 5 6 9 12 ...'
        s1_str = clean_s0_tokens(s1_raw)
        
        writer.writerow([w1_str, w2_str, s0_str,s1_str])

print("Raw data CSV saved: solutions_%s.csv" % target_re)

# %% [code]
# Visualization of results for a single Type
group="Mixed stroma" # Set subtype name


group_re=group.replace(" ","_")
df_sub = pd.read_csv('solutions_%s.csv' %group_re) # Raw Data import

df_sub["pattern"] = df_sub["s0_tokens"].astype(str).str.strip()

all_patterns = sorted(set(df_sub["pattern"].unique()))
style_map = {}  # Pattern → (color, marker) mapping

color_list = plt.cm.tab10(np.linspace(0, 1, 10))
marker_list = ["o", "s", "^",'d','8', "D",'X', "v", "*", "p", "P", "h", "H","<",">"]
style_combinations = list(itertools.product(color_list, marker_list))

random.seed(42)  # Fixed seed
random.shuffle(style_combinations)

# Assign fixed styles to all patterns
for i, pattern in enumerate(all_patterns):
    style_map[pattern] = style_combinations[i % len(style_combinations)]

# ===== Create plot =====
fig, (ax1) = plt.subplots(1, 1, figsize=(20, 20))

# HQA plot (using a common style map)
for pattern, (c, m) in style_map.items():
    sub1 = df_sub[df_sub["pattern"] == pattern]
    if not sub1.empty:
        ax1.scatter(
            sub1["w1"], sub1["w2"],
            c=[c], marker=m, label=pattern,
            edgecolors="k", s=1450, alpha=0.8
        )


# Settings
ax1.set_xlabel("w1", fontweight='bold'); ax1.set_ylabel("w2", fontweight='bold')
ax1.set_title("%s Group" %group, fontsize=16, fontweight='bold')


# Legend (simplified)
# ===== Copy only this part =====
dummy_handles = []
dummy_labels = []

# Create dummy handles for all patterns that actually appear
all_actual_patterns = sorted(set([
    p for p in all_patterns if not df_sub[df_sub["pattern"] == p].empty
]))

for pattern in all_actual_patterns:
    c, m = style_map[pattern]
    # Create handles only (with empty data)
    handle = ax1.scatter([], [], c=[c], marker=m, s=150, 
                        edgecolors="k", alpha=0.8, label=pattern)
    dummy_handles.append(handle)
    dummy_labels.append(pattern)

# Display legend
fig.legend(dummy_handles, dummy_labels, 
           loc='lower center', bbox_to_anchor=(0.4, 0.3),
           ncol=6, fontsize=10)


plt.tight_layout()
plt.subplots_adjust(bottom=0.5)
plt.savefig('scatter_patterns_%s.png' %group_re , dpi=300, bbox_inches='tight')
plt.show()

# %% [code]
# Visualization of results for 5 Types
df1 = pd.read_csv('solutions_Stem_like.csv') # Raw Data import
df2 = pd.read_csv('solutions_Inflammatory.csv' ) 
df3 = pd.read_csv('solutions_Gastric.csv' ) 
df4 = pd.read_csv('solutions_Intestinal.csv' ) 
df5 = pd.read_csv('solutions_Mixed_stroma.csv' ) 


# Create pattern column
df1["pattern"] = df1["s0_tokens"].astype(str).str.strip()
df2["pattern"] = df2["s0_tokens"].astype(str).str.strip()
df3["pattern"] = df3["s0_tokens"].astype(str).str.strip()
df4["pattern"] = df4["s0_tokens"].astype(str).str.strip()
df5["pattern"] = df5["s0_tokens"].astype(str).str.strip()


all_patterns = sorted(set(df1["pattern"].unique()) | set(df2["pattern"].unique()) | set(df3["pattern"].unique())|set(df4["pattern"].unique())|set(df5["pattern"].unique()))
style_map = {}  # Pattern → (color, marker) mapping

color_list = plt.cm.tab10(np.linspace(0, 1, 10))
marker_list = ["o", "s", "^",'d','8', "D",'X', "v", "*", "p", "P", "h", "H","<",">"]
style_combinations = list(itertools.product(color_list, marker_list))

random.seed(42)  # Fixed seed
random.shuffle(style_combinations)

# Assign fixed styles to all patterns
for i, pattern in enumerate(all_patterns):
    style_map[pattern] = style_combinations[i % len(style_combinations)]

# ===== Create 5 subplots =====
fig, (ax1, ax2, ax3, ax4, ax5) = plt.subplots(1, 5, figsize=(80, 20))

# HQA plot (using a common style map)
for pattern, (c, m) in style_map.items():
    sub1 = df1[df1["pattern"] == pattern]
    if not sub1.empty:
        ax1.scatter(
            sub1["w1"], sub1["w2"],
            c=[c], marker=m, label=pattern,
            edgecolors="k", s=1450, alpha=0.8
        )

# HQA2 plot (using the same style map!)
for pattern, (c, m) in style_map.items():
    sub2 = df2[df2["pattern"] == pattern]
    if not sub2.empty:
        ax2.scatter(
            sub2["w1"], sub2["w2"],
            c=[c], marker=m, label=pattern,
            edgecolors="k", s=1450, alpha=0.8
        )
for pattern, (c, m) in style_map.items():
    sub3 = df3[df3["pattern"] == pattern]
    if not sub3.empty:
        ax3.scatter(
            sub3["w1"], sub3["w2"],
            c=[c], marker=m, label=pattern,
            edgecolors="k", s=1450, alpha=0.8
        )
for pattern, (c, m) in style_map.items():
    sub4 = df4[df4["pattern"] == pattern]
    if not sub4.empty:
        ax4.scatter(
            sub4["w1"], sub4["w2"],
            c=[c], marker=m, label=pattern,
            edgecolors="k", s=1450, alpha=0.8
        )
for pattern, (c, m) in style_map.items():
    sub5 = df5[df5["pattern"] == pattern]
    if not sub5.empty:
        ax5.scatter(
            sub5["w1"], sub5["w2"],
            c=[c], marker=m, label=pattern,
            edgecolors="k", s=1450, alpha=0.8
        )

# Settings
ax1.set_xlabel("w1", fontweight='bold'); ax1.set_ylabel("w2", fontweight='bold')
ax1.set_title("Stem like", fontsize=16, fontweight='bold')
ax2.set_xlabel("w1", fontweight='bold')
ax2.set_title("Inflammatory", fontsize=16, fontweight='bold')
ax3.set_xlabel("w1", fontweight='bold')
ax3.set_title("Gastric", fontsize=16, fontweight='bold')
ax4.set_xlabel("w1", fontweight='bold')
ax4.set_title("Intestinal", fontsize=16, fontweight='bold')
ax5.set_xlabel("w1", fontweight='bold')
ax5.set_title("Mixed stroma", fontsize=16, fontweight='bold')


# Legend (simplified)
# ===== Copy only this part =====
dummy_handles = []
dummy_labels = []

# Create dummy handles for all patterns that actually appear
all_actual_patterns = sorted(set([
    p for p in all_patterns if not df1[df1["pattern"] == p].empty or 
    not df2[df2["pattern"] == p].empty or 
    not df3[df3["pattern"] == p].empty or 
    not df4[df4["pattern"] == p].empty or 
    not df5[df5["pattern"] == p].empty
]))

for pattern in all_actual_patterns:
    c, m = style_map[pattern]
    # Create handles only (with empty data)
    handle = ax1.scatter([], [], c=[c], marker=m, s=150, 
                        edgecolors="k", alpha=0.8, label=pattern)
    dummy_handles.append(handle)
    dummy_labels.append(pattern)

# Display legend
fig.legend(dummy_handles, dummy_labels, 
           loc='lower center', bbox_to_anchor=(0.4, 0.2),
           ncol=6, fontsize=10)


plt.tight_layout()
plt.subplots_adjust(bottom=0.5)
plt.savefig('scatter_patterns_all.png', dpi=300, bbox_inches='tight')
plt.show()
```
