## supplementary code 1 for "A Spin-Glass Metabolic Hamiltonian optimized by Quantum Annealing Reveals Configurational Regimes of Cancer Metabolism": Supplementary_Code_1_SC_MSG_analysis_and_figures.py

```
# Supplementary Code for:
# "A Spin-Glass Metabolic Hamiltonian optimized by Quantum Annealing
#  Reveals Configurational Regimes of Cancer Metabolism"
#
# Purpose
# -------
# Reproducible analysis and figure-generation template covering:
#   1. Input preprocessing and reaction-level expression construction
#   2. PCA and collective coordinates
#   3. Gaussian mixture model clustering with BIC model selection
#   4. Model-derived order parameter and frustration score
#   5. Pathway-level Shannon configuration-diversity analysis
#   6. Statistical-potential / landscape plots
#   7. Parameter-sweep and phase-map plots
#   8. Kaplan-Meier survival analysis and Cox regression
#   9. Figure-panel cross-reference map
#
# IMPORTANT
# ---------
# This script is structured to reproduce the analytical procedures described in
# the revised manuscript. Exact numerical reproduction requires the final
# Supplementary Data files used in the study, with column names matching the
# CONFIG section below.
#
# Recommended Python version: >= 3.10
#
# Required packages:
# pip install numpy pandas scipy scikit-learn matplotlib lifelines statsmodels openpyxl
#
# Optional packages:
# pip install seaborn dwave-system dimod
#
# =============================================================================
# 0. IMPORTS AND CONFIGURATION
# =============================================================================

from __future__ import annotations

import json
import math
import warnings
from pathlib import Path
from typing import Dict, Iterable, List, Optional, Sequence, Tuple

import numpy as np
import pandas as pd
import matplotlib.pyplot as plt

from scipy import stats
from scipy.ndimage import gaussian_filter
from scipy.stats import gaussian_kde, entropy, pearsonr, kruskal, mannwhitneyu

from sklearn.decomposition import PCA
from sklearn.mixture import GaussianMixture
from sklearn.preprocessing import StandardScaler, MinMaxScaler
from sklearn.metrics import silhouette_score

from lifelines import KaplanMeierFitter, CoxPHFitter
from lifelines.statistics import logrank_test, multivariate_logrank_test

warnings.filterwarnings("ignore", category=RuntimeWarning)

RANDOM_SEED = 42
np.random.seed(RANDOM_SEED)

BASE_DIR = Path(".")
DATA_DIR = BASE_DIR / "data"
RESULT_DIR = BASE_DIR / "results"
FIGURE_DIR = RESULT_DIR / "figures"
TABLE_DIR = RESULT_DIR / "tables"

for directory in (RESULT_DIR, FIGURE_DIR, TABLE_DIR):
    directory.mkdir(parents=True, exist_ok=True)

# Expected files
REACTION_EXPRESSION_FILE = DATA_DIR / "Supplementary_Data_2_reaction_expression.csv"
REACTION_ANNOTATION_FILE = DATA_DIR / "Supplementary_Data_1_reaction_annotation.csv"
B_MATRIX_FILE = DATA_DIR / "Supplementary_Data_3_B_matrix.csv"
CLINICAL_FILE = DATA_DIR / "clinical_data.csv"
OPTIMIZED_STATE_FILE = DATA_DIR / "optimized_binary_states.csv"
PATHWAY_MAP_FILE = DATA_DIR / "reaction_pathway_map.csv"
PARAMETER_SWEEP_FILE = DATA_DIR / "w1_w2_parameter_sweep.csv"

# Expected key columns
PATIENT_COL = "patient_id"
MOLECULAR_SUBTYPE_COL = "molecular_subtype"
SC_MSG_SUBTYPE_COL = "sc_msg_subtype"

OS_TIME_COL = "OS_time"
OS_EVENT_COL = "OS_event"
DFS_TIME_COL = "DFS_time"
DFS_EVENT_COL = "DFS_event"

AGE_COL = "age"
SEX_COL = "sex"
STAGE_COL = "pathological_stage"

REACTION_ID_COL = "reaction_id"
DG_COL = "delta_g"
PATHWAY_COL = "pathway"

EPSILON = 1e-12


# =============================================================================
# 1. UTILITY FUNCTIONS
# =============================================================================

def read_table(path: Path, index_col: Optional[int | str] = None) -> pd.DataFrame:
    """Read CSV, TSV, or Excel files with basic validation."""
    if not path.exists():
        raise FileNotFoundError(
            f"Required input file not found: {path}\n"
            "Place the final Supplementary Data file in the data/ directory "
            "or change the path in the CONFIG section."
        )

    suffix = path.suffix.lower()
    if suffix == ".csv":
        return pd.read_csv(path, index_col=index_col)
    if suffix in {".tsv", ".txt"}:
        return pd.read_csv(path, sep="\t", index_col=index_col)
    if suffix in {".xlsx", ".xls"}:
        return pd.read_excel(path, index_col=index_col)
    raise ValueError(f"Unsupported file format: {suffix}")


def save_figure(fig: plt.Figure, filename: str, dpi: int = 300) -> None:
    """Save a publication-quality figure."""
    fig.tight_layout()
    fig.savefig(FIGURE_DIR / filename, dpi=dpi, bbox_inches="tight")
    plt.close(fig)


def zscore_columns(df: pd.DataFrame) -> pd.DataFrame:
    """Z-score each feature across patients."""
    scaler = StandardScaler()
    values = scaler.fit_transform(df.astype(float))
    return pd.DataFrame(values, index=df.index, columns=df.columns)


def minmax_01(values: np.ndarray) -> np.ndarray:
    """Rescale a one- or two-dimensional array to [0, 1]."""
    scaler = MinMaxScaler(feature_range=(0, 1))
    arr = np.asarray(values)
    if arr.ndim == 1:
        arr = arr.reshape(-1, 1)
    scaled = scaler.fit_transform(arr)
    return scaled.ravel() if scaled.shape[1] == 1 else scaled


def safe_log_probability(probability: np.ndarray, epsilon: float = EPSILON) -> np.ndarray:
    """Calculate -log(P + epsilon), then shift the minimum to zero."""
    potential = -np.log(np.asarray(probability, dtype=float) + epsilon)
    finite = np.isfinite(potential)
    if finite.any():
        potential = potential - np.nanmin(potential[finite])
    return potential


# =============================================================================
# 2. INPUT PREPROCESSING AND REACTION-LEVEL EXPRESSION
# Corresponding Methods:
#   - Input data and preprocessing
#   - Molecular subtype-specific transcriptomic embedding
# Relevant panels:
#   - Figure 1A
#   - Figure 3A-G
#   - Figure 4A-L
# =============================================================================

def load_reaction_expression() -> pd.DataFrame:
    """
    Load patient-by-reaction expression matrix.

    Required format:
      rows    = patients
      columns = reaction IDs
    The first column may be patient_id.
    """
    df = read_table(REACTION_EXPRESSION_FILE)

    if PATIENT_COL in df.columns:
        df = df.set_index(PATIENT_COL)

    numeric = df.apply(pd.to_numeric, errors="coerce")
    numeric = numeric.dropna(axis=1, how="all")
    numeric = numeric.fillna(numeric.median(numeric_only=True))
    return numeric


def aggregate_gene_expression_to_reactions(
    gene_expression: pd.DataFrame,
    gene_reaction_map: pd.DataFrame,
    gene_col: str = "gene_symbol",
    reaction_col: str = REACTION_ID_COL,
    aggregation: str = "mean",
) -> pd.DataFrame:
    """
    Construct patient-by-reaction expression scores from a patient-by-gene matrix.

    The manuscript states that expression values of multiple genes mapped to one
    reaction were averaged. This function implements that procedure.
    """
    required = {gene_col, reaction_col}
    missing = required.difference(gene_reaction_map.columns)
    if missing:
        raise ValueError(f"Missing mapping columns: {sorted(missing)}")

    reaction_scores = {}
    for reaction_id, group in gene_reaction_map.groupby(reaction_col):
        genes = [g for g in group[gene_col].dropna().astype(str) if g in gene_expression.columns]
        if not genes:
            continue

        sub = gene_expression[genes].astype(float)
        if aggregation == "mean":
            reaction_scores[reaction_id] = sub.mean(axis=1)
        elif aggregation == "sum":
            reaction_scores[reaction_id] = sub.sum(axis=1)
        elif aggregation == "max":
            reaction_scores[reaction_id] = sub.max(axis=1)
        else:
            raise ValueError("aggregation must be 'mean', 'sum', or 'max'")

    result = pd.DataFrame(reaction_scores, index=gene_expression.index)
    result.index.name = PATIENT_COL
    return result


def subtype_transcriptomic_field(
    reaction_expression: pd.DataFrame,
    metadata: pd.DataFrame,
    subtype_col: str = MOLECULAR_SUBTYPE_COL,
) -> pd.DataFrame:
    """
    Calculate subtype-specific transcriptomic field:
        S_i^(k) = -log(1 + mean_expression_i^(k))
    """
    aligned = metadata.loc[reaction_expression.index]
    output = {}

    for subtype, patient_ids in aligned.groupby(subtype_col).groups.items():
        mean_expression = reaction_expression.loc[list(patient_ids)].mean(axis=0)
        output[subtype] = -np.log1p(np.clip(mean_expression, a_min=0, a_max=None))

    return pd.DataFrame(output).T


# =============================================================================
# 3. PCA AND COLLECTIVE COORDINATES
# Corresponding Methods:
#   - Construction of collective coordinates (m1, m2)
#   - Identification of dominant collective axes
#   - Transcriptomic PCA benchmark
# Relevant panels:
#   - Figure 2A-C
#   - Supplementary PCA-adjusted survival analyses
# =============================================================================

def run_pca(
    feature_matrix: pd.DataFrame,
    n_components: int = 10,
    standardize: bool = True,
) -> Tuple[pd.DataFrame, pd.DataFrame, pd.DataFrame]:
    """
    Run PCA with patients as observations and reactions/pathways as features.

    Returns:
      scores            patient x PC
      loadings          feature x PC
      explained_variance table
    """
    matrix = feature_matrix.copy().astype(float)

    if standardize:
        values = StandardScaler().fit_transform(matrix)
    else:
        values = matrix.to_numpy()

    max_components = min(n_components, values.shape[0], values.shape[1])
    pca = PCA(n_components=max_components, random_state=RANDOM_SEED)
    scores_array = pca.fit_transform(values)

    pc_names = [f"PC{i+1}" for i in range(max_components)]
    scores = pd.DataFrame(scores_array, index=matrix.index, columns=pc_names)
    loadings = pd.DataFrame(pca.components_.T, index=matrix.columns, columns=pc_names)

    explained = pd.DataFrame(
        {
            "principal_component": pc_names,
            "explained_variance_ratio": pca.explained_variance_ratio_,
            "cumulative_explained_variance": np.cumsum(pca.explained_variance_ratio_),
        }
    )

    scores.to_csv(TABLE_DIR / "PCA_scores.csv")
    loadings.to_csv(TABLE_DIR / "PCA_loadings.csv")
    explained.to_csv(TABLE_DIR / "PCA_explained_variance.csv", index=False)

    return scores, loadings, explained


def plot_pca_scatter(
    scores: pd.DataFrame,
    metadata: pd.DataFrame,
    group_col: str,
    filename: str = "PCA_scatter.png",
) -> None:
    """Plot PC1 versus PC2 by molecular or SC-MSG-derived subtype."""
    data = scores.join(metadata[[group_col]], how="inner")

    fig, ax = plt.subplots(figsize=(7, 6))
    for group_name, group in data.groupby(group_col):
        ax.scatter(group["PC1"], group["PC2"], s=28, alpha=0.75, label=str(group_name))

    ax.set_xlabel("PC1")
    ax.set_ylabel("PC2")
    ax.set_title("Principal component analysis")
    ax.legend(frameon=False, fontsize=8)
    save_figure(fig, filename)


def construct_collective_coordinates_from_pathway_entropy(
    pathway_entropy_matrix: pd.DataFrame,
) -> pd.DataFrame:
    """
    Manuscript-compatible construction:
      1. Standardize pathway-level entropy values across patients.
      2. Apply PCA.
      3. Use PC1 and PC2 scores.
      4. Rescale each axis to [0, 1].
    """
    scores, _, _ = run_pca(pathway_entropy_matrix, n_components=2, standardize=True)
    coordinates = pd.DataFrame(index=scores.index)
    coordinates["m1"] = minmax_01(scores["PC1"].to_numpy())
    coordinates["m2"] = minmax_01(scores["PC2"].to_numpy())
    coordinates.to_csv(TABLE_DIR / "collective_coordinates_m1_m2.csv")
    return coordinates


# =============================================================================
# 4. GAUSSIAN MIXTURE MODEL CLUSTERING WITH BIC MODEL SELECTION
# Corresponding Methods:
#   - Definition of SC-MSG-derived subtypes
# Relevant panels:
#   - Figure 3K-L
#   - Figure 4A-L
# =============================================================================

def select_gmm_by_bic(
    feature_matrix: pd.DataFrame,
    min_components: int = 1,
    max_components: int = 10,
    covariance_types: Sequence[str] = ("full", "tied", "diag", "spherical"),
) -> Tuple[GaussianMixture, pd.DataFrame]:
    """
    Select a Gaussian mixture model using the minimum Bayesian information
    criterion (BIC).
    """
    X = feature_matrix.astype(float).to_numpy()
    rows = []
    best_model = None
    best_bic = np.inf

    for covariance_type in covariance_types:
        for n_components in range(min_components, max_components + 1):
            model = GaussianMixture(
                n_components=n_components,
                covariance_type=covariance_type,
                random_state=RANDOM_SEED,
                n_init=20,
                max_iter=1000,
                reg_covar=1e-6,
            )
            model.fit(X)
            bic = model.bic(X)
            aic = model.aic(X)

            labels = model.predict(X)
            silhouette = (
                silhouette_score(X, labels)
                if len(np.unique(labels)) > 1 and len(labels) > len(np.unique(labels))
                else np.nan
            )

            rows.append(
                {
                    "n_components": n_components,
                    "covariance_type": covariance_type,
                    "BIC": bic,
                    "AIC": aic,
                    "silhouette_score": silhouette,
                }
            )

            if bic < best_bic:
                best_bic = bic
                best_model = model

    bic_table = pd.DataFrame(rows).sort_values("BIC")
    bic_table.to_csv(TABLE_DIR / "GMM_BIC_model_selection.csv", index=False)

    if best_model is None:
        raise RuntimeError("No GMM model could be fitted.")

    return best_model, bic_table


def assign_sc_msg_subtypes(
    order_frustration: pd.DataFrame,
    n_components: Optional[int] = 5,
) -> pd.DataFrame:
    """
    Cluster patients in the (|m|, F_norm) space.

    When n_components=None, BIC chooses the number of clusters.
    When n_components=5, the output is labeled T1-T5 after sorting clusters by
    increasing normalized frustration and then increasing |m|.
    """
    required = {"abs_m", "F_norm"}
    missing = required.difference(order_frustration.columns)
    if missing:
        raise ValueError(f"Missing columns: {sorted(missing)}")

    X_df = order_frustration[["abs_m", "F_norm"]].copy()
    X = StandardScaler().fit_transform(X_df)

    if n_components is None:
        best_model, bic_table = select_gmm_by_bic(pd.DataFrame(X, index=X_df.index))
    else:
        best_model = GaussianMixture(
            n_components=n_components,
            covariance_type="full",
            random_state=RANDOM_SEED,
            n_init=50,
            max_iter=1000,
            reg_covar=1e-6,
        ).fit(X)
        _, bic_table = select_gmm_by_bic(
            pd.DataFrame(X, index=X_df.index),
            min_components=1,
            max_components=max(8, n_components),
        )

    raw_labels = best_model.predict(X)
    probabilities = best_model.predict_proba(X)

    output = X_df.copy()
    output["raw_cluster"] = raw_labels
    output["cluster_probability"] = probabilities.max(axis=1)

    centroids = output.groupby("raw_cluster")[["F_norm", "abs_m"]].mean()
    ordered_raw_clusters = (
        centroids.sort_values(["F_norm", "abs_m"], ascending=[True, True]).index.tolist()
    )
    label_map = {raw: f"T{i+1}" for i, raw in enumerate(ordered_raw_clusters)}
    output[SC_MSG_SUBTYPE_COL] = output["raw_cluster"].map(label_map)

    output.to_csv(TABLE_DIR / "sc_msg_subtype_assignments.csv")
    return output


def plot_gmm_bic(bic_table: pd.DataFrame, filename: str = "GMM_BIC_selection.png") -> None:
    """Plot BIC values across candidate GMM models."""
    fig, ax = plt.subplots(figsize=(8, 5))
    for covariance_type, group in bic_table.groupby("covariance_type"):
        group = group.sort_values("n_components")
        ax.plot(
            group["n_components"],
            group["BIC"],
            marker="o",
            label=covariance_type,
        )

    ax.set_xlabel("Number of Gaussian components")
    ax.set_ylabel("Bayesian information criterion")
    ax.set_title("GMM model selection")
    ax.legend(frameon=False)
    save_figure(fig, filename)


# =============================================================================
# 5. MODEL-DERIVED ORDER PARAMETER AND FRUSTRATION SCORE
# Corresponding Methods:
#   - Model-derived order parameter
#   - Patient-level extension of the order parameter
#   - Metabolic frustration index
# Relevant panels:
#   - Figure 3A-D, F-G
#   - Figure 4A, E-G, K-L
#
# NOTE (post-review correction): the order parameter formula below
# (sigma_r = sign(-dG_r), m_i = mean_r[sigma_r * z_ir] over all reactions)
# matches the Methods section as written and has been verified against the
# corrected Supplementary Table 6. The frustration index below
# (`calculate_frustration_index`) replaces an earlier formulation
# (`calculate_binary_frustration`, retained below for reference) that relied
# on patient-specific optimized binary states; the current definition uses
# only the continuous reaction-expression matrix, ΔG, and the B matrix, and
# is the definition reported in Figure 3D. See AUDIT_NOTES.md (accompanying
# the Figure 3 analysis package) for the full derivation and validation.
# =============================================================================

def calculate_thermodynamic_sign(delta_g: pd.Series) -> pd.Series:
    """
    Encode intrinsic thermodynamic direction.
    Exergonic reactions (Delta G < 0) are assigned +1.
    Endergonic reactions (Delta G > 0) are assigned -1.
    Zero values are assigned 0.
    """
    sigma = -np.sign(delta_g.astype(float))
    return pd.Series(sigma, index=delta_g.index, name="sigma")


def calculate_order_parameter(
    reaction_expression: pd.DataFrame,
    delta_g: pd.Series,
) -> pd.DataFrame:
    """
    Patient-level weighted model-derived order parameter:
        m_i = (1/N) * sum_r sigma_r * z_ir

    z_ir is the per-reaction z-score across patients.
    """
    common = reaction_expression.columns.intersection(delta_g.index)
    if len(common) == 0:
        raise ValueError("No overlapping reaction IDs between expression and Delta G.")

    z = zscore_columns(reaction_expression[common])
    sigma = calculate_thermodynamic_sign(delta_g.loc[common])

    contribution = z.mul(sigma, axis=1)
    m = contribution.mean(axis=1)

    output = pd.DataFrame({"m": m, "abs_m": m.abs()}, index=reaction_expression.index)
    output.to_csv(TABLE_DIR / "patient_order_parameter.csv")
    contribution.to_csv(TABLE_DIR / "reaction_level_order_contributions.csv")
    return output


def calculate_frustration_index(
    reaction_expression: pd.DataFrame,
    delta_g: pd.Series,
    B_matrix: pd.DataFrame,
) -> pd.DataFrame:
    """
    Metabolic frustration index used for Figure 3D (manuscript definition).

    For each reaction r and patient i, define a spin variable capturing
    whether the reaction's activation agrees with its thermodynamically
    preferred direction:
        s_ir = sigma_r * sign(z_ir)
    where sigma_r = sign(-dG_r) and z_ir is the per-reaction z-score.

    The frustration index is the B-matrix-weighted magnitude of
    energetically unfavorable ("frustrated") reaction pairs:
        F_i = sum_{r<r', B_rr' != 0}  |B_rr'| * 1[ B_rr' * s_ir * s_ir' > 0 ]

    This does not require patient-specific optimized binary states; it uses
    only the continuous reaction-expression matrix, the reaction free
    energies, and the reaction-pair coupling matrix (B), consistent with the
    order parameter m computed by `calculate_order_parameter`.

    Returns a DataFrame indexed by patient with columns:
        F_raw   : the raw frustration index above
        F_norm  : F_raw min-max normalized to [0, 1]
                  (F_norm is provided only for downstream steps, e.g.
                  `assign_sc_msg_subtypes`, that expect a bounded
                  scale; F_raw is the quantity reported in Figure 3D.)
    """
    common = reaction_expression.columns.intersection(delta_g.index)
    common = common.intersection(B_matrix.index).intersection(B_matrix.columns)
    if len(common) == 0:
        raise ValueError("No overlapping reaction IDs between expression, Delta G, and B.")

    z = zscore_columns(reaction_expression[common])
    sigma = calculate_thermodynamic_sign(delta_g.loc[common])

    B = B_matrix.loc[common, common].astype(float).to_numpy()
    B = (B + B.T) / 2.0
    np.fill_diagonal(B, 0.0)

    n_rxn = len(common)
    iu, ju = np.triu_indices(n_rxn, k=1)
    Bij = B[iu, ju]
    nonzero = Bij != 0
    iu, ju, Bij = iu[nonzero], ju[nonzero], Bij[nonzero]

    # s_ir = sigma_r * sign(z_ir), shape (n_patients, n_reactions)
    S = np.sign(z.to_numpy()) * sigma.to_numpy()[np.newaxis, :]

    n_patients = S.shape[0]
    F_raw = np.empty(n_patients)
    for p in range(n_patients):
        si = S[p, iu]
        sj = S[p, ju]
        term = Bij * si * sj
        F_raw[p] = np.sum(np.abs(Bij)[term > 0])

    F_norm = minmax_01(F_raw)

    output = pd.DataFrame(
        {"F_raw": F_raw, "F_norm": F_norm},
        index=reaction_expression.index,
    )
    output.to_csv(TABLE_DIR / "patient_frustration_scores.csv")
    return output


def calculate_binary_frustration(
    state_matrix: pd.DataFrame,
    B_matrix: pd.DataFrame,
) -> pd.DataFrame:
    """
    ALTERNATIVE / not used for Figure 3D in the current manuscript.

    Calculate normalized frustration for each patient/configuration directly
    from patient-specific optimized binary states (requires
    optimized_binary_states.csv). Kept for reference and for use cases where
    per-patient optimized configurations are available; see
    `calculate_frustration_index` above for the definition actually reported
    in Figure 3D, which does not require this input.

    This implementation uses:
        raw_F = 0.5 * sigma^T B sigma
        F_norm = (raw_F - min(raw_F)) / (max(raw_F) - min(raw_F))

    Binary 0/1 reaction states are converted to spin states -1/+1.
    """
    common = state_matrix.columns.intersection(B_matrix.index).intersection(B_matrix.columns)
    if len(common) == 0:
        raise ValueError("No overlapping reaction IDs between state matrix and B matrix.")

    B = B_matrix.loc[common, common].astype(float).to_numpy()
    B = (B + B.T) / 2.0
    np.fill_diagonal(B, 0.0)

    X = state_matrix[common].astype(float).to_numpy()
    spins = np.where(X > 0, 1.0, -1.0)

    raw_F = 0.5 * np.einsum("bi,ij,bj->b", spins, B, spins)

    denominator = np.max(raw_F) - np.min(raw_F)
    if denominator <= EPSILON:
        F_norm = np.zeros_like(raw_F)
    else:
        F_norm = (raw_F - np.min(raw_F)) / denominator

    output = pd.DataFrame(
        {"F_raw": raw_F, "F_norm": F_norm},
        index=state_matrix.index,
    )
    output.to_csv(TABLE_DIR / "patient_frustration_scores.csv")
    return output


def calculate_continuous_frustration(
    reaction_expression: pd.DataFrame,
    B_matrix: pd.DataFrame,
) -> pd.DataFrame:
    """
    Alternative patient-level frustration using continuous z-scored reaction
    states instead of optimized binary states.
    """
    common = (
        reaction_expression.columns
        .intersection(B_matrix.index)
        .intersection(B_matrix.columns)
    )
    z = zscore_columns(reaction_expression[common]).to_numpy()

    B = B_matrix.loc[common, common].astype(float).to_numpy()
    B = (B + B.T) / 2.0
    np.fill_diagonal(B, 0.0)

    raw_F = 0.5 * np.einsum("bi,ij,bj->b", z, B, z)
    F_norm = minmax_01(raw_F)

    return pd.DataFrame(
        {"F_raw_continuous": raw_F, "F_norm_continuous": F_norm},
        index=reaction_expression.index,
    )


def correlation_order_frustration(
    metrics: pd.DataFrame,
    filename: str = "order_vs_frustration.png",
) -> Tuple[float, float]:
    """Pearson correlation and scatterplot for m versus F_norm."""
    data = metrics[["m", "F_norm"]].dropna()
    r_value, p_value = pearsonr(data["m"], data["F_norm"])

    slope, intercept, _, _, _ = stats.linregress(data["m"], data["F_norm"])
    x_line = np.linspace(data["m"].min(), data["m"].max(), 200)

    fig, ax = plt.subplots(figsize=(6, 5))
    ax.scatter(data["m"], data["F_norm"], alpha=0.7, s=28)
    ax.plot(x_line, intercept + slope * x_line)
    ax.set_xlabel("Model-derived order parameter, m")
    ax.set_ylabel("Normalized frustration, F")
    ax.set_title(f"Pearson r = {r_value:.3f}, p = {p_value:.3g}")
    save_figure(fig, filename)

    return r_value, p_value


# =============================================================================
# 6. PATHWAY-LEVEL CONFIGURATION DIVERSITY
# Corresponding Methods:
#   - Pathway-level analysis
#   - HQA partition projection
# Relevant panels:
#   - Figure 1G-H
#   - Figure 4G
# =============================================================================

def shannon_binary_entropy(active_fraction: float) -> float:
    """
    Shannon entropy of a binary active/inactive distribution:
        H = -p log2(p) - (1-p) log2(1-p)
    """
    p = float(np.clip(active_fraction, EPSILON, 1.0 - EPSILON))
    return -(p * np.log2(p) + (1.0 - p) * np.log2(1.0 - p))


def pathway_configuration_diversity(
    binary_states: pd.DataFrame,
    pathway_map: pd.DataFrame,
) -> pd.DataFrame:
    """
    Calculate patient-level pathway Shannon configuration-diversity indices.

    Required pathway_map columns:
      reaction_id, pathway
    """
    required = {REACTION_ID_COL, PATHWAY_COL}
    missing = required.difference(pathway_map.columns)
    if missing:
        raise ValueError(f"Missing pathway-map columns: {sorted(missing)}")

    result = pd.DataFrame(index=binary_states.index)

    for pathway_name, group in pathway_map.groupby(PATHWAY_COL):
        reactions = [
            r for r in group[REACTION_ID_COL].astype(str)
            if r in binary_states.columns
        ]
        if not reactions:
            continue

        active_fraction = binary_states[reactions].astype(float).mean(axis=1)
        result[pathway_name] = active_fraction.apply(shannon_binary_entropy)

    result.to_csv(TABLE_DIR / "patient_pathway_configuration_diversity.csv")
    return result


def compare_pathway_diversity_between_groups(
    pathway_entropy: pd.DataFrame,
    metadata: pd.DataFrame,
    group_a: str = "Stem-like",
    group_b: str = "Inflammatory",
    group_col: str = MOLECULAR_SUBTYPE_COL,
) -> pd.DataFrame:
    """
    Compare pathway-level configuration diversity between two groups using the
    Mann-Whitney U test.
    """
    aligned_meta = metadata.loc[pathway_entropy.index]
    rows = []

    for pathway in pathway_entropy.columns:
        a = pathway_entropy.loc[aligned_meta[group_col] == group_a, pathway].dropna()
        b = pathway_entropy.loc[aligned_meta[group_col] == group_b, pathway].dropna()

        if len(a) < 2 or len(b) < 2:
            continue

        statistic, p_value = mannwhitneyu(a, b, alternative="two-sided")
        rows.append(
            {
                "pathway": pathway,
                f"mean_{group_a}": a.mean(),
                f"mean_{group_b}": b.mean(),
                "difference": a.mean() - b.mean(),
                "U_statistic": statistic,
                "p_value": p_value,
            }
        )

    output = pd.DataFrame(rows).sort_values("p_value")
    if not output.empty:
        output["FDR_BH"] = stats.false_discovery_control(output["p_value"].to_numpy())
    output.to_csv(TABLE_DIR / "pathway_diversity_group_comparison.csv", index=False)
    return output


def plot_pathway_difference(
    comparison: pd.DataFrame,
    top_n: int = 20,
    filename: str = "pathway_diversity_difference.png",
) -> None:
    """Horizontal bar plot of the largest pathway-level differences."""
    selected = (
        comparison.assign(abs_difference=lambda d: d["difference"].abs())
        .nlargest(top_n, "abs_difference")
        .sort_values("difference")
    )

    fig, ax = plt.subplots(figsize=(8, max(5, 0.3 * len(selected))))
    ax.barh(selected["pathway"], selected["difference"])
    ax.axvline(0, linewidth=1)
    ax.set_xlabel("Mean diversity difference: Stem-like - Inflammatory")
    ax.set_ylabel("Pathway")
    ax.set_title("Pathway-level configuration-diversity difference")
    save_figure(fig, filename)


# =============================================================================
# 7. EMPIRICAL PROBABILITY-DERIVED STATISTICAL POTENTIAL LANDSCAPES
# Corresponding Methods:
#   - Construction of subtype-specific probability-derived landscapes
#   - Two-dimensional Gaussian KDE with Scott's rule
# Relevant panels:
#   - Figure 1B-C, E-F
#   - Figure 2A-C
#   - Figure 4K-L
#
# NOTE (post-review correction): the original Figure 3H-J (one-dimensional
# statistical potential F(m) and basin/barrier estimates between stem-like
# and inflammatory subtypes) have been removed from the manuscript. An
# independent check using `plot_1d_potential_by_group` below with the plain,
# Methods-defined order parameter `m` (no subtype-discriminant projection)
# showed no significant difference in m between stem-like and inflammatory
# patients (Mann-Whitney p = 0.69) and no discernible basin or energy-barrier
# structure; see AUDIT_NOTES.md for details. The functions in this section
# are retained because they are also used for Figures 1, 2, and 4.
# =============================================================================

def kde2d_landscape(
    x: Sequence[float],
    y: Sequence[float],
    gridsize: int = 150,
    bandwidth: str | float = "scott",
    padding_fraction: float = 0.08,
) -> Dict[str, np.ndarray]:
    """
    Estimate a 2D Gaussian KDE and convert it to a statistical potential:
        F(x, y) = -ln[P(x, y) + epsilon]
    The minimum potential is shifted to zero.
    """
    x = np.asarray(x, dtype=float)
    y = np.asarray(y, dtype=float)
    valid = np.isfinite(x) & np.isfinite(y)
    x = x[valid]
    y = y[valid]

    if len(x) < 3:
        raise ValueError("At least three valid observations are required for 2D KDE.")

    x_range = x.max() - x.min()
    y_range = y.max() - y.min()
    x_pad = max(x_range * padding_fraction, 1e-6)
    y_pad = max(y_range * padding_fraction, 1e-6)

    x_grid = np.linspace(x.min() - x_pad, x.max() + x_pad, gridsize)
    y_grid = np.linspace(y.min() - y_pad, y.max() + y_pad, gridsize)
    X, Y = np.meshgrid(x_grid, y_grid)

    positions = np.vstack([X.ravel(), Y.ravel()])
    samples = np.vstack([x, y])
    kde = gaussian_kde(samples, bw_method=bandwidth)
    density = np.reshape(kde(positions), X.shape)
    density = density / np.sum(density)

    potential = safe_log_probability(density)
    return {
        "X": X,
        "Y": Y,
        "density": density,
        "potential": potential,
    }


def plot_landscape_3d(
    landscape: Dict[str, np.ndarray],
    xlabel: str,
    ylabel: str,
    title: str,
    filename: str,
) -> None:
    """Three-dimensional statistical-potential surface."""
    fig = plt.figure(figsize=(8, 6))
    ax = fig.add_subplot(111, projection="3d")
    ax.plot_surface(
        landscape["X"],
        landscape["Y"],
        landscape["potential"],
        linewidth=0,
        antialiased=True,
        alpha=0.95,
    )
    ax.set_xlabel(xlabel)
    ax.set_ylabel(ylabel)
    ax.set_zlabel("Statistical potential, -ln(P)")
    ax.set_title(title)
    save_figure(fig, filename)


def plot_landscape_contour(
    landscape: Dict[str, np.ndarray],
    xlabel: str,
    ylabel: str,
    title: str,
    filename: str,
) -> None:
    """Two-dimensional filled-contour plot."""
    fig, ax = plt.subplots(figsize=(7, 6))
    contour = ax.contourf(
        landscape["X"],
        landscape["Y"],
        landscape["potential"],
        levels=30,
    )
    fig.colorbar(contour, ax=ax, label="Statistical potential, -ln(P)")
    ax.set_xlabel(xlabel)
    ax.set_ylabel(ylabel)
    ax.set_title(title)
    save_figure(fig, filename)


def subtype_landscapes(
    coordinates: pd.DataFrame,
    metadata: pd.DataFrame,
    x_col: str,
    y_col: str,
    group_col: str = MOLECULAR_SUBTYPE_COL,
) -> Dict[str, Dict[str, np.ndarray]]:
    """Generate and save a landscape for every subtype."""
    data = coordinates.join(metadata[[group_col]], how="inner")
    landscapes = {}

    for subtype, group in data.groupby(group_col):
        landscape = kde2d_landscape(group[x_col], group[y_col])
        landscapes[str(subtype)] = landscape

        safe_name = str(subtype).replace(" ", "_").replace("/", "_")
        plot_landscape_3d(
            landscape,
            xlabel=x_col,
            ylabel=y_col,
            title=f"{subtype}: probability-derived statistical potential",
            filename=f"landscape_3D_{safe_name}.png",
        )
        plot_landscape_contour(
            landscape,
            xlabel=x_col,
            ylabel=y_col,
            title=f"{subtype}: probability-derived statistical potential",
            filename=f"landscape_contour_{safe_name}.png",
        )

    return landscapes


def difference_landscape(
    landscape_a: Dict[str, np.ndarray],
    landscape_b: Dict[str, np.ndarray],
    title: str,
    filename: str,
) -> None:
    """
    Plot a direct difference map. Landscapes must use the same coordinate grid.
    For strict comparison, construct both using common global x/y limits.
    """
    if landscape_a["potential"].shape != landscape_b["potential"].shape:
        raise ValueError("Landscapes must have the same grid shape.")

    difference = landscape_a["potential"] - landscape_b["potential"]

    fig, ax = plt.subplots(figsize=(7, 6))
    contour = ax.contourf(
        landscape_a["X"],
        landscape_a["Y"],
        difference,
        levels=30,
    )
    fig.colorbar(contour, ax=ax, label="Difference in statistical potential")
    ax.set_xlabel("m1")
    ax.set_ylabel("m2")
    ax.set_title(title)
    save_figure(fig, filename)


def one_dimensional_potential(
    values: Sequence[float],
    gridsize: int = 300,
    bandwidth: str | float = "scott",
) -> pd.DataFrame:
    """Calculate a 1D probability-derived statistical potential F(m)."""
    values = np.asarray(values, dtype=float)
    values = values[np.isfinite(values)]

    grid = np.linspace(values.min(), values.max(), gridsize)
    kde = gaussian_kde(values, bw_method=bandwidth)
    density = kde(grid)
    density = density / density.sum()
    potential = safe_log_probability(density)

    return pd.DataFrame({"coordinate": grid, "density": density, "potential": potential})


def plot_1d_potential_by_group(
    metrics: pd.DataFrame,
    metadata: pd.DataFrame,
    coordinate_col: str = "m",
    group_col: str = MOLECULAR_SUBTYPE_COL,
    filename: str = "one_dimensional_potential_by_subtype.png",
) -> None:
    """Plot F(m) curves for all subtypes."""
    data = metrics.join(metadata[[group_col]], how="inner")

    fig, ax = plt.subplots(figsize=(8, 5))
    for subtype, group in data.groupby(group_col):
        curve = one_dimensional_potential(group[coordinate_col])
        ax.plot(curve["coordinate"], curve["potential"], label=str(subtype))

    ax.set_xlabel(coordinate_col)
    ax.set_ylabel("Statistical potential, -ln(P)")
    ax.set_title("Subtype-specific statistical-potential profiles")
    ax.legend(frameon=False, fontsize=8)
    save_figure(fig, filename)


# =============================================================================
# 8. PARAMETER-SWEEP / PHASE-MAP ANALYSIS
# Corresponding Methods:
#   - SC-MSG objective-function sensitivity analysis
#   - Identification of metabolic phases
# Relevant panels:
#   - Figure 1B-C, F
#   - Figure 2B-C
# =============================================================================

def plot_parameter_heatmap(
    sweep_data: pd.DataFrame,
    value_col: str,
    title: str,
    filename: str,
    w1_col: str = "w1",
    w2_col: str = "w2",
) -> None:
    """Plot a w1-w2 parameter-sweep heatmap."""
    pivot = sweep_data.pivot_table(
        index=w2_col,
        columns=w1_col,
        values=value_col,
        aggfunc="mean",
    ).sort_index(ascending=True)

    fig, ax = plt.subplots(figsize=(7, 6))
    image = ax.imshow(
        pivot.to_numpy(),
        origin="lower",
        aspect="auto",
        extent=[
            pivot.columns.min(),
            pivot.columns.max(),
            pivot.index.min(),
            pivot.index.max(),
        ],
    )
    fig.colorbar(image, ax=ax, label=value_col)
    ax.set_xlabel("w1: cofactor-interaction weight")
    ax.set_ylabel("w2: transcriptomic-field weight")
    ax.set_title(title)
    save_figure(fig, filename)


def calculate_transition_strength(
    stem_sweep: pd.DataFrame,
    inflammatory_sweep: pd.DataFrame,
    order_col: str = "m",
) -> pd.DataFrame:
    """Calculate Delta m = m_inflammatory - m_stem across matched w1-w2 points."""
    keys = ["w1", "w2"]
    merged = stem_sweep[keys + [order_col]].merge(
        inflammatory_sweep[keys + [order_col]],
        on=keys,
        suffixes=("_stem", "_inflammatory"),
    )
    merged["delta_m"] = (
        merged[f"{order_col}_inflammatory"] - merged[f"{order_col}_stem"]
    )
    merged.to_csv(TABLE_DIR / "parameter_sweep_transition_strength.csv", index=False)
    return merged


# =============================================================================
# 9. GROUP COMPARISON AND COMPOSITION PLOTS
# Relevant panels:
#   - Figure 3C-E, K-L
#   - Figure 4A, E-G, J
# =============================================================================

def kruskal_wallis_by_group(
    data: pd.DataFrame,
    value_col: str,
    group_col: str,
) -> Tuple[float, float]:
    """Kruskal-Wallis test across multiple groups."""
    arrays = [
        group[value_col].dropna().to_numpy()
        for _, group in data.groupby(group_col)
        if group[value_col].notna().sum() > 0
    ]
    statistic, p_value = kruskal(*arrays)
    return statistic, p_value


def plot_metric_by_group(
    data: pd.DataFrame,
    value_col: str,
    group_col: str,
    filename: str,
) -> None:
    """Boxplot plus jittered observations."""
    groups = list(data[group_col].dropna().astype(str).unique())
    arrays = [
        data.loc[data[group_col].astype(str) == group, value_col].dropna().to_numpy()
        for group in groups
    ]

    fig, ax = plt.subplots(figsize=(max(7, len(groups) * 1.2), 5))
    ax.boxplot(arrays, labels=groups, showfliers=False)

    rng = np.random.default_rng(RANDOM_SEED)
    for i, arr in enumerate(arrays, start=1):
        jitter = rng.normal(i, 0.05, size=len(arr))
        ax.scatter(jitter, arr, s=18, alpha=0.55)

    statistic, p_value = kruskal(*arrays)
    ax.set_ylabel(value_col)
    ax.set_xlabel(group_col)
    ax.set_title(f"Kruskal-Wallis p = {p_value:.3g}")
    ax.tick_params(axis="x", rotation=35)
    save_figure(fig, filename)


def plot_composition_stacked(
    data: pd.DataFrame,
    x_col: str,
    composition_col: str,
    filename: str,
) -> pd.DataFrame:
    """Stacked proportional bar chart for subtype composition."""
    counts = pd.crosstab(data[x_col], data[composition_col])
    proportions = counts.div(counts.sum(axis=1), axis=0)

    fig, ax = plt.subplots(figsize=(8, 5))
    proportions.plot(kind="bar", stacked=True, ax=ax)
    ax.set_ylabel("Proportion")
    ax.set_xlabel(x_col)
    ax.set_title(f"{composition_col} composition within {x_col}")
    ax.legend(title=composition_col, bbox_to_anchor=(1.02, 1), loc="upper left")
    save_figure(fig, filename)

    counts.to_csv(TABLE_DIR / f"composition_counts_{x_col}_by_{composition_col}.csv")
    proportions.to_csv(TABLE_DIR / f"composition_proportions_{x_col}_by_{composition_col}.csv")
    return proportions


# =============================================================================
# 10. SURVIVAL ANALYSIS
# Corresponding Methods:
#   - Kaplan-Meier analysis
#   - Log-rank tests
#   - Multivariable Cox proportional-hazards models
#   - Transcriptomic PCA-adjusted benchmark
# Relevant panels:
#   - Figure 4B-D, H-I
#   - Supplementary Table 1
# =============================================================================

def clean_survival_data(
    data: pd.DataFrame,
    time_col: str,
    event_col: str,
) -> pd.DataFrame:
    """Validate and clean survival variables."""
    output = data.copy()
    output[time_col] = pd.to_numeric(output[time_col], errors="coerce")
    output[event_col] = pd.to_numeric(output[event_col], errors="coerce")
    output = output.dropna(subset=[time_col, event_col])
    output = output.loc[output[time_col] >= 0]
    output[event_col] = output[event_col].astype(int)
    return output


def kaplan_meier_by_group(
    data: pd.DataFrame,
    time_col: str,
    event_col: str,
    group_col: str,
    filename: str,
    title: str,
) -> Dict[str, float]:
    """
    Kaplan-Meier curves and overall multivariate log-rank test.
    """
    data = clean_survival_data(data, time_col, event_col)
    groups = data[group_col].dropna().astype(str).unique()

    fig, ax = plt.subplots(figsize=(7, 6))
    kmf = KaplanMeierFitter()

    for group_name in sorted(groups):
        subset = data.loc[data[group_col].astype(str) == group_name]
        if subset.empty:
            continue
        kmf.fit(
            subset[time_col],
            event_observed=subset[event_col],
            label=str(group_name),
        )
        kmf.plot_survival_function(ax=ax, ci_show=False)

    overall = multivariate_logrank_test(
        data[time_col],
        data[group_col],
        data[event_col],
    )

    ax.set_xlabel("Time")
    ax.set_ylabel("Survival probability")
    ax.set_title(f"{title}\nOverall log-rank p = {overall.p_value:.3g}")
    ax.legend(frameon=False)
    save_figure(fig, filename)

    return {
        "test_statistic": float(overall.test_statistic),
        "p_value": float(overall.p_value),
    }


def pairwise_logrank(
    data: pd.DataFrame,
    time_col: str,
    event_col: str,
    group_col: str,
    group_a: str,
    group_b: str,
) -> Dict[str, float]:
    """Pairwise log-rank test."""
    data = clean_survival_data(data, time_col, event_col)
    a = data.loc[data[group_col].astype(str) == str(group_a)]
    b = data.loc[data[group_col].astype(str) == str(group_b)]

    result = logrank_test(
        a[time_col],
        b[time_col],
        event_observed_A=a[event_col],
        event_observed_B=b[event_col],
    )

    return {
        "group_a": group_a,
        "group_b": group_b,
        "n_a": len(a),
        "n_b": len(b),
        "test_statistic": float(result.test_statistic),
        "p_value": float(result.p_value),
    }


def median_dichotomization(
    data: pd.DataFrame,
    value_col: str,
    output_col: str,
    low_label: str = "Low",
    high_label: str = "High",
) -> pd.DataFrame:
    """Dichotomize a continuous variable at its cohort median."""
    output = data.copy()
    median_value = output[value_col].median()
    output[output_col] = np.where(
        output[value_col] >= median_value,
        high_label,
        low_label,
    )
    return output


def prepare_cox_covariates(
    data: pd.DataFrame,
    columns: Sequence[str],
) -> pd.DataFrame:
    """
    Convert categorical clinical covariates to indicator variables and retain
    continuous covariates.
    """
    subset = data[list(columns)].copy()

    categorical = [
        col for col in subset.columns
        if subset[col].dtype == "object" or str(subset[col].dtype).startswith("category")
    ]
    subset = pd.get_dummies(subset, columns=categorical, drop_first=True, dtype=float)
    return subset


def fit_cox_model(
    data: pd.DataFrame,
    time_col: str,
    event_col: str,
    covariates: Sequence[str],
    output_name: str,
) -> CoxPHFitter:
    """Fit and save a multivariable Cox proportional-hazards model."""
    survival = data[[time_col, event_col]].copy()
    X = prepare_cox_covariates(data, covariates)
    model_data = pd.concat([survival, X], axis=1).dropna()

    cph = CoxPHFitter()
    cph.fit(model_data, duration_col=time_col, event_col=event_col)

    cph.summary.to_csv(TABLE_DIR / f"{output_name}_Cox_summary.csv")
    return cph


def likelihood_ratio_increment_test(
    reduced_model: CoxPHFitter,
    full_model: CoxPHFitter,
    added_parameters: int = 1,
) -> Dict[str, float]:
    """
    Likelihood-ratio test comparing nested Cox models.
    """
    lr_stat = 2.0 * (
        full_model.log_likelihood_ - reduced_model.log_likelihood_
    )
    p_value = stats.chi2.sf(lr_stat, df=added_parameters)
    return {"LR_statistic": float(lr_stat), "p_value": float(p_value)}


def pca_adjusted_survival_analysis(
    analysis_data: pd.DataFrame,
    pca_scores: pd.DataFrame,
    outcome: str = "OS",
    number_of_pcs: int = 5,
) -> Dict[str, float]:
    """
    Compare:
      Reduced model = age + sex + pathological stage + PC1...PCk
      Full model    = reduced model + m

    This corresponds to the manuscript's transcriptomic PCA-adjusted benchmark.
    """
    if outcome.upper() == "OS":
        time_col, event_col = OS_TIME_COL, OS_EVENT_COL
    elif outcome.upper() == "DFS":
        time_col, event_col = DFS_TIME_COL, DFS_EVENT_COL
    else:
        raise ValueError("outcome must be 'OS' or 'DFS'")

    pcs = [f"PC{i}" for i in range(1, number_of_pcs + 1)]
    merged = analysis_data.join(pca_scores[pcs], how="inner")

    base_covariates = [AGE_COL, SEX_COL, STAGE_COL] + pcs
    full_covariates = base_covariates + ["m"]

    reduced = fit_cox_model(
        merged,
        time_col,
        event_col,
        base_covariates,
        output_name=f"{outcome}_clinical_PC1_PC{number_of_pcs}",
    )
    full = fit_cox_model(
        merged,
        time_col,
        event_col,
        full_covariates,
        output_name=f"{outcome}_clinical_PC1_PC{number_of_pcs}_plus_m",
    )

    comparison = likelihood_ratio_increment_test(reduced, full, added_parameters=1)
    comparison.update(
        {
            "outcome": outcome,
            "number_of_PCs": number_of_pcs,
            "reduced_log_likelihood": float(reduced.log_likelihood_),
            "full_log_likelihood": float(full.log_likelihood_),
        }
    )
    return comparison


# =============================================================================
# 11. OPTIONAL QUBO / SC-MSG OBJECTIVE CALCULATION
# Corresponding Methods:
#   - Sung-Cheong Metabolic Spin-Glass Model
#   - Optimization of effective metabolic ground states
# Relevant panel:
#   - Figure 1A
#
# This section calculates the model objective for an already supplied state.
# Exact D-Wave hybrid execution requires valid D-Wave credentials and the
# final QUBO scaling used by the authors.
# =============================================================================

def sc_msg_objective(
    x: np.ndarray,
    delta_g: np.ndarray,
    B: np.ndarray,
    S: np.ndarray,
    w1: float,
    w2: float,
) -> float:
    """
    Effective SC-MSG objective:
        H(x) = sum_i DeltaG_i x_i
             + w1 * sum_{i<j} B_ij x_i x_j
             + w2 * sum_i S_i x_i

    All inputs must follow the same reaction order.
    """
    x = np.asarray(x, dtype=float)
    delta_g = np.asarray(delta_g, dtype=float)
    B = np.asarray(B, dtype=float)
    S = np.asarray(S, dtype=float)

    linear_energy = np.dot(delta_g, x)
    interaction_energy = 0.5 * np.dot(x, B @ x)
    transcriptomic_energy = np.dot(S, x)

    return float(linear_energy + w1 * interaction_energy + w2 * transcriptomic_energy)


def build_qubo_dictionary(
    delta_g: pd.Series,
    B_matrix: pd.DataFrame,
    transcriptomic_field: pd.Series,
    w1: float,
    w2: float,
) -> Dict[Tuple[str, str], float]:
    """
    Convert the SC-MSG objective into a QUBO dictionary compatible with dimod.

    Diagonal:
        Q_ii = DeltaG_i + w2*S_i

    Off-diagonal:
        Q_ij = w1*B_ij, i < j
    """
    reactions = (
        delta_g.index
        .intersection(B_matrix.index)
        .intersection(B_matrix.columns)
        .intersection(transcriptomic_field.index)
    )

    Q = {}
    for reaction in reactions:
        Q[(reaction, reaction)] = float(
            delta_g.loc[reaction] + w2 * transcriptomic_field.loc[reaction]
        )

    for i, reaction_i in enumerate(reactions):
        for reaction_j in reactions[i + 1:]:
            coefficient = float(w1 * B_matrix.loc[reaction_i, reaction_j])
            if abs(coefficient) > EPSILON:
                Q[(reaction_i, reaction_j)] = coefficient

    return Q


def solve_qubo_with_dwave_hybrid(Q: Dict[Tuple[str, str], float]):
    """
    Optional D-Wave hybrid solver.

    Requires:
      pip install dwave-system dimod
      valid D-Wave API configuration

    Returns the lowest-energy sample and energy.
    """
    try:
        from dwave.system import LeapHybridSampler
    except ImportError as exc:
        raise ImportError(
            "Install dwave-system before using the D-Wave hybrid solver."
        ) from exc

    sampler = LeapHybridSampler()
    sampleset = sampler.sample_qubo(Q)
    best = sampleset.first
    return best.sample, float(best.energy)


# =============================================================================
# 12. COMPLETE ANALYSIS WORKFLOW
# =============================================================================

def main() -> None:
    # -------------------------------------------------------------------------
    # Load core data
    # -------------------------------------------------------------------------
    reaction_expression = load_reaction_expression()

    annotations = read_table(REACTION_ANNOTATION_FILE)
    if REACTION_ID_COL not in annotations.columns or DG_COL not in annotations.columns:
        raise ValueError(
            f"Reaction annotation must contain '{REACTION_ID_COL}' and '{DG_COL}'."
        )
    annotations[REACTION_ID_COL] = annotations[REACTION_ID_COL].astype(str)
    delta_g = annotations.set_index(REACTION_ID_COL)[DG_COL].astype(float)

    B_matrix = read_table(B_MATRIX_FILE, index_col=0)
    B_matrix.index = B_matrix.index.astype(str)
    B_matrix.columns = B_matrix.columns.astype(str)

    clinical = read_table(CLINICAL_FILE)
    if PATIENT_COL in clinical.columns:
        clinical = clinical.set_index(PATIENT_COL)

    optimized_states = read_table(OPTIMIZED_STATE_FILE)
    if PATIENT_COL in optimized_states.columns:
        optimized_states = optimized_states.set_index(PATIENT_COL)

    pathway_map = read_table(PATHWAY_MAP_FILE)

    # Align patients across primary input files
    common_patients = (
        reaction_expression.index
        .intersection(clinical.index)
        .intersection(optimized_states.index)
    )
    reaction_expression = reaction_expression.loc[common_patients]
    clinical = clinical.loc[common_patients]
    optimized_states = optimized_states.loc[common_patients]

    # -------------------------------------------------------------------------
    # PCA on patient-by-reaction expression matrix
    # -------------------------------------------------------------------------
    pca_scores, pca_loadings, pca_explained = run_pca(
        reaction_expression,
        n_components=10,
        standardize=True,
    )
    plot_pca_scatter(
        pca_scores,
        clinical,
        group_col=MOLECULAR_SUBTYPE_COL,
        filename="PCA_reaction_expression_by_molecular_subtype.png",
    )

    # -------------------------------------------------------------------------
    # Model-derived order and frustration
    # -------------------------------------------------------------------------
    order_metrics = calculate_order_parameter(reaction_expression, delta_g)
    frustration_metrics = calculate_frustration_index(reaction_expression, delta_g, B_matrix)

    metrics = order_metrics.join(frustration_metrics, how="inner")
    correlation_order_frustration(metrics)

    # -------------------------------------------------------------------------
    # Pathway-level configuration diversity
    # -------------------------------------------------------------------------
    pathway_entropy = pathway_configuration_diversity(
        optimized_states,
        pathway_map,
    )

    pathway_comparison = compare_pathway_diversity_between_groups(
        pathway_entropy,
        clinical,
        group_a="Stem-like",
        group_b="Inflammatory",
        group_col=MOLECULAR_SUBTYPE_COL,
    )
    if not pathway_comparison.empty:
        plot_pathway_difference(pathway_comparison)

    # -------------------------------------------------------------------------
    # m1 and m2 collective coordinates from pathway entropy PCA
    # -------------------------------------------------------------------------
    collective_coordinates = construct_collective_coordinates_from_pathway_entropy(
        pathway_entropy
    )

    subtype_landscapes(
        collective_coordinates,
        clinical,
        x_col="m1",
        y_col="m2",
        group_col=MOLECULAR_SUBTYPE_COL,
    )

    # Landscape in m-F space
    mf_coordinates = metrics[["m", "F_norm"]]
    subtype_landscapes(
        mf_coordinates,
        clinical,
        x_col="m",
        y_col="F_norm",
        group_col=MOLECULAR_SUBTYPE_COL,
    )

    # Verification check (not a main-text figure as of the current revision):
    # this reproduces the test used to confirm that stem-like and
    # inflammatory subtypes do NOT occupy distinguishable free-energy basins
    # once the plain, Methods-defined order parameter m is used directly
    # (i.e., without a subtype-discriminant projection axis). This is the
    # basis for removing the original Figure 3H-J; see AUDIT_NOTES.md.
    plot_1d_potential_by_group(
        metrics,
        clinical,
        coordinate_col="m",
        group_col=MOLECULAR_SUBTYPE_COL,
        filename="verification_1D_potential_by_subtype_no_basin_found.png",
    )

    # -------------------------------------------------------------------------
    # GMM SC-MSG-derived subtype assignment
    # -------------------------------------------------------------------------
    sc_msg_assignments = assign_sc_msg_subtypes(
        metrics[["abs_m", "F_norm"]],
        n_components=5,
    )

    full_data = (
        clinical
        .join(metrics, how="inner")
        .join(
            sc_msg_assignments[
                [SC_MSG_SUBTYPE_COL, "cluster_probability"]
            ],
            how="inner",
        )
    )

    plot_metric_by_group(
        full_data,
        value_col="m",
        group_col=SC_MSG_SUBTYPE_COL,
        filename="order_parameter_by_sc_msg_subtype.png",
    )
    plot_metric_by_group(
        full_data,
        value_col="F_norm",
        group_col=SC_MSG_SUBTYPE_COL,
        filename="frustration_by_sc_msg_subtype.png",
    )

    plot_composition_stacked(
        full_data,
        x_col=SC_MSG_SUBTYPE_COL,
        composition_col=MOLECULAR_SUBTYPE_COL,
        filename="molecular_subtype_composition_by_sc_msg_subtype.png",
    )

    # -------------------------------------------------------------------------
    # Survival analyses
    # -------------------------------------------------------------------------
    kaplan_meier_by_group(
        full_data,
        time_col=OS_TIME_COL,
        event_col=OS_EVENT_COL,
        group_col=SC_MSG_SUBTYPE_COL,
        filename="KM_OS_all_sc_msg_subtypes.png",
        title="Overall survival by SC-MSG-derived subtype",
    )

    kaplan_meier_by_group(
        full_data,
        time_col=DFS_TIME_COL,
        event_col=DFS_EVENT_COL,
        group_col=SC_MSG_SUBTYPE_COL,
        filename="KM_DFS_all_sc_msg_subtypes.png",
        title="Disease-free survival by SC-MSG-derived subtype",
    )

    # Stem-like subset
    stem_like = full_data.loc[
        full_data[MOLECULAR_SUBTYPE_COL].astype(str).str.lower() == "stem-like"
    ].copy()

    if not stem_like.empty:
        kaplan_meier_by_group(
            stem_like,
            time_col=OS_TIME_COL,
            event_col=OS_EVENT_COL,
            group_col=SC_MSG_SUBTYPE_COL,
            filename="KM_OS_stem_like_sc_msg_subtypes.png",
            title="Overall survival in stem-like tumors",
        )

        kaplan_meier_by_group(
            stem_like,
            time_col=DFS_TIME_COL,
            event_col=DFS_EVENT_COL,
            group_col=SC_MSG_SUBTYPE_COL,
            filename="KM_DFS_stem_like_sc_msg_subtypes.png",
            title="Disease-free survival in stem-like tumors",
        )

        pairwise_results = []
        for outcome, time_col, event_col in [
            ("OS", OS_TIME_COL, OS_EVENT_COL),
            ("DFS", DFS_TIME_COL, DFS_EVENT_COL),
        ]:
            for group_a, group_b in [("T2", "T1"), ("T5", "T1")]:
                try:
                    result = pairwise_logrank(
                        stem_like,
                        time_col,
                        event_col,
                        SC_MSG_SUBTYPE_COL,
                        group_a,
                        group_b,
                    )
                    result["outcome"] = outcome
                    pairwise_results.append(result)
                except Exception:
                    pass

        pd.DataFrame(pairwise_results).to_csv(
            TABLE_DIR / "stem_like_pairwise_logrank_tests.csv",
            index=False,
        )

    # Median dichotomization of m for DFS
    high_low_m = median_dichotomization(
        full_data,
        value_col="m",
        output_col="m_group",
        low_label="Low m",
        high_label="High m",
    )
    kaplan_meier_by_group(
        high_low_m,
        time_col=DFS_TIME_COL,
        event_col=DFS_EVENT_COL,
        group_col="m_group",
        filename="KM_DFS_high_vs_low_m.png",
        title="Disease-free survival by model-derived order parameter",
    )

    # PCA-adjusted Cox models
    cox_comparisons = []
    for outcome in ("OS", "DFS"):
        for k in (5, 10):
            try:
                result = pca_adjusted_survival_analysis(
                    full_data,
                    pca_scores,
                    outcome=outcome,
                    number_of_pcs=k,
                )
                cox_comparisons.append(result)
            except Exception as exc:
                cox_comparisons.append(
                    {
                        "outcome": outcome,
                        "number_of_PCs": k,
                        "error": str(exc),
                    }
                )

    pd.DataFrame(cox_comparisons).to_csv(
        TABLE_DIR / "PCA_adjusted_Cox_model_comparisons.csv",
        index=False,
    )

    # -------------------------------------------------------------------------
    # Parameter-sweep plots, when available
    # -------------------------------------------------------------------------
    if PARAMETER_SWEEP_FILE.exists():
        sweep = read_table(PARAMETER_SWEEP_FILE)

        if "active_reaction_count" in sweep.columns:
            plot_parameter_heatmap(
                sweep,
                value_col="active_reaction_count",
                title="Active reaction count across SC-MSG parameter space",
                filename="parameter_sweep_active_reaction_count.png",
            )

        if "m" in sweep.columns:
            plot_parameter_heatmap(
                sweep,
                value_col="m",
                title="Order parameter across SC-MSG parameter space",
                filename="parameter_sweep_order_parameter.png",
            )

        if "F_norm" in sweep.columns:
            plot_parameter_heatmap(
                sweep,
                value_col="F_norm",
                title="Frustration across SC-MSG parameter space",
                filename="parameter_sweep_frustration.png",
            )

    # Save merged patient-level analysis table
    full_data.to_csv(TABLE_DIR / "complete_patient_level_analysis_table.csv")

    print("Analysis completed.")
    print(f"Tables:  {TABLE_DIR.resolve()}")
    print(f"Figures: {FIGURE_DIR.resolve()}")


# =============================================================================
# 13. FIGURE-PANEL CROSS-REFERENCE MAP
# =============================================================================
#
# Figure 1A
#   - Reaction-level expression mapping
#   - SC-MSG Hamiltonian / QUBO construction
#   - Cofactor interaction B-matrix
#
# Figure 1B-C
#   - w1-w2 parameter sweep
#   - Active reaction-count density / difference maps
#
# Figure 1D
#   - Distribution of active reactions per optimized solution
#
# Figure 1E
#   - Jaccard distances between active reaction sets
#
# Figure 1F
#   - Parameter-space phase-map projection
#
# Figure 1G
#   - Pathway-level Shannon configuration-diversity analysis
#
# Figure 1H
#   - Functional/pathway annotation of top reactions
#
# Figure 2A-B
#   - PCA-derived collective coordinates
#   - Two-dimensional Gaussian KDE
#   - F = -ln(P + epsilon)
#   - Statistical-potential difference map
#
# Figure 2C
#   - Subtype-specific probability-derived statistical-potential landscapes
#
# Figure 3A-C
#   - Patient-level model-derived order parameter
#   - Distribution and subtype comparison
#
# Figure 3D
#   - Pearson correlation of m and frustration
#
# Figure 3E
#   - Operational metabolic-regime classification
#
# Figure 3F-G
#   - Reaction-level contributions grouped by Delta G bins
#   - Reaction contribution heatmaps
#
# [Original Figure 3H-J removed post-review: one-dimensional statistical
#  potential F(m) and basin/barrier estimates between stem-like and
#  inflammatory subtypes, computed from a subtype-discriminant projection
#  axis. This construction was found to be circular (the axis was built
#  from the same stem-like/inflammatory contrast the panels were used to
#  demonstrate) and did not reproduce when tested with the plain,
#  Methods-defined order parameter m (see `plot_1d_potential_by_group` call
#  in main(), and AUDIT_NOTES.md). Figure 3K and 3L below are renumbered
#  Figure 3H and 3I in the current manuscript.]
#
# Figure 3H-I (previously 3K-L)
#   - GMM clustering in (|m|, F_norm)
#   - Molecular/SC-MSG-derived subtype composition
#
# Figure 4A
#   - m and F distributions across T1-T5
#
# Figure 4B-D, H-I
#   - Kaplan-Meier survival curves
#   - Overall and pairwise log-rank tests
#
# Figure 4E-G
#   - Mann-Whitney or Kruskal-Wallis group comparisons
#   - Order, frustration, and pathway entropy
#
# Figure 4J
#   - Stacked subtype-composition plot
#
# Figure 4K-L
#   - Patient projection and density/landscape plots in the (m, F) plane
#
# Supplementary Table 1
#   - Multivariable Cox regression adjusted for:
#       age, sex, pathological stage, and PC1-PC5 or PC1-PC10
#   - Nested-model likelihood-ratio tests for incremental contribution of m
#
# =============================================================================

if __name__ == "__main__":
    main()
```
