## supplementary code 2 for "A Spin-Glass Metabolic Hamiltonian optimized by Quantum Annealing Reveals Configurational Regimes of Cancer Metabolism": Supplementary_Code_2_order_parameter_and_survival_benchmarks.py

```
"""
Supplementary Code 2 for:
"A Spin-Glass Metabolic Hamiltonian optimized by Quantum Annealing
 Reveals Configurational Regimes of Cancer Metabolism"
Sung, Baek, Park, Bang, and Cheong.

Corrected computation of the patient-level model-derived order parameter (m)
and its associated transcriptome-adjusted survival benchmark analyses
(Supplementary Tables 1-6).

--------------------------------------------------------------------------
SCOPE OF THIS SCRIPT (see the accompanying Legend for the exact wording)
--------------------------------------------------------------------------
This script does ONLY the following:
    1. Recomputes the patient-level model-derived order parameter m strictly
       according to the formula given in Methods, using all 235 reactions
       (sigma_r = sign(-dG_r); z_ir = per-reaction z-score across the full
       497-patient cohort), superseding the reaction-subset/sign-convention
       used to generate the order-parameter values originally reported in
       Supplementary Table 6.
    2. Reports the fraction of variance in m explained by the leading
       transcriptomic principal components (PC1-PC5, PC1-PC10) of the
       235-reaction expression matrix.
    3. Fits multivariable Cox proportional-hazards models for overall
       survival (OS) and disease-free survival (DFS), comparing a reduced
       model (age, sex, pathological stage, PC1-Pk) to a full model that
       additionally includes m, and reports the incremental
       likelihood-ratio (LR) test for m (Supplementary Tables 1-3).
    4. Runs three additional robustness/sensitivity analyses referenced in
       the Discussion (Supplementary Tables 4-6):
           (a) attenuation under adjustment for PC1-PC20,
           (b) robustness to adjustment for clinical molecular subtype,
           (c) bootstrap 95% CI for the change in Harrell's C-index,
           (d) five-fold cross-validated OUT-OF-SAMPLE partial
               log-likelihood and C-index (a survival-model performance
               check, NOT a molecular-subtype classifier), and
           (e) a Schoenfeld residual test of the proportional-hazards
               assumption for m.
    5. Performs a built-in reproducibility check: every computed quantity
       above is compared against the value reported in the revised
       manuscript, and a PASS/REVIEW table is printed at the end.

This script does NOT include: QUBO/Hamiltonian construction or solving,
GMM/BIC subtype clustering, w1-w2 parameter-sweep or landscape figures, the
metabolic frustration index, or any Figure-3-specific analysis. Those are
covered by separate Supplementary Code items (Code 1: main analysis and
figure-generation pipeline; the Figure 3 reproduction package; the
null-model and sensitivity-analysis packages for the Discussion's H0-A-D
controls). Keeping this script narrowly scoped to the order parameter and
its Supplementary Table 1-6 survival benchmarks is intentional and mirrors
the Legend below.

--------------------------------------------------------------------------
IMPORTANT INTERPRETIVE CAVEAT AND TERMINOLOGY NOTE
--------------------------------------------------------------------------
"m" is termed the model-derived order parameter, matching its description
in the Abstract and Results ("A model-derived order parameter is
significantly associated with overall and disease-free survival..."). It
is constructed from the sign of each reaction's standard transformed Gibbs
free energy (sigma_r = sign(-dG_r)), so the quantity is thermodynamically
motivated even though it is not labeled "thermodynamic" in running prose.

This script does NOT, by itself, establish that the survival
association reported below (Tables 1-6) is specific to that thermodynamic
(dG-based) sign convention, as opposed to any similarly constructed
balanced combination of the 235 correlated reaction z-scores. That
question is addressed separately by the H0-B permutation analysis
(see the accompanying null-model-analysis package), which found the real
sigma_r assignment's Cox coefficient for m indistinguishable from three
permutation-null distributions (full permutation, empirical p = 0.524;
pathway-block-flip, p = 0.258; within-pathway shuffle, p = 0.907). The
manuscript's own Discussion therefore interprets m's prognostic value as
arising from the coordinated, network-constrained structure of the
underlying reaction set rather than from a validated reaction-level
thermodynamic directionality (see "Interpreting m as a network-coherent
composite biomarker" in the Discussion). Readers of this script alone
should not conclude, from the Tables 1-6 results here, that m's prognostic
value has been shown to be thermodynamically specific -- that specific
claim is the one thing this script cannot and does not test. Retain the
name "order parameter m" (dropping the qualifier "thermodynamic" in
running prose where appropriate) unless this caveat is repeated alongside
it.

--------------------------------------------------------------------------
Environment used to generate the numbers reported in the manuscript
--------------------------------------------------------------------------
Python      3.10.13
numpy       1.26.4
pandas      2.2.2
scipy       1.13.0
scikit-learn 1.4.2
lifelines   0.28.0

Install with:
    pip install numpy==1.26.4 pandas==2.2.2 scipy==1.13.0 \
                scikit-learn==1.4.2 lifelines==0.28.0

--------------------------------------------------------------------------
Required input files (place in a single directory, pass as --data-dir)
--------------------------------------------------------------------------
    Supplementary_Data_1_Human_reactions_235_with_deltaG_equation.csv
        columns: Reaction, dG
    Supplementary_Data_2_Y497_reaction235_patient_matrix_mean.csv
        rows = 235 reactions, columns = 497 patients, values = mean
        reaction activity (already reaction-mapped from gene expression)
    y497_clinic.txt   (tab-separated)
        columns required: Sample, Age, Gender, AJCC, TYPE, death,
        durationOS, Recurrent, duration of disease-free survival

--------------------------------------------------------------------------
Usage
--------------------------------------------------------------------------
    python Supplementary_Code_2_order_parameter_and_survival_benchmarks.py \\
        --data-dir /path/to/supplementary_data \\
        --out-dir  /path/to/output

Outputs (written to --out-dir):
    Supplementary_Table_1_Cox_OS_PC5.csv
    Supplementary_Table_2_Cox_DFS_PC5.csv
    Supplementary_Table_3_Cox_PC10_sensitivity.csv
    Supplementary_Table_4_Cox_PC20_and_subtype_sensitivity.csv
    Supplementary_Table_5_bootstrap_Cindex_and_5fold_CV.csv
    Supplementary_Table_6_Schoenfeld_residual_test.csv
    patient_order_parameter_m.csv
    reproducibility_check_report.txt

NOTE ON TABLE NUMBERING: Supplementary Table numbering follows the mapping
used internally during preparation of this script (Table 1 = OS/PC1-5,
Table 2 = DFS/PC1-5, Table 3 = PC1-10 sensitivity, Table 4 = PC1-20 and
molecular-subtype sensitivity, Table 5 = bootstrap/cross-validation, Table 6
= Schoenfeld test). Confirm this mapping against the final Supplementary
Table numbers used in the typeset manuscript before submission, and update
the file names / in-text citations if they differ.
"""

from __future__ import annotations

import argparse
import os
import sys
import warnings
from dataclasses import dataclass, field
from typing import Dict, List, Optional, Sequence, Tuple

import numpy as np
import pandas as pd
from scipy import stats
from sklearn.decomposition import PCA
from sklearn.model_selection import KFold
from lifelines import CoxPHFitter

warnings.filterwarnings("ignore", category=RuntimeWarning)
warnings.filterwarnings("ignore", category=FutureWarning)

DG_FILE = "Supplementary_Data_1_Human_reactions_235_with_deltaG_equation.csv"
EXPR_FILE = "Supplementary_Data_2_Y497_reaction235_patient_matrix_mean.csv"
CLINIC_FILE = "y497_clinic.txt"

OS_TIME_COL = "durationOS"
OS_EVENT_COL = "death"
DFS_TIME_COL = "duration of disease-free survival"
DFS_EVENT_COL = "Recurrent"
AGE_COL = "Age"
SEX_COL = "Gender"
STAGE_COL = "AJCC"
SUBTYPE_COL = "TYPE"

RANDOM_SEED = 0


# =============================================================================
# 1. DATA LOADING AND THE CORRECTED ORDER PARAMETER (Methods, as written)
# =============================================================================

@dataclass
class Dataset:
    m: pd.Series
    pcs: pd.DataFrame          # PC1..PC20 of the 235-reaction, per-reaction z-scored matrix
    explained_variance: np.ndarray
    clinic: pd.DataFrame
    n_reactions: int


def load_and_compute(data_dir: str, n_pcs: int = 20) -> Dataset:
    dg = pd.read_csv(os.path.join(data_dir, DG_FILE)).set_index("Reaction")["dG"]
    X = pd.read_csv(os.path.join(data_dir, EXPR_FILE), index_col=0)
    clinic = pd.read_csv(os.path.join(data_dir, CLINIC_FILE), sep="\t").set_index("Sample")

    rxn = X.index.intersection(dg.index)
    if len(rxn) < 200:
        raise ValueError(
            f"Only {len(rxn)} reactions found in common between the expression "
            f"matrix and the dG table; expected all 235. Check the input files."
        )
    dg = dg.loc[rxn]
    X = X.loc[rxn]

    common_patients = X.columns.intersection(clinic.index)
    X = X[common_patients]
    clinic = clinic.loc[common_patients].copy()

    # --- corrected order parameter, literal Methods formula ---
    #   sigma_r = sign(-dG_r)
    #   z_ir    = per-reaction z-score across the full patient cohort
    #   m_i     = (1/235) * sum_r sigma_r * z_ir
    sigma = np.sign(-dg.values)
    Z = X.sub(X.mean(axis=1), axis=0).div(X.std(axis=1), axis=0)
    m = pd.Series((sigma[:, None] * Z.values).mean(axis=0), index=X.columns, name="m")

    # --- transcriptomic principal components of the same 235-reaction matrix ---
    pca = PCA(n_components=n_pcs, random_state=RANDOM_SEED)
    pc_scores = pca.fit_transform(Z.T.values)
    pcs = pd.DataFrame(pc_scores, index=Z.columns, columns=[f"PC{i+1}" for i in range(n_pcs)])

    clinic = clinic.dropna(subset=[STAGE_COL])
    common = m.index.intersection(pcs.index).intersection(clinic.index)
    m = m.loc[common]
    pcs = pcs.loc[common]
    clinic = clinic.loc[common]

    clinic["death"] = clinic[OS_EVENT_COL].astype(float)
    clinic[DFS_EVENT_COL] = clinic[DFS_EVENT_COL].astype(float)
    clinic["Gender_male"] = (clinic[SEX_COL].astype(str).str.lower() == "male").astype(float)
    stage_dummies = pd.get_dummies(clinic[STAGE_COL], prefix="stage", drop_first=True).astype(float)
    subtype_dummies = pd.get_dummies(clinic[SUBTYPE_COL], prefix="type", drop_first=True).astype(float)
    clinic = pd.concat([clinic, stage_dummies, subtype_dummies], axis=1)

    return Dataset(
        m=m,
        pcs=pcs,
        explained_variance=pca.explained_variance_ratio_,
        clinic=clinic,
        n_reactions=len(rxn),
    )


# =============================================================================
# 2. VARIANCE OF m EXPLAINED BY TRANSCRIPTOMIC PCs
# =============================================================================

def variance_of_m_explained_by_pcs(m: pd.Series, pcs: pd.DataFrame, k: int) -> float:
    """R^2 of an OLS regression of m on PC1..PCk (matches the manuscript's
    'PC1-PC5 / PC1-PC10 explained X% of the variance in m')."""
    cols = [f"PC{i+1}" for i in range(k)]
    X = pcs[cols].values
    X = np.column_stack([np.ones(len(X)), X])
    beta, *_ = np.linalg.lstsq(X, m.values, rcond=None)
    fitted = X @ beta
    ss_res = np.sum((m.values - fitted) ** 2)
    ss_tot = np.sum((m.values - m.values.mean()) ** 2)
    return 1.0 - ss_res / ss_tot


# =============================================================================
# 3. COX MODELS AND LIKELIHOOD-RATIO TESTS
# =============================================================================

def clinical_covariates(clinic: pd.DataFrame) -> List[str]:
    stage_cols = [c for c in clinic.columns if c.startswith("stage_")]
    return [AGE_COL, "Gender_male"] + stage_cols


def fit_cox(df: pd.DataFrame, cols: Sequence[str], time_col: str, event_col: str) -> CoxPHFitter:
    cph = CoxPHFitter()
    data = df[[time_col, event_col] + list(cols)].dropna()
    cph.fit(data, duration_col=time_col, event_col=event_col)
    return cph


def lr_test(reduced: CoxPHFitter, full: CoxPHFitter, df: int = 1) -> Tuple[float, float]:
    lr_stat = 2.0 * (full.log_likelihood_ - reduced.log_likelihood_)
    p_value = stats.chi2.sf(lr_stat, df=df)
    return lr_stat, p_value


def hr_ci_p(cph: CoxPHFitter, name: str, per_sd: Optional[float] = None) -> Tuple[float, float, float, float]:
    coef = cph.summary.loc[name, "coef"]
    lo = cph.summary.loc[name, "coef lower 95%"]
    hi = cph.summary.loc[name, "coef upper 95%"]
    p = cph.summary.loc[name, "p"]
    if per_sd is not None:
        coef, lo, hi = coef * per_sd, lo * per_sd, hi * per_sd
    return np.exp(coef), np.exp(lo), np.exp(hi), p


def pc_adjusted_cox_benchmark(
    ds: Dataset, outcome: str, n_pcs: int, extra_covariates: Optional[List[str]] = None
) -> Dict[str, float]:
    """Reduced model: clinical + PC1..PCk (+ extra_covariates).
    Full model:    reduced + m.
    Returns HR/CI/p for m in the full model, plus the LR test for adding m."""
    if outcome.upper() == "OS":
        time_col, event_col = OS_TIME_COL, OS_EVENT_COL
    elif outcome.upper() == "DFS":
        time_col, event_col = DFS_TIME_COL, DFS_EVENT_COL
    else:
        raise ValueError("outcome must be 'OS' or 'DFS'")

    df = ds.clinic.join(ds.pcs).join(ds.m)
    pc_cols = [f"PC{i+1}" for i in range(n_pcs)]
    base_cols = clinical_covariates(ds.clinic) + pc_cols + (extra_covariates or [])

    reduced = fit_cox(df, base_cols, time_col, event_col)
    full = fit_cox(df, base_cols + ["m"], time_col, event_col)
    lr_stat, lr_p = lr_test(reduced, full)
    hr, lo, hi, p = hr_ci_p(full, "m")

    return {
        "outcome": outcome,
        "n_pcs": n_pcs,
        "n": len(df.dropna(subset=base_cols + ["m", time_col, event_col])),
        "HR_m": hr, "CI_low": lo, "CI_high": hi, "p_m": p,
        "LR_stat": lr_stat, "LR_p": lr_p,
        "reduced_loglik": reduced.log_likelihood_, "full_loglik": full.log_likelihood_,
    }


# =============================================================================
# 4. ADDITIONAL ROBUSTNESS / SENSITIVITY ANALYSES (Supplementary Tables 4-6)
# =============================================================================

def bootstrap_delta_cindex(
    ds: Dataset, n_boot: int = 1000, seed: int = RANDOM_SEED
) -> Tuple[float, float, float]:
    df = ds.clinic.join(ds.pcs[[f"PC{i+1}" for i in range(5)]]).join(ds.m).reset_index(drop=True)
    base_cols = clinical_covariates(ds.clinic) + [f"PC{i+1}" for i in range(5)]
    rng = np.random.default_rng(seed)
    deltas = []
    idx_all = df.index.to_numpy()
    for _ in range(n_boot):
        boot_idx = rng.choice(idx_all, size=len(df), replace=True)
        dfb = df.loc[boot_idx].reset_index(drop=True)
        try:
            ci_base = fit_cox(dfb, base_cols, OS_TIME_COL, OS_EVENT_COL).concordance_index_
            ci_full = fit_cox(dfb, base_cols + ["m"], OS_TIME_COL, OS_EVENT_COL).concordance_index_
            deltas.append(ci_full - ci_base)
        except Exception:
            continue
    deltas = np.array(deltas)
    lo, hi = np.percentile(deltas, [2.5, 97.5])
    return float(deltas.mean()), float(lo), float(hi)


def five_fold_cv_performance(
    ds: Dataset, n_splits: int = 5, seed: int = RANDOM_SEED
) -> Dict[str, float]:
    df = ds.clinic.join(ds.pcs[[f"PC{i+1}" for i in range(5)]]).join(ds.m).reset_index(drop=True)
    base_cols = clinical_covariates(ds.clinic) + [f"PC{i+1}" for i in range(5)]
    kf = KFold(n_splits=n_splits, shuffle=True, random_state=seed)

    def cv_scores(cols):
        scores = []
        for train_idx, test_idx in kf.split(df):
            train, test = df.iloc[train_idx], df.iloc[test_idx]
            cph = fit_cox(train, cols, OS_TIME_COL, OS_EVENT_COL)
            pll = cph.score(test[[OS_TIME_COL, OS_EVENT_COL] + cols], scoring_method="log_likelihood")
            ci = cph.score(test[[OS_TIME_COL, OS_EVENT_COL] + cols], scoring_method="concordance_index")
            scores.append((pll, ci))
        return np.array(scores)

    base_scores = cv_scores(base_cols)
    full_scores = cv_scores(base_cols + ["m"])
    return {
        "baseline_partial_loglik": base_scores[:, 0].mean(),
        "full_partial_loglik": full_scores[:, 0].mean(),
        "baseline_Cindex": base_scores[:, 1].mean(),
        "full_Cindex": full_scores[:, 1].mean(),
    }


def schoenfeld_test_for_m(ds: Dataset, transform: str = "km") -> float:
    """Schoenfeld-residual proportional-hazards test for m, adjusted for the
    same clinical + PC1-5 covariates used in the main Cox model.

    lifelines' `check_assumptions` prints, for each covariate, two rows (one
    per KM- and rank-transformed time; column order is
    ``test_statistic, p, -log2(p)``). This function parses the row for
    covariate 'm' under the requested `transform` ('km' or 'rank') and
    returns its p-value. Values printed as "<0.005" are returned as 0.005
    (a conservative /upper/ bound on the true p-value) with a note printed
    to stderr; if the reported reference p-value is not this small, that is
    fine as this only affects very significant, non-borderline results.
    """
    df = ds.clinic.join(ds.pcs[[f"PC{i+1}" for i in range(5)]]).join(ds.m)
    base_cols = clinical_covariates(ds.clinic) + [f"PC{i+1}" for i in range(5)]
    cph = fit_cox(df, base_cols + ["m"], OS_TIME_COL, OS_EVENT_COL)

    import io
    buf = io.StringIO()
    old_stdout = sys.stdout
    sys.stdout = buf
    try:
        cph.check_assumptions(
            df[[OS_TIME_COL, OS_EVENT_COL, "m"] + base_cols], p_value_threshold=0.05, show_plots=False
        )
    except Exception:
        pass
    sys.stdout = old_stdout
    output = buf.getvalue()

    p_m = np.nan
    current_covariate = None
    for line in output.splitlines():
        if not line.strip():
            continue
        starts_new_covariate = not line.startswith(" ")
        parts = line.split()
        if starts_new_covariate and len(parts) >= 4 and parts[1] in ("km", "rank"):
            current_covariate, row_transform, p_str = parts[0], parts[1], parts[-2]
        elif (not starts_new_covariate) and len(parts) >= 3 and parts[0] in ("km", "rank"):
            row_transform, p_str = parts[0], parts[-2]
        else:
            continue
        if current_covariate == "m" and row_transform == transform:
            p_m = 0.005 if p_str.startswith("<") else float(p_str)
    return p_m


# =============================================================================
# 5. BUILT-IN REPRODUCIBILITY CHECK
# =============================================================================

REFERENCE = {
    "var_explained_PC5_in_m": 0.379,
    "var_explained_PC10_in_m": 0.400,
    "OS_PC5_HR_m": 4.44, "OS_PC5_LR_p": 0.010,
    "DFS_PC5_HR_m": 4.01, "DFS_PC5_LR_p": 0.011,
    "OS_PC10_LR_p": 0.007,
    "DFS_PC10_LR_p": 0.010,
    "OS_PC20_HR_m_per_sd": 1.54, "OS_PC20_p": 0.083,
    "DFS_PC20_HR_m_per_sd": 1.38, "DFS_PC20_p": 0.175,
    "OS_subtype_HR_m_per_sd": 1.40, "OS_subtype_p": 0.006,
    "DFS_subtype_HR_m_per_sd": 1.38, "DFS_subtype_p": 0.005,
    "bootstrap_delta_Cindex": 0.010,
    "bootstrap_CI_low": 0.001, "bootstrap_CI_high": 0.025,
    "cv_baseline_loglik": -1.854, "cv_full_loglik": -1.839,
    "cv_baseline_Cindex": 0.684, "cv_full_Cindex": 0.691,
    "schoenfeld_p_m": 0.53,
}


def check(report: List[str], name: str, computed: float, reference: float,
          tol: float = 0.05, tol_type: str = "abs") -> None:
    if np.isnan(computed):
        ok = False
    elif tol_type == "abs":
        ok = abs(computed - reference) <= tol
    elif tol_type == "sig":
        ok = (computed < 0.05) == (reference < 0.05)
    else:
        raise ValueError(tol_type)
    status = "PASS" if ok else "REVIEW"
    report.append(f"[{status}] {name}: computed={computed:.4g}, reference={reference:.4g}")


# =============================================================================
# 6. MAIN
# =============================================================================

def main() -> None:
    parser = argparse.ArgumentParser(description=__doc__, formatter_class=argparse.RawDescriptionHelpFormatter)
    parser.add_argument("--data-dir", default=".")
    parser.add_argument("--out-dir", default="./code2_output")
    parser.add_argument("--n-boot", type=int, default=1000)
    args = parser.parse_args()

    os.makedirs(args.out_dir, exist_ok=True)
    report: List[str] = []

    print("Loading data and computing the corrected order parameter m ...")
    ds = load_and_compute(args.data_dir, n_pcs=20)
    ds.m.to_frame().to_csv(os.path.join(args.out_dir, "patient_order_parameter_m.csv"))
    print(f"  n = {len(ds.m)} patients, {ds.n_reactions} reactions, "
          f"SD(m) = {ds.m.std():.4f}\n")

    # --- variance of m explained by PCs ---
    var5 = variance_of_m_explained_by_pcs(ds.m, ds.pcs, 5)
    var10 = variance_of_m_explained_by_pcs(ds.m, ds.pcs, 10)
    print(f"Variance of m explained: PC1-5 = {var5:.3f}, PC1-10 = {var10:.3f}")
    check(report, "Variance of m explained by PC1-5", var5, REFERENCE["var_explained_PC5_in_m"], tol=0.05)
    check(report, "Variance of m explained by PC1-10", var10, REFERENCE["var_explained_PC10_in_m"], tol=0.05)

    # --- Supplementary Table 1: OS, Clinical + PC1-5 (+/- m) ---
    t1 = pc_adjusted_cox_benchmark(ds, "OS", 5)
    pd.DataFrame([t1]).to_csv(os.path.join(args.out_dir, "Supplementary_Table_1_Cox_OS_PC5.csv"), index=False)
    check(report, "OS (PC1-5) HR for m", t1["HR_m"], REFERENCE["OS_PC5_HR_m"], tol=0.5)
    check(report, "OS (PC1-5) LR p for m", t1["LR_p"], REFERENCE["OS_PC5_LR_p"], tol=0.02)

    # --- Supplementary Table 2: DFS, Clinical + PC1-5 (+/- m) ---
    t2 = pc_adjusted_cox_benchmark(ds, "DFS", 5)
    pd.DataFrame([t2]).to_csv(os.path.join(args.out_dir, "Supplementary_Table_2_Cox_DFS_PC5.csv"), index=False)
    check(report, "DFS (PC1-5) HR for m", t2["HR_m"], REFERENCE["DFS_PC5_HR_m"], tol=0.5)
    check(report, "DFS (PC1-5) LR p for m", t2["LR_p"], REFERENCE["DFS_PC5_LR_p"], tol=0.02)

    # --- Supplementary Table 3: PC1-10 sensitivity, OS and DFS ---
    t3_os = pc_adjusted_cox_benchmark(ds, "OS", 10)
    t3_dfs = pc_adjusted_cox_benchmark(ds, "DFS", 10)
    pd.DataFrame([t3_os, t3_dfs]).to_csv(
        os.path.join(args.out_dir, "Supplementary_Table_3_Cox_PC10_sensitivity.csv"), index=False
    )
    check(report, "OS (PC1-10) LR p for m", t3_os["LR_p"], REFERENCE["OS_PC10_LR_p"], tol=0.02)
    check(report, "DFS (PC1-10) LR p for m", t3_dfs["LR_p"], REFERENCE["DFS_PC10_LR_p"], tol=0.02)

    # --- Supplementary Table 4: PC1-20 attenuation + molecular-subtype-adjusted ---
    m_sd = ds.m.std()
    t4_pc20_os = pc_adjusted_cox_benchmark(ds, "OS", 20)
    t4_pc20_dfs = pc_adjusted_cox_benchmark(ds, "DFS", 20)
    type_cols = [c for c in ds.clinic.columns if c.startswith("type_")]
    t4_subtype_os = pc_adjusted_cox_benchmark(ds, "OS", 5, extra_covariates=type_cols)
    t4_subtype_dfs = pc_adjusted_cox_benchmark(ds, "DFS", 5, extra_covariates=type_cols)

    def per_sd(row):
        r = dict(row)
        beta = np.log(row["HR_m"])
        r["HR_m_per_SD"] = np.exp(beta * m_sd)
        return r

    t4_rows = [per_sd(t4_pc20_os), per_sd(t4_pc20_dfs), per_sd(t4_subtype_os), per_sd(t4_subtype_dfs)]
    for row, label in zip(t4_rows, ["PC20_OS", "PC20_DFS", "subtype_OS", "subtype_DFS"]):
        row["model"] = label
    pd.DataFrame(t4_rows).to_csv(
        os.path.join(args.out_dir, "Supplementary_Table_4_Cox_PC20_and_subtype_sensitivity.csv"), index=False
    )
    check(report, "OS (PC1-20) HR_m per-SD", t4_rows[0]["HR_m_per_SD"], REFERENCE["OS_PC20_HR_m_per_sd"], tol=0.15)
    check(report, "OS (PC1-20) p for m", t4_rows[0]["p_m"], REFERENCE["OS_PC20_p"], tol=0.05)
    check(report, "DFS (PC1-20) HR_m per-SD", t4_rows[1]["HR_m_per_SD"], REFERENCE["DFS_PC20_HR_m_per_sd"], tol=0.15)
    check(report, "DFS (PC1-20) p for m", t4_rows[1]["p_m"], REFERENCE["DFS_PC20_p"], tol=0.05)
    check(report, "OS (subtype-adj) HR_m per-SD", t4_rows[2]["HR_m_per_SD"], REFERENCE["OS_subtype_HR_m_per_sd"], tol=0.15)
    check(report, "OS (subtype-adj) p for m", t4_rows[2]["p_m"], REFERENCE["OS_subtype_p"], tol=0.02)
    check(report, "DFS (subtype-adj) HR_m per-SD", t4_rows[3]["HR_m_per_SD"], REFERENCE["DFS_subtype_HR_m_per_sd"], tol=0.15)
    check(report, "DFS (subtype-adj) p for m", t4_rows[3]["p_m"], REFERENCE["DFS_subtype_p"], tol=0.02)

    # --- Supplementary Table 5: bootstrap Delta C-index + 5-fold CV ---
    print("\nRunning bootstrap (this may take a moment) ...")
    boot_mean, boot_lo, boot_hi = bootstrap_delta_cindex(ds, n_boot=args.n_boot)
    cv = five_fold_cv_performance(ds)
    t5 = pd.DataFrame([{
        "bootstrap_mean_delta_Cindex": boot_mean,
        "bootstrap_CI_low": boot_lo, "bootstrap_CI_high": boot_hi,
        **cv,
    }])
    t5.to_csv(os.path.join(args.out_dir, "Supplementary_Table_5_bootstrap_Cindex_and_5fold_CV.csv"), index=False)
    check(report, "Bootstrap mean Delta C-index", boot_mean, REFERENCE["bootstrap_delta_Cindex"], tol=0.01)
    check(report, "Bootstrap CI excludes zero (lower bound > 0)", boot_lo, 0.0, tol=0.0, tol_type="sig")
    check(report, "5-fold CV baseline partial log-lik", cv["baseline_partial_loglik"], REFERENCE["cv_baseline_loglik"], tol=0.05)
    check(report, "5-fold CV full partial log-lik", cv["full_partial_loglik"], REFERENCE["cv_full_loglik"], tol=0.05)
    check(report, "5-fold CV baseline C-index", cv["baseline_Cindex"], REFERENCE["cv_baseline_Cindex"], tol=0.02)
    check(report, "5-fold CV full C-index", cv["full_Cindex"], REFERENCE["cv_full_Cindex"], tol=0.02)

    # --- Supplementary Table 6: Schoenfeld residual test ---
    schoenfeld_p = schoenfeld_test_for_m(ds)
    pd.DataFrame([{"covariate": "m", "schoenfeld_p": schoenfeld_p}]).to_csv(
        os.path.join(args.out_dir, "Supplementary_Table_6_Schoenfeld_residual_test.csv"), index=False
    )
    check(report, "Schoenfeld test p for m (no PH violation)", schoenfeld_p, REFERENCE["schoenfeld_p_m"], tol=0.15)

    # --- write reproducibility report ---
    report_path = os.path.join(args.out_dir, "reproducibility_check_report.txt")
    with open(report_path, "w") as f:
        f.write("Supplementary Code 2 - reproducibility check\n")
        f.write("=" * 70 + "\n")
        for line in report:
            f.write(line + "\n")
        n_pass = sum(1 for line in report if line.startswith("[PASS]"))
        f.write("=" * 70 + "\n")
        f.write(f"SUMMARY: {n_pass}/{len(report)} checks within tolerance.\n")

    print("\n" + "\n".join(report))
    print(f"\nWrote Supplementary Tables 1-6 and reproducibility report to: {args.out_dir}")


if __name__ == "__main__":
    main()
```
